# Nucleotide-binding motifs nucleated folding of the first enzymes

**DOI:** 10.64898/2026.08.08.743454

**Authors:** Koh Seya, Tatsuya Corlett, Hamza Giaffar, Andre Lecona Buttelli, Claudèle Lemay-St-Denis, Kosuke Fujishima, Shina Caroline Lynn Kamerlin, Rachel Kolodny, Eric Smith, Liam M. Longo

**Author notes:** Equal contribution (Koh Seya, Tatsuya Corlett, Hamza Giaffar).

## Abstract

The earliest stages of protein evolution remain a mystery: the nature of the first protein forms, their roles in emergent biological systems, and the forces that shaped them are largely unknown. Here, we combine insights from metabolic modelling, the organization of protein structure space, and protein folding mechanisms to probe the emergence of two ubiquitous cofactor-binding folds: Rossmanns and P-loop NTPases. While both folds are essential for contemporary life, we show that Rossmanns catalyze reactions deeper within the metabolic core and are more central in structure space than P-loop NTPases. Folding mechanism analysis further reveals that, whereas P-loop NTPases may require non-local interactions to fold, Rossmann folding can be nucleated by a structural module at the heart of the fold that contains a nucleotide-binding motif. Because this motif also directly mediates biochemical activity, this result suggests how early proteins may have compactly satisfied both folding and biochemical activity. Our results imply that folding constraints favored early enzymatic forms with compact binding motifs and modest catalytic roles. We conclude that the early emergence of the Rossmann fold reflects the chemical and physical constraints of protein folding, explaining both its profound antiquity and sustained longevity.

## Introduction

Folded biopolymers mediate virtually all cellular processes, and the special properties of folded forms—manifest in their capacity for specific catalysis and molecular recognition—are necessary for the realization of complex life. Chief among the folded biopolymers are proteins, which, through the adoption of a diverse repertoire of structures, have become the primary biological catalysts. The earliest stages of protein evolution, however, are poorly resolved. It remains unclear what the earliest protein forms were, what roles they played in nascent biological systems, and to what extent they were shaped by contingency or constrained by chemical and physical law. Addressing these questions is essential to understanding both the organization of the known protein universe and the origin of life.

Protein origins questions are difficult to answer because the comparative approaches that underpin almost all protein reconstructions carry inherent inferential limits. Cross-species comparisons converge approximately on the last universal common ancestor (LUCA) [55], while within-fold comparisons converge on fully articulated domains. Highly conserved motifs and ancient events of fragment-sharing between folds have been carefully enumerated [10, 22, 29], yet the interpretation of these patterns for the fold emergence process remains unclear, particularly for the most ancient folds [7]. Most importantly, comparative approaches are largely silent on causation: they do little to separate the often-intertwined contributions of changing physical folding conditions across time and biological selection and refunctionalization. This difficulty is compounded by the fact that the biological and environmental context in which early fold emergence occurred is underspecified and contested.

These causal questions are sharpest for the most ancient fold lineages, which would have emerged when biological refinement of folding conditions was most primitive and limited—for example, generating destabilizing substitutions from more error-prone replication and translation and not yet having evolved the support of chaperones—while their record has also been impacted by selection for the longest time. Further confounding interpretation, owing to their great age, these lineages have had the most opportunity for recruitment into core cellular processes and functional diversification. The Rossmann [44] and P-loop NTPase folds [54] are emblematic of these challenges [29]: both mediate essential functions without which contemporary life would be impossible, and both have diversified extensively [25, 33]. Here, using these two fold lineages as examples, we show that by integrating multiple lines of evidence—the organization of metabolism [14, 12, 5], the organization of structure space [4, 36, 56], the relative topological complexity among folds [41], and the mechanisms of protein folding [52, 53, 35, 38]—we can constrain and explain key aspects of the fold-originating process.

We find that while more P-loop NTPase domains are required by most organisms, the Rossmann fold is more closely associated with the deepest core of metabolism, more central in structure space, and adopts a simpler, more accessible topology. Against this backdrop, models of protein folding reveal that the functional core of various Rossmann domains, which harbors a nucleotide-binding motif, can be a key participant in the folding process itself. This observation demonstrates how core requirements of foldability and activity could have been jointly realized by nascent cofactor-utilizing enzymes within a single, compact structural motif. We hypothesize that the constraint of foldability, which favors a short, tight binding loop, then subsequently enabled Rossmann functional diversification because the binding surface imparts fewer constraints on its cognate ligands. Taken together, our results imply that the first folds to support metabolism did so through simple interactions yielding modest functional contributions, such as catalytic enhancement by uniform binding [1, 37]. We conclude that the early emergence of the Rossmann fold, and *α*/*β* proteins in general, primarily reflects the chemical and physical constraints of protein folding, explaining both the profound antiquity and sustained longevity of the Rossmann domain and its cofactor-binding core.

## Results and Discussion

### Early Rossmanns diversified cofactor utilization whereas early P-loops diversified fold topology

Rossmanns and P-loops are among the most diversified fold lineages, comprising 333 (99.8 percentile) and 232 (99.6 percentile) families in the ECOD database, respectively. For all prokaryote genomes tested, including endosymbionts, at least 1 Rossmann and 7 P-loop domains were detected. In the largest proteomes, more than 500 domains of each type can be present. While more P-loop than Rossmann domains are found in the smallest genomes, the gap between Rossmann and P-loop domain count narrows, and eventually inverts, for the largest genomes (Figure 1A). Fitting the domain count–genome size data to a power law reveals superlinear scaling of Rossmann domains in both bacteria and archaea and sublinear or nearly linear scaling of P-loop domains in bacteria and archaea, respectively (Figure S17). Thus, genome scaling laws suggest that more P-loop domains are required for a functioning cell but that Rossmanns have more successfully diversified, consistent with their greater number of families. 8 Rossmann and 14 P-loop families are highly conserved, present in the majority of organisms within the majority of phyla (Figure 1B, prokaryote distribution score cutoff of 0.8; see Methods for calculation). The likely presence of multiple families of each fold lineage in the LUCA suggests that both Rossmanns and P-loops underwent major functional diversification at the earliest stages of cellular evolution, an interpretation that is supported by efforts to reconstruct the LUCA proteome (Figure S18).

**Figure 1:**
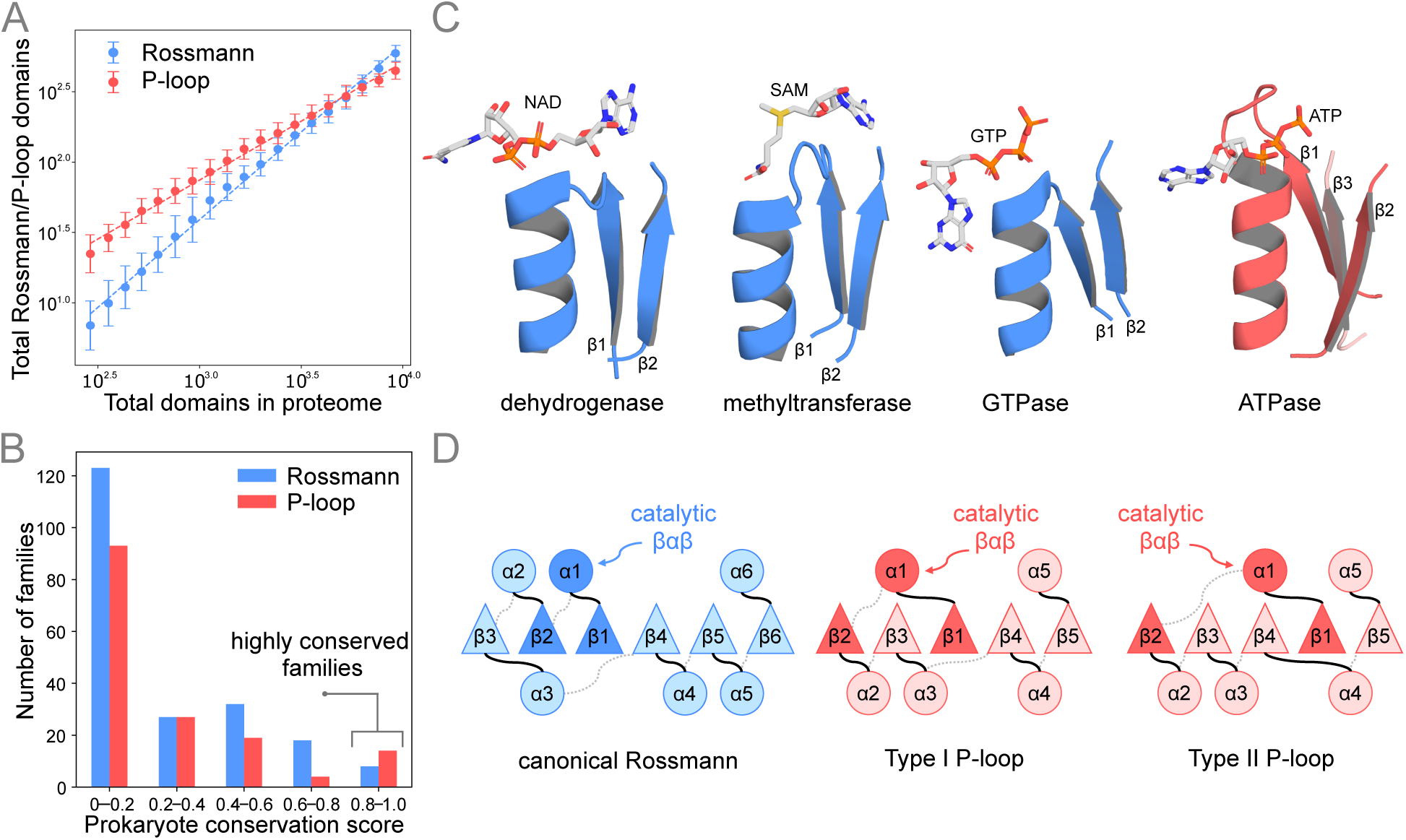
Conservation and diversification of Rossmanns and P-loops. **A.** Proteome-wide scaling of Rossmann and P-loop domains. Number of Rossmann (blue) and P-loop (red) domains plotted against total proteome size across representative prokaryotic genomes. Points show quality score-weighted means ± standard deviations within equal-width proteome size bins (scaling rows); dashed lines are power-law fits. P-loop domains predominate in smaller proteomes, whereas Rossmann domains scale more steeply, diversifying disproportionately in larger proteomes. **B.** Representative structures of nucleotide-binding *βαβ* motifs in Rossmann (blue, left three panels) and P-loop (red, rightmost panel) domains. Rossmann domains accommodate a diverse range of cofactors, including NAD^+^, SAM, and GTP, whereas P-loops selectively bind ATP or GTP via a highly conserved phosphate-binding loop. Representative structures are based on ECOD domains e2nadA1, e3g07A2, e5j2tA1, and e6ln3A1. **C.** Distribution of prokaryote conservation scores for Rossmann (blue) and P-loop (red) evolutionary lineages. Conservation score reflects the proportion of prokaryotic lineages in which each family is detected; families with scores exceeding 0.8 are considered highly conserved. **D.** Topology diagrams of canonical Rossmann and P-loop folds. Schematic secondary structure diagrams of the Rossmann (left), type I P-loop (center), and type II P-loop (right) folds. *β*-strands and *α*-helices are represented as triangles and circles, respectively, numbered by their order in the primary sequence. The *βαβ* motif associated with the conserved nucleotide-binding site in each fold is darkened. Loops on the N-terminal face of the *β*-sheet are shown as solid black lines and loops on the C-terminal face are shown as dashed grey lines.

Treating families with a prokaryote distribution score of greater than 0.80 as putative LUCA families, we examined the nature of early Rossmann and P-loop diversification. Whereas the most conserved Rossmann domains are associated with multiple cofactors—including GTP, SAM, NAD—the most conserved P-loop domains are associated with only ATP and GTP (Figure 1C). For Rossmann domains, early functional diversification through cofactor recruitment was coupled to changes in the structure of the cofactor-binding loop (Figure 1C). For example, cofactor switching between NAD and SAM can be achieved by a 3-residue insertion/deletion [51]. The cofactor-binding loop of P-loops, on the other hand, is highly conserved, adopting a canonical form across all of the most distributed families. Conversely, whereas highly conserved Rossmanns all adopt a common topology with a *β*-strand order of 321456, highly conserved P-loops adopt two distinct topologies (Figure 1D): one in which *β*1 and *β*2 are separated by a single intervening strand (type I, *β*-strand order 23145) and one in which *β*1 and *β*2 are separated by two intervening strands (type II, *β*-strand order 23415). Notably, some P-loop families are characterized by both type I and type II strand topologies, suggesting that topological rearrangement in P-loops is a relatively facile process (Table S2). This observation may be consistent with early efforts at *α*/*β* protein design, which achieved limited control of strand topology [21]. Among the highly conserved P-loop proteins, the two different strand topologies may play distinct roles, with type II P-loops being associated with several large complexes in which conformational changes are coupled to phosphoanhydride hydrolysis (e.g., DEAD helicases, ATP synthase, ABC transporters, and *σ*^54^ transcription factors; Table S2).

### Rossmanns supported early metabolic exploration

Both Rossmanns and P-loops are key contributors to metabolic processes. As a consequence of their ability to utilize diverse cofactors, Rossmanns are associated with an exceptional number of metabolic reactions (1,386, top-ranked fold lineage), far more than P-loops (165, rank 13; Figure 2A). With the exception of translocases, Rossmanns are strongly associated with all of the reaction classes in the Enzyme Commission classification scheme (Figure 2B). P-loops, although more restricted in their reaction scope, are likewise associated with diverse reaction types. Metabolic reactions catalyzed by P-loops are almost exclusively associated with the type I topology (Figure 2C). Although it may be tempting to infer that the greater number of reactions associated with Rossmann domains implies that this fold lineage is more ancient, we note that Rossmanns and P-loops play fundamentally distinct biochemical roles. This fact may invalidate preferential attachment-based arguments, in which more ancient folds would be expected to accumulate more functional associations over time [18]. Moreover, the majority of the P-loop and Rossmann families are not highly conserved (Figure 1B); thus, the metabolic dominance of Rossmanns may reflect recent innovations rather than core metabolic roles.

**Figure 2:**
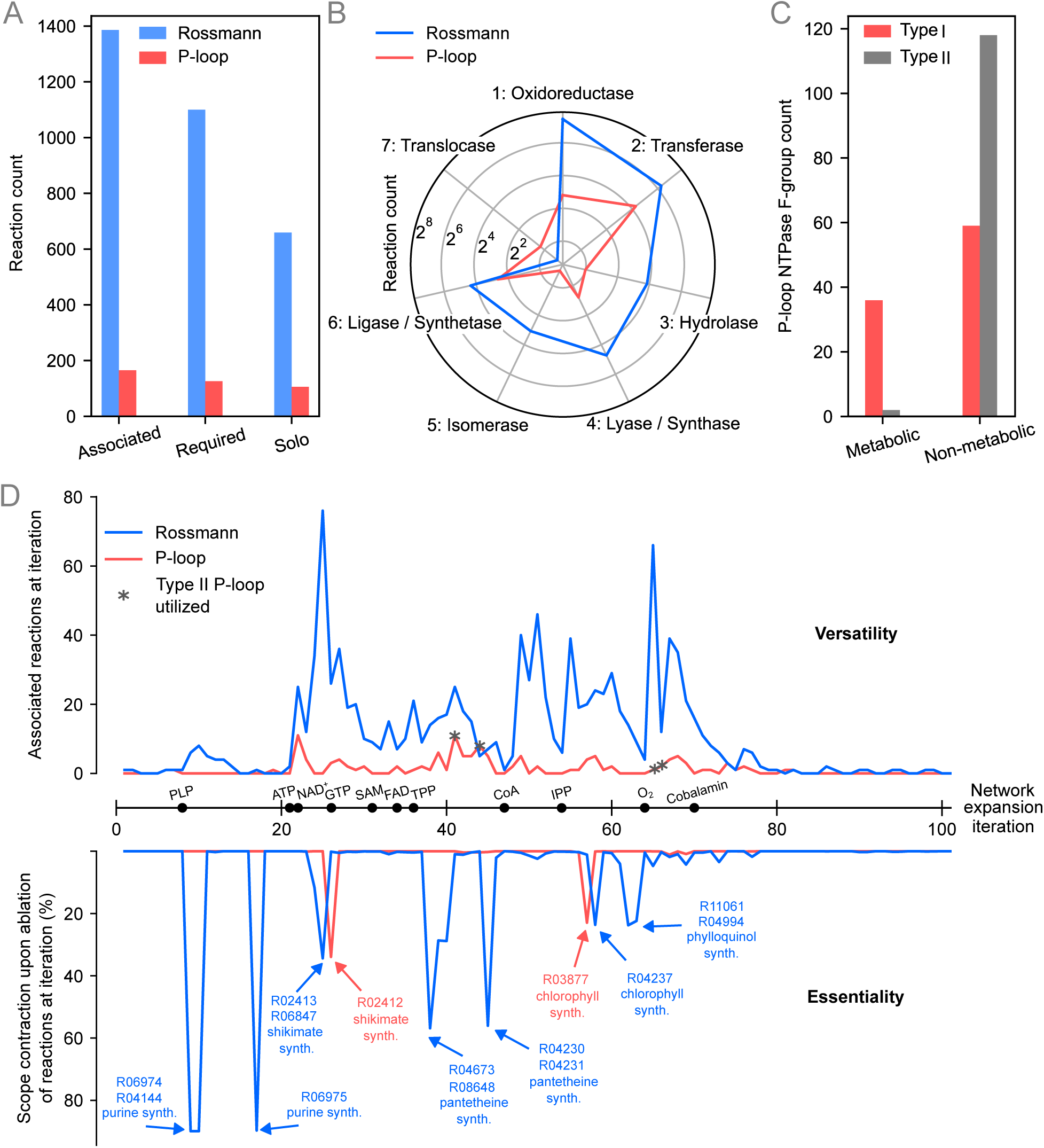
Metabolic roles of Rossmanns and P-loops. **A.** The number of metabolic reactions in network expansion where Rossmann (blue) or P-loop (red) domains exist in one or more enzyme orthogroups associated with a reaction (‘Associated’) or all enzyme orthogroups associated with a reaction (‘Required’). Enzyme orthogroups in which only a Rossmann or P-loop domain is detected are classified as ‘Solo’. **B.** Radar plot comparing EC class distributions between metabolic reactions in network expansion associated with Rossmann (blue) and P-loop (red). Axes represent first-digit EC numbers (1–7), with the radii corresponding to the associated reaction counts within each class. **C.** The number of type I and type II P-loop F-groups that are associated with metabolic reactions in the metabolic model versus those that are not. **D.** The number of reactions discovered (*y*-axis, top panel) at each iteration of network expansion (*x*-axis, with key cofactor discovery indicated by black circles) for which Rossmann (blue) or P-loop (red) domains are associated. The iterations where a type II P-loop domain catalyzes a metabolic reaction are indicated with grey asterisks. The bottom panel shows the extent of scope contraction (*y*-axis, in percent of full scope) upon ablation of reactions that iteration that require a Rossmann or P-loop domain.

We have recently argued that constraint-based reconstructions of metabolic evolution can be used to make inferences about pre-LUCA enzyme emergence and diversification [5]. Using the network expansion approach [14] and an updated model of biosphere-scale metabolic evolution [12] (see Methods for more details), the significance of Rossmann and P-loop domains for early metabolism was analyzed in two ways (Figure 2D): First, we quantified the number of reactions associated with each fold lineage at early stages of metabolic development. Second, by removing reactions that are dependent on Rossmann or P-loop domains, we assessed the essentiality of these fold lineages based on whether metabolic network growth is blocked or whether alternative reaction pathways or enzymes compensate. We find that Rossmann domains are consistently associated with more reactions than P-loops at nearly all stages of metabolic complexity, both early and late in the constraint-based trajectory (Figure 2D, upper). Likewise, removing Rossmann domains from the metabolic model results in seven points at which subsequent network growth is impeded, compared to just two for P-loop domains (Figure 2D, lower). The fact that the two earliest stopping points are induced by removal of Rossmanns, with the third point approximately concurrent with the stopping point induced by a P-loop removal, supports the proposed early participation of Rossmanns in metabolic evolution. Finally, removal of Rossmann yields more significant scope contractions (Figure 2D, lower)—a 90.7% (400/4,315) contraction of the scope compared to a 36.7% (2,731/4,315) scope contraction for complete removal of P-loops. We note that participation of type II P-loop domains occurs in the latter two-thirds of the trajectory, and is not associated with a significant scope contraction.

Taken together, our metabolic analysis suggests that Rossmann domains were early participants in metabolic evolution, with key later roles played by type I P-loops and subsequent participation by type II P-loops. The profound essentiality of Rossmann domains is a direct consequence of their diversified cofactor associations, and group ablation of Rossmann-dependent reactions requiring each of the cofactor classes yields significant scope contraction for thiamine pyrophosphate (TPP, 57%), flavin adenine dinucleotide (FAD, 57%) and nicotinamide adenine dinucleotide (NAD^+^, 51%) (Figure S19).

### Rossmanns are at the center of ***α***/***β*** structure space

The interpretation of centrality in protein space is not inherently temporal. A highly central region could represent either a founder form, to which many other forms attach because they were derived from it, or a point of convergence, toward which independently derived lineages repeatedly evolve. We hypothesize that, for old protein domains in particular, these alternatives are partly collapsed because the properties of robust foldability and stability that would favor a domain’s emergence under primitive conditions would also be expected to characterize convergent lineages that have survived long eras of selection. Emergence under conditions of weaker folding support and less robust heredity is favored by the same properties that make a domain an attractor in structure space: robust foldability, moderate to high stability, and accessibility from diverse sequences. Conversely, an attractor reached repeatedly from multiple evolutionary trajectories is more likely to appear in reconstructions as a source of new fold lineages. Although major evolutionary innovations (e.g., molecular chaperones) may have shifted historical emergence points away from present-day attractors [50, 43], we hypothesize that, at the global scale, centrality nevertheless encodes a coarse pseudotemporal signal that can be evaluated through comparison with independent historical and biophysical evidence.

Various notions of centrality have been considered, with fragment sharing networks being the traditional perspective applied to studying the global structure of protein space [10, 22]. The observation that globally unrelated protein folds share short regions of homology inspired many to interpret these fold-bridging fragments as potential clues into the primitive fold-emergence process [30, 42]. The rationale behind studying fold-bridging fragments is that they may reflect shared origins, provide clues into the biophysically privileged forms, and clarify the degree to which different protein structure classes can be modularly assembled. Recently, learned latent spaces have been shown to accurately recapitulate the global and local structure of protein space [15, 47, 56], with some models providing a measure of protein similarity that is informed by both sequence and structure considerations. Thus, global centrality within a learned latent space can provide an alternative metric for pseudotemporal inference. Using fragment similarity networks (Figure 3A) and latent space analysis (Figure 3B), we analyzed the relative centrality of the Rossmann and P-loop fold lineages within protein space.

**Figure 3:**
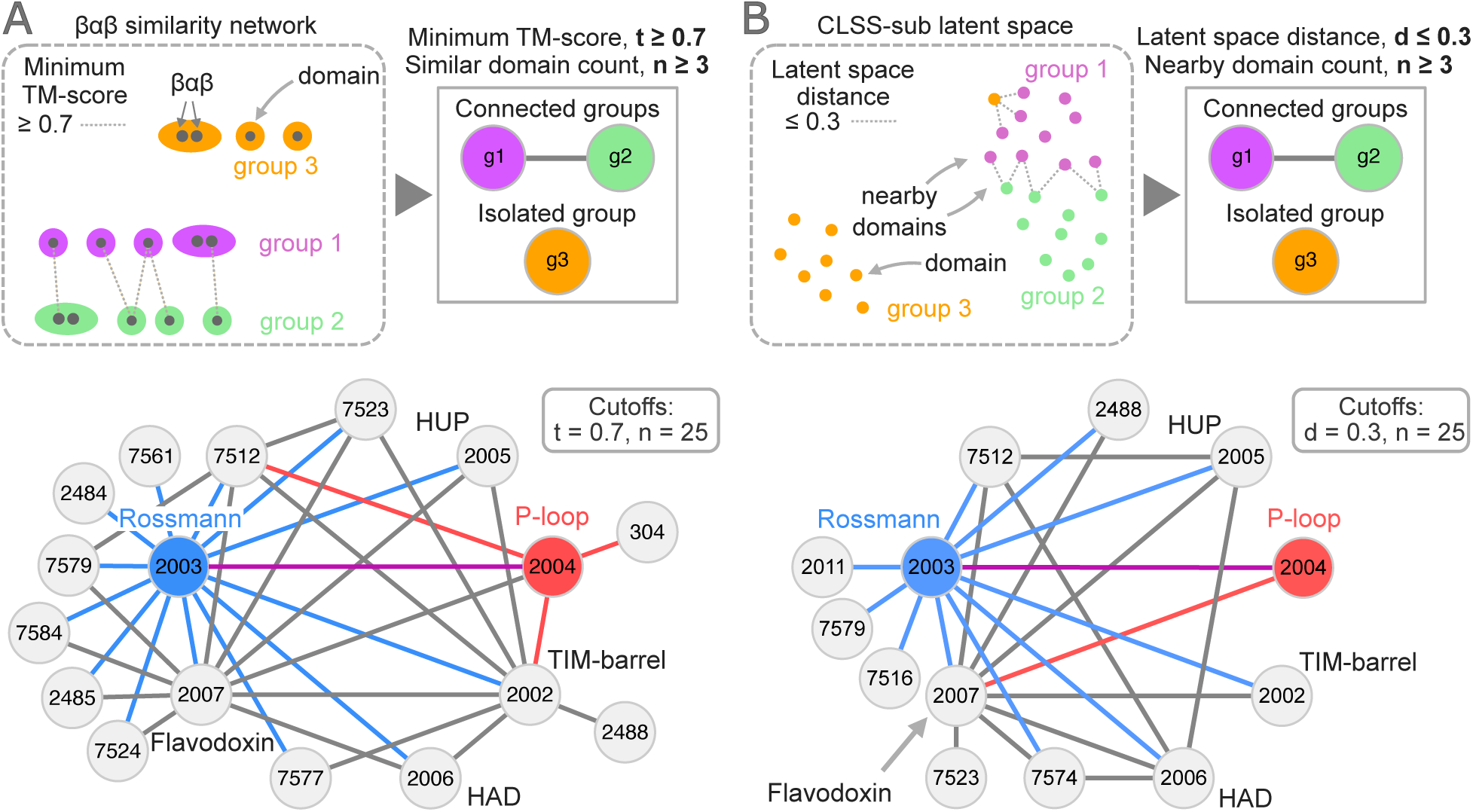
Fold space centrality of Rossmanns and P-loops. **A.** Schematic illustration of the *βαβ* similarity network analysis (top). Each grey dot represents a single *βαβ*. A single domain may contain multiple *βαβ* motifs. *βαβ* motif pairs for which the minimum TM-score is above the cutoff, *t*, are classified as similar. Connections are drawn between two X-groups only if both X-groups have at least *n* domains that connect to the other X-group through a pair of similar *βαβ* motifs. Network constructed using the method shown in the top panel (bottom). The displayed network corresponds to *t* = 0.7 and *n* = 25, and X-group IDs are indicated for each node. **B.** (top) Schematic illustration of the CLSS-sub-based network analysis. Each dot represents a single domain. Domains whose distances in the latent space are below the distance cutoff, *d*, are counted as nearby domains. Connections are drawn between X-group pairs that each contain at least *n* domains nearby the other X-group. (Bottom) Network constructed using the method shown in the top panel. The displayed network corresponds to *d* = 0.3 and *n* = 25, and X-group IDs are indicated for each node.

Proteins from the *α*/*β* class, such as Rossmanns and P-loops, are formed by alternating *α*-helices and *β*-sheets. In three-dimensional space, these secondary structure elements frequently adopt *βαβ* motifs, in which the *β*-strands are part of the central *β*-sheet and the *α*-helices pack against the plane of the *β*-sheet. To generate a fold lineage similarity network, *βαβ* motifs were extracted from the set of representative ECOD domains spanning all known fold lineages (Figure 3A; see Methods for more details). As *α*/*β* proteins comprise both closed *βαβ* motifs, in which the *β*-strands directly hydrogen bond with each other, and open *βαβ* motifs, in which the *β*-strands are part of the same *β*-sheet but are separated by one or more intervening strands, both types of *βαβ* motifs were considered in our analysis. Figure 3A shows the fold-lineage connections based on *βαβ* similarity, requiring a minimum TM-score of 0.7 and a domain count threshold of 25. See Figure S20 for a justification of the minimum TM-score cutoff. This analysis positions Rossmanns (node degree 14) as being fundamentally more connected in protein space than P-loops (node degree 5), and varying the cutoffs does not alter this conclusion (Figure S20). Indeed, Rossmanns are consistently among the most connected fold lineages among the *α*/*β* proteins (Figure S21), with only flavodoxin (X-group 2007) and TIM-barrel (X-group 2002) achieving similar degrees of connectivity (node degrees 11 and 9, respectively, based on a minimum TM-score ≥ 0.7 and domain count ≥ 25). As the Rossmann and P-loop fold lineages are similar in size and occur in proteomes to a similar degree (Figure 1A), the centrality differences observed between them are unlikely to be an artefact of sampling bias. Randomly subsampling (i) 200 or 100 families or (ii) 3,000 or 2,000 *βαβ* motifs from the Rossmann and P-loop fold lineages does not change these results (Figure S22), further supporting this interpretation.

We next turned to machine learning approaches and applied the two recently reported Contrastive Learning Sequence–Structure (CLSS) protein language models [56]. These models were trained by contrastive loss between protein domain (sub)sequences and structures. The two models, CLSS-sub and CLSS-full, differ in that CLSS-sub was trained on a randomly selected subsequence (length 10 residues to the full length of the domain) whereas CLSS-full was trained on full-domain sequences only. Figure 3B shows the fold-lineage connections based on closeness in the full-dimensional CLSS-sub latent space, considering a Euclidean (straight-line) distance cutoff of 0.3 and a domain count cutoff of 25. Note that the latent space of CLSS is normalized such that distance *d* ∈ [0, 2]. See Figure S23 for a justification of these cutoffs, which yield networks that recapitulate several core features of protein space, such as the grouping of the small *β*-barrel folds into a super-fold cluster or metafold. Once again, this analysis positions Rossmanns (node degree 10) as being fundamentally more connected in protein space than P-loops (node degree 2). Varying the cutoffs does not alter this conclusion (Figures S24 and S25). CLSS-full yields effectively the same result (Figure S26), though in this model, fold lineages are significantly more separated in the latent space.

In summary, fragment-based and latent space-based perspectives both highlight Rossmanns as being more central in fold space than P-loops, and indicate that this lineage is one of the most central fold lineages overall. These results are consistent with the more facile emergence of Rossmann forms, and suggest that the structure of fold space implied by a fragment sharing analysis is recapitulated by self-supervised learned representations of protein space.

### The Rossmann topology is more accessible

The most primitive forms of Rossmann and P-loop proteins are unknown. Although Rossmannoid domains can give rise to P-loop domains through *β*-strand swapping (e.g., a 32145 flavodoxin-like domain can yield a canonical 23145 P-loop topology through a *β*-strand swap) (Figure 4A), the directionality of this transition, if it occurred at all, is likewise unclear. The existence of a “type 0” P-loop protein, with no intervening *β*-strand between *β*1 and *β*2, suggests that it is at least possible for P-loop functionality to be encoded on a Rossmannoid chassis (Figure 4B), though this fold is highly derived (prokaryote distribution score = 0.08), and unlikely to represent a primal P-loop form. Nevertheless, an analysis of the strand topologies of these two forms may provide insights into the accessibility of these structures (Figure 4).

**Figure 4:**
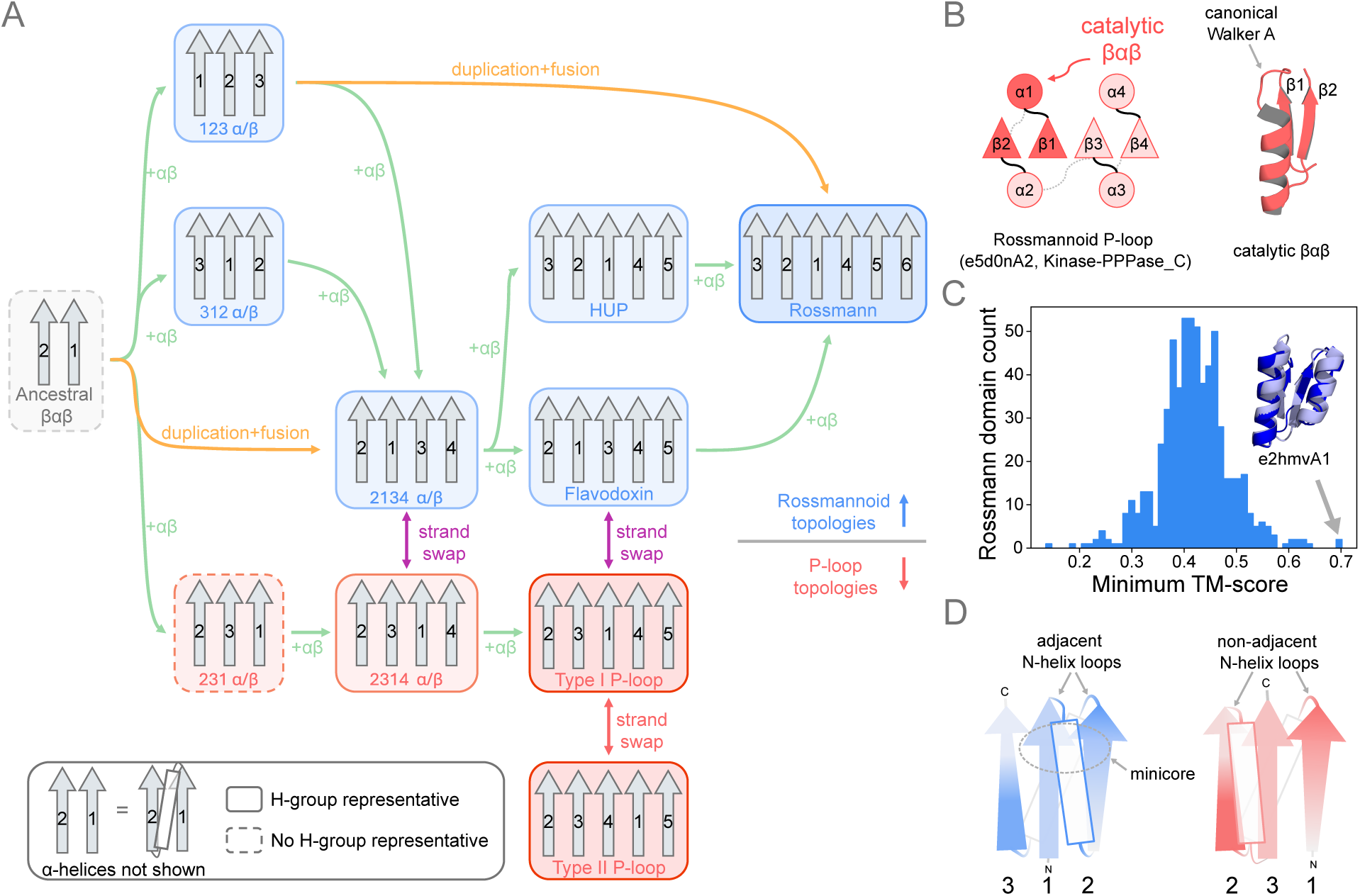
Rossmann and P-loop topological accessibility. **A.** Proposed topological transitions of *α*/*β* proteins from an ancestral *βαβ* motif, considering insertion, duplication, and strand swap events. Rossmannoid topologies are enclosed by blue boxes, whereas P-loop-associated topologies are enclosed by red boxes. To emphasize topological relationships, *α*-helices are omitted. Solid outlines indicate topologies that were observed in a survey of representative ECOD domains, whereas dashed outlines indicate that no corresponding structures were observed. **B.** Topology and structure of a Rossmannoid P-loop (e5d0nA2), which has a strand order of 2134. The dark red region indicates the Walker A-containing *βαβ* motif. **C.** The minimum TM-score distribution between the first and second winds of canonical Rossmann domains with an inset structure alignment between the two winds of the most symmetric Rossmann domain (e2hmvA1). **D.** Structural differences between the 213 and 231 topologies.

Rossmanns have a *β*-strand topology of 321456, often referred to as “doubly wound” because the fold propagates outward from *β*1 and *β*4 in opposing directions (Figure 1D). Consequently, Rossmanns have a pseudo-axis of *C*_2_ rotational symmetry, which may imply that primitive Rossmanns emerged from the joining of two 123 subdomains (one step from a 3-strand precursor), rather than through the independent lengthening at both ends of the *β*-sheet (two steps from a 4-strand precursor; Figure 4A). This evolutionary model makes two predictions: that 123 structures are viable and that symmetric Rossmann structures are viable. A survey of representative ECOD domains reveals that while *α*/*β* proteins with a 123 structure do occur in contemporary proteomes, they often bear decorations that may stabilize the fold (Figure S27A). A symmetry analysis of Rossmann structures suggests that highly symmetric Rossmanns are possible (Figure 4C), as in e2hmvA1, which has a TM-score of 0.70 between the two “winds” of the fold. This Rossmann family is potentially ancient (prokaryote distribution score = 0.84) and, intriguingly, it was excluded from consideration in Figure 1 because the HMM profile associated with this family is mapped to both Rossmanns and TIM barrels in the ECOD hierarchy (see Methods). P-loops, on the other hand, lack a simple repetitive topology.

Alternatively, a 213 half Rossmann with a cross-over between *β*2 and *β*3 may have been the primal Rossmann form. Structures of this type also exist in contemporary proteomes, though once again, with decoration (Figure S27B). With a 213 origin, a single *βαβ* addition or insertion can give rise to one of two Rossmannoid forms, 3214 or 2134, both of which have also been shown to be foldable in protein design studies [20] or are observed as standalone natural folds (Table S6).

Unlike the 123 and 213 topologies, the 231 topology that is relevant to P-loops if P-loops are not derived from an ancient Rossmannoid domain, was not detected in isolation from a manual survey of representative domains (Table S6). The implication, then, is that 123 and 213 may be more accessible fragments, suggesting that Rossmannoid forms—either through duplication of a 321, the extension of a 213, or both—are more readily achieved. Is it reasonable to imagine that 213 and 231 should have radically different properties, given that both structures are three-stranded *α*/*β* proteins with one adjacent and one cross-over *β*-strand? We note that these two forms are not equivalent, as the localization of the loops at the top of the fold, and the crossing point of the *α*-helix differ (Figure 4D). Only in the 213 form are the functional loops adjacent and the *α*-helix most readily available to form a substrate binding pocket. Taken together, these results suggest that the Rossmann topology is more accessible and that Rossmannoid precursors may be more robust starting points for fold origins.

### Rossmann structures are more local

The analysis of protein sequences and structures to infer discoverability follows two primary arguments: The first argument relates to designability [27], the number of sequences that can encode a structure. If a fold has a high designability, the implication is that it occupies a large volume of protein space, and is thus more discoverable by a random sequence-generating process. Previously, Wagner and coworkers proposed that designability is correlated with the contact density of a fold [11]. The second argument relates to foldability, as inferred by topological complexity. Folds with lower topological complexity exhibit faster folding rates and this relationship has been quantified in the measure contact order [41]. Faster folding offers less opportunity for partially folded forms to aggregate [19] or be degraded [39, 8].

An analysis of the contact density of Rossmann and P-loop domains reveals that these two folds are similar (Figure S28A), with generally higher contact densities than other folds (88.2 and 88.1 percentile median contact density, respectively). This result is consistent with the pre-LUCA emergence of Rossmanns and P-loops, as a higher contact density suggests both higher designability and higher discoverability. In contrast, Rossmann domains have a lower contact order than P-loop domains across all domain lengths (Figure S28B)—a straightforward consequence of the more local topology of the Rossmann fold (Figure 4)—implying that Rossmanns are generally faster at folding than P-loops.

### The Rossmann cofactor-binding loop folds early

The relationships among biochemical activity, stability, and foldability are complex. While stability–activity [46, 48, 16] and foldability–activity [28] trade-offs have been argued for specific folds, several analyses have suggested that the extent of these trade-offs can depend on both the structural context and the type of activity involved [49]. Broadly, trade-offs are argued to occur because nothing inherent to polypeptides restricts all or even most biophysical optimizations to be congruent with fast and stable structure generation. Biochemical activity may require, for example, the strained juxtaposition of functional groups or the insertion of flexible loops that impose an entropic penalty on folding. While stability and foldability are distinct properties, the regions of a protein that fold early versus late are determined in part by the relative local stability of structural elements within a fold, leading to a tendency for stability–activity and foldability–activity tradeoffs to be correlated. In contemporary proteins, the cooperativity of the folding reaction allows compensation between parts of the protein that are most constrained by activity requirements and those that can be more directly optimized for stability and/or foldability. The premise that such compensation is a fundamental element of functional exploration motivated the prediction that polarized folds [6], which have a stabilizing core that hosts stability-decoupled decorating loops, are more innovable. In the earliest folds, for which cooperativity, foldability, and stability may all have been most circumscribed, compensation relying on coordination of separated regions would have been rarer, leaving emergent forms to reconcile activity and foldability within or near the active site or restricting them to functional classes that exhibit weaker intrinsic trade-offs.

To clarify how emergent Rossmann and P-loop domains may have overcome these limitations, we applied Wako–Saitō–Muñoz–Eaton (WSME) statistical mechanical models [52, 53, 35, 3, 38], which can estimate residue-level folding tendencies from native state contacts. We present a WSME-based folding analysis for a selection of 11 Rossmann and 10 P-loop proteins (Figure 5A and B). For each domain, a 15-residue fragment centered on the core nucleotide-binding loop (Table S4 and Figure 5C) was identified, hereafter referred to as the cofactor-binding motif (CBM). Folding trajectories for the domains were then calculated using three variants of WSME: the standard implementation using a single-sequence approximation (WSME-SSA) [35], the standard implementation using an exact solution (WSME-exact) [3], and an updated version of WSME that allows for interactions between non-contiguous folded subregions (WSME-L) [38]. For a tutorial introduction to WSME analysis, and a description of the different models, refer to Appendix I and the Methods section, respectively.

**Figure 5:**
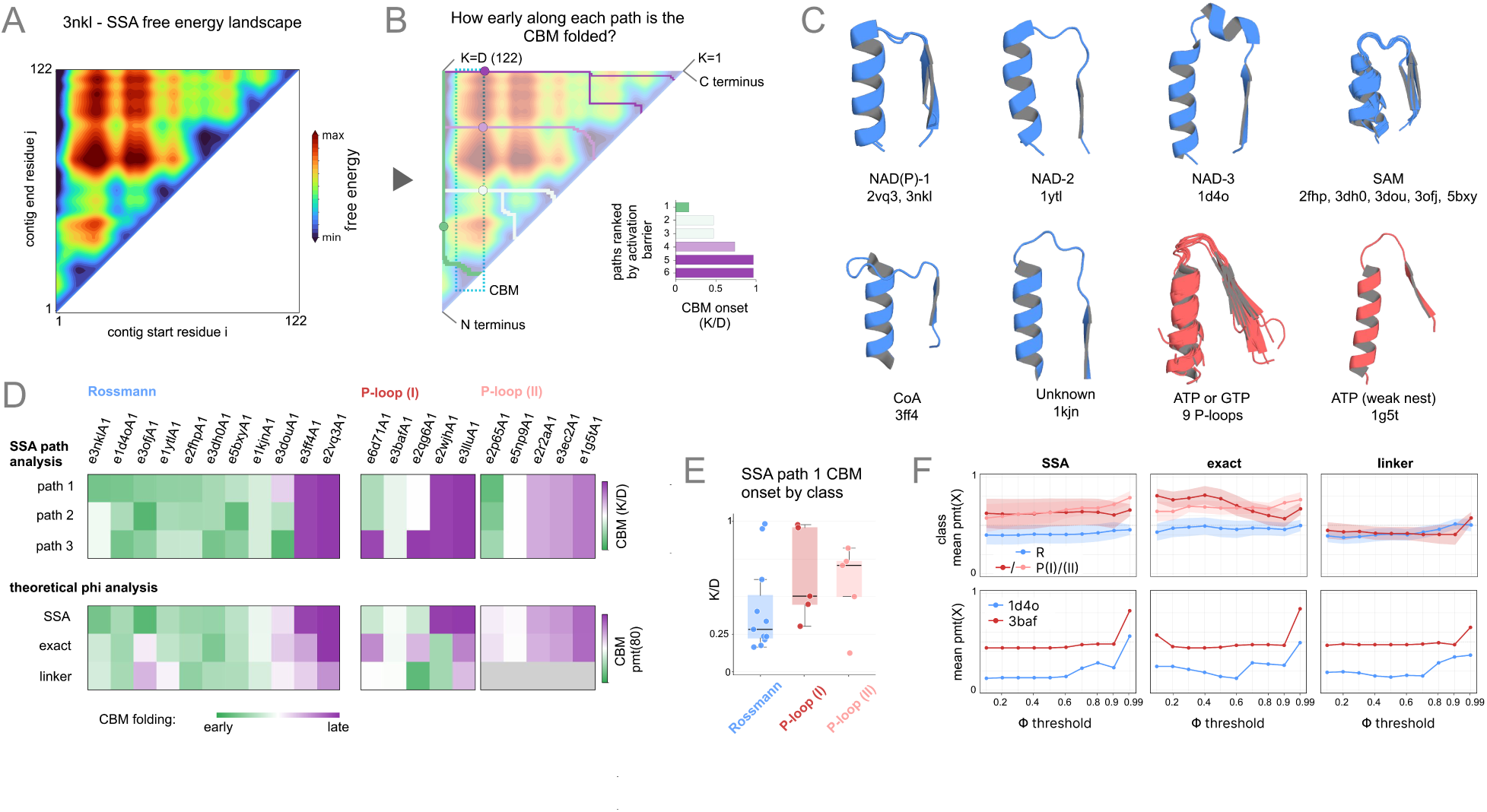
Folding paths of Rossmanns and P-loops. **A.** Contig-based free energy landscape for e3nklA1 estimated via WSME-SSA. **B.** Lowest-action folding paths (each progressing from the diagonal to the upper left all-fold) through the e3nklA1 landscape colored by the point in the trajectory in which the cofactor-binding loop is folded, with green indicating early folding and purple indicating later folding. Inset, paths of the cofactor-binding motif (CBM) ranked by activation-barrier height versus functional loop onset of the normalized contig length (*K*/*D*). Note that the favored paths commit to folding of the CBM earliest. **C.** *βαβ* motifs bearing the cofactor-binding elements for the 21-protein set analyzed here, colored by fold lineage (Rossmann, blue; P-loop, red) and grouped by cofactor and topology. **D.** Top, heatmap of SSA functional loop onset *K*/*D* for the three lowest-action paths. Bottom, heatmap summarizing a theoretical *φ*-value analysis estimating residue-level folding order across all WSME methods. *p*_mt_(*X*) is the percentile rank of the functional loop mean *t_X_* (path-coordinate at which residue *φ*-value first crosses *X*) against the distribution of all possible same-length contigs within the domain; low values (green) indicate cofactor-binding loops that fold earlier than the typical chain segment and high values (purple) indicate later folding. Within each class block, columns are ordered by ascending *K*/*D* for the lowest-action folding path (path-1). WSME-L cells for P-loop type II are excluded (see main text). **E.** SSA path-1 *K*/*D* distribution by class. **F.** Top, class-mean *p*_mt_(*X*) versus *φ*-value threshold *X* for SSA, WSME-exact, and WSME-L. Bands indicate ±1 standard error of the mean. Bottom, the member of each class with the fastest folding CBM identified from SSA (Rossmann, e1d4oA1; P-loop, e3bafA1) traced under all three methods; the earliest Rossmann remains earlier than the earliest P-loop in every case.

Remarkably, we find that the CBM of various Rossmann domains attains its structure early along the folding paths with the lowest activation barriers (Figure 5D top; WSME-SSA). For 5 of the 11 Rossmann domains analyzed, the motif is fully formed within the first quarter of the path (*K*/*D* < 0.25), and the earliest-folding motif, in domain e3nklA1, is formed within the first 16% of the path (Figure 5E). Folding of the nucleotide-binding motif sometime earlier than *K*/*D* = 0.25 implies its participation in an ∼40 residue folding kernel (given an average representative Rossmann length of 163.3 residues). This length scale corresponds roughly to 2–3 secondary structure elements, which have long been recognized as natural packing modules in globular proteins [26]. Later work would identify this length scale in the context of protein folding as a foldon, a quasi-independently consolidating unit of two to three secondary structure elements about 30–40 residues in length [40, 32, 13]. Thus, our results demonstrate that various nucleotide-binding loops are viable participants in the primary nucleating kernels of the Rossmann fold, and thus early folders. This result indicates that these structures are largely neutral to—or potentially supportive of—fold nucleation, given the potential compensations that occur elsewhere in the early folding region.

Perturbative *φ*-value analysis of the lowest-barrier path under WSME-SSA, WSME-exact, and WSME-L similarly finds that the CBM of Rossmann domains can fold early (Figure 5F; *p*_mt_(0.8) < 0.4, where *p*_mt_(0.8) is the percentile rank of the reaction-coordinate position at which the motif’s average *φ*-value first reaches 0.8, relative to the distribution of that same statistic over all same-length contiguous windows in the domain, where 0 = earliest and 1 = latest). By this criterion, 6 domains have early-folding motifs—e1d4oA1, e3nklA1, e3dh0A1, e2fhpA1, e5bxyA1, and e1ytlA1—encompassing multiple CBM types with distinct cofactor preferences. The observation that all three WSME models produce highly similar results, with the average earliness of folding essentially unchanged across the three treatments (*p*_mt_(0.8) ≈ 0.42–0.47), suggests that independent folding of non-contiguous sites (as accounted for in WSME-exact) and inter-contig interactions (as accounted for in WSME-L) are not major contributors to this aspect of the Rossmann folding trajectory. Thus, a roughly foldon-sized context containing a CBM of the Rossmann domain can act as an autonomous folding core whose early folding order is encoded largely by local native contacts.

For two Rossmann domains (e3ff4A1 and e2vq3A1), however, a late-folding motif is observed across all three WSME implementations (Figure 5D; *K*/*D* > 0.9). The late folding of e3ff4A1 likely reflects the pronounced elongation of its nucleotide-binding loop (Figure 5C), and demonstrates that some loop configurations can preclude early folding. The domain e2vq3A1, on the other hand, has both a canonical dinucleotide-binding motif and a canonical fold topology, yet its CBM folds late. This result is consistent with the observation that folding nuclei can be [17] but are not strictly conserved [23].

P-loop CBMs behave less uniformly than the Rossmann motifs and are formed later along their folding paths, with the motif of just 1 of the 10 domains folded at *K*/*D* < 0.25 (Figure 5E) and none, on average under local-contact treatments, reaching *p*_mt_(0.8) < 0.4 (Figure 5F). However, admitting non-local linker interactions with WSME-L yields markedly earlier folding within type I domains (average *p*_mt_(0.8) of 0.60 for WSME-exact versus 0.41 for WSME-L), bringing them in line with the average Rossmann value (average WSME-L *p*_mt_(0.8) of 0.41 and 0.46 for type I P-loop and Rossmann domains, respectively; these class means include the late-folding outliers). We interpret this result as evidence for contextual early folding: only with non-local support from the C-terminal subdomain does the P-loop CBM fold ahead of the background. Thus, although the P-loop CBM is not consistently part of the nucleating region of the fold, it appears to be folding-permissive rather than obstructive, at least relative to the more topologically complex parent P-loop domain. We hypothesize that in the presence of ligands, more consistent early folding of the P-loop (and Rossmann) cofactor-binding motifs may be observed.

### An integrated reconstruction of the Rossmann and P-loop early evolution

The observation that motifs associated with biochemical activity are predicted to fold early, particularly within Rossmann domains (Figure 5), confirms that activity (in this case, cofactor binding) and foldability can be achieved simultaneously, while also lending specific insights into the contexts that avoid the activity tradeoffs. The congruence of nucleation and active-site formation may both explain the early emergence of the Rossmann cofactor-binding loop and connect that priority to its extraordinary subsequent longevity. The late folding of a non-canonical cofactor-binding motif that bears a long insertion (Figure 5D) suggests that the compact nature of the Rossmann functional loop—and, potentially, its participation in a mini-core formed by *β*1, *β*2, and *α*2 (Figure S29)—is partially responsible for its early attainment along the folding trajectory.

If early discovery and evolutionary retention benefits from folding robustness without compromise for function, and if short-loop forms most readily confer this benefit, we may expect that the associated families will be enriched for cofactor-binding, as a way to off-board both functional activity and evolvability to modular small molecules and off the body of the protein. Previously, analyses of the CheY family of flavodoxins [31, 34]—notably, the main other large group with characteristics similar to those of Rossmanns shown here (Figure 3, Figure 4A)—observed highly conserved residues in a short loop that is part of a folding nucleus, a feature that is also reflected by WSME folding analysis (Figure S31). The suggestion that rigidity may support cofactor binding and foldability alike furnishes one of the criteria by which selection correlates early discoverability with a preference for cofactor-binding.

The relatively compact loop structure of the Rossmann motif, in addition to being folding competent, appears to have favored functional exploration. We hypothesize that Rossmanns have been uniquely successful at cofactor switching in part because the core binding *βαβ* motif is a surface against which cofactors bind, not a hand that wraps around the cofactor, as in the case of P-loops (Figure 1C). We note that this feature is largely retained across Rossmann-binding modes (Figure 1C and Figure S30), even when considering long-loop forms (Figure 5C). It has been demonstrated that the NAD-binding Rossmann *βαβ* can be converted to a SAM-binding site by a simple 3-residue deletion [51]—in effect, by changing the topography of the flat surface. In contrast, with its hand-like grasp of di- and triphosphate moieties, the Walker A motif has primarily admitted transitions between mononucleotides. Thus, folding constraints during the emergence of the Rossmann domain favoring flatter, more compact loop structures may have ultimately produced a more successful platform for cofactor diversification by enforcing fewer constraints on the properties of its cognate ligands. When viewed this way, the observation that Rossmanns are predicted to be early by metabolic constraint modeling (Figure 2D) is causally linked to the folding constraints that shaped the emergence of this fold.

We find that the Rossmann fold is (a) central in fold space (Figure 3), (b) associated with more local interactions (Figure S28B), and (c) characterized by a simple, repetitive internal structure (Figure 4C). Whereas structural repetitiveness suggests a facile pathway for emergence [9, 24, 45], structural centrality and localness are signatures of biophysical optimality. In a context where the joint problems of high-fidelity translation and robust folding are actively being solved, these features imply a greater degree of biophysical robustness that would support emergence and persistence. Moreover, the Rossmann cofactor binding motif operates within the context of a simple, robust fold, without the need for a high degree of structural complexity.

Taken together, our results suggest a bias toward comparatively passive roles for the first coded peptides, in which simple peptides entered a system dominated by cofactor-mediated chemistry and found utility in their association with cofactors, metals, and RNA. While the metaphor of the palimpsest [2] has been attractive from perspectives on biogenesis assuming genetics-first and anchored in the Central Dogma, we infer a more graded and partial hand-off of molecular biology from small molecules to polymers: participation through binding and modest catalytic enhancement, followed by accretion into simple, stable structures that in turn supported more complex chemical roles. In short, emergent systems may have been conservatively supported by nascent proteins and peptides before their exquisite potential for structural choreography and functional innovation could be fully realized.

## Acknowledgements

We gratefully acknowledge support from the National Aeronautics and Space Administration (NASA) under awards 80NSSC25K7873 (LML, SCLK) and 80NSSC24K0344 (ES), the Human Frontier Science Program (HFSP) under award RGEC29/2025 (LML), the Japanese Society for the Promotion of Science under award 26KJ1170 (TC), the Zuckerman STEM Leadership Program (CLSD), the Data Science Research Center at the University of Haifa (CLSD, RK), and the Israel Science Foundation (ISF) under award 1764/21 (RK). Travel support from the Earth-Life Science Institute (ELSI) (LML) facilitated international scientific exchange that made this work possible.

## Author Contributions

Conceptualization: KS, TC, HG, RK, ES, LML Investigation: KS, TC, HG, ALB, CL

Data curation: KS, TC, HG, ALB, CL

Formal analysis: KS, TC, HG, ALB, CL, RK, ES, LML Software: HG, ALB

Visualization: KS, TC, HG

Methodology: KS, TC, HG, ALB, CL, SCLK, RK, ES, LML

Project administration: KF, RK, ES, LML

Funding acquisition: RK, SCLK, LML

Supervision: KF, RK, ES, LML

Writing – original draft: KS, TC, HG, ES, LML

Writing – review & editing: ALB, CL, KF, SCLK, RK

## Declaration of Generative AI and AI-assisted Technologies

During the preparation of this work, the authors used Claude (Anthropic) to identify typographical or grammatical mistakes and assist with scripting. All suggestions were manually reviewed and selectively incorporated into the text and code. The authors take full responsibility for the content of the publication. AI models were used for literature searching with human oversight and validation.

## Methods

### Definition of Rossmann and P-loop evolutionary lineages

Protein evolutionary classification follows the Evolutionary Classification of Domains (ECOD) database [57]. The Rossmann and P-loop fold lineages correspond to X-groups 2003 (Rossmann-like) and 2004 (P-loop NTPase-like), respectively. Families within a fold lineage (X-group) correspond to F-groups in the ECOD hierarchy. Note that Rossmannoid fold lineages such as flavodoxin (X-group 2007) are not considered Rossmann domains as defined here [33].

### Protein domain mapping, proteome scaling plot calculation, and phyletic distribution score calculation

Rossmann and P-loop domains were identified within the representative proteomes from the Genome Taxonomy Database (GTDB) version 220 [58], which includes both bacteria and archaea. Searching was performed with HMMER version 3.4 (http://hmmer.org) using the ECOD HMM profiles. All HMM profiles within ECOD, including those that do not belong to the Rossmann or P-loop evolutionary lineages, were considered. By considering HMM profiles from all evolutionary lineages, rather than just those associated with the Rossmann or P-loop evolutionary lineages, we minimize the potential misclassification of Rossmannoid domains as Rossmanns. HMM profiles that are associated with multiple X-groups in version 289 of the ECOD representative domains file were excluded from consideration. Hits were filtered according to an independent E-value (i-Evalue) cutoff = 0.01 and a fractional HMM profile coverage cutoff = 0.75. The size of the search space was set to be the total number of genes analyzed, *Z* = 333,384,550 genes. In the event that multiple domains map to the same region of a sequence and the overlap is above a fractional domain overlap cutoff = 0.75, the domain with the lowest i-Evalue was given priority. An implementation of the domain mapping protocol is provided at https://github.com/AndreLecona/Dotate.

Scaling plots were calculated as the total number of Rossmann or P-loop domains versus the total number of annotated domains within a proteome, referred to here as the proteome size. Proteomes within each proteome size bin are summarized as a mean ±1 standard deviation. Summary statistics were calculated after log transformation with genome quality scores, which reflect the estimated completeness and extent of contamination of a genome, used as weights.

Distribution scores for each Rossmann and P-loop family were calculated using the GTDB Bacteria and Archaea phylogenetic trees as

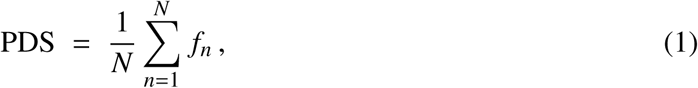

where *f_n_* is the fraction of genomes containing a given family within phylum *n*, and *N* is the number of phyla.To calculate a prokaryote phyletic distribution score, the phyletic distribution scores for bacteria and archaea were averaged. The phyletic distribution scores for each Rossmann and P-loop family are provided in Table S1.

### P-loop topology classification

P-loop families were classified into three topology types (0, I, and II) based on the *β*-strand order of the interior *β*-sheet. Type I P-loop proteins have a single intervening *β*-strand between *β*1 (which directly precedes the Walker A motif) and *β*2. The canonical *β*-sheet topology of the Type I P-loop core is 2-3-1-4-5. Type II P-loop proteins have two intervening *β*-strands between *β*1 and *β*2, with a canonical core *β*-sheet topology of 2-3-4-1-5. Topology classification was performed manually using representative structures (Table S2). One family, 2004.1.1.133, is associated with both Type I and Type II *β*-sheet topologies. One family has no intervening strand between *β*1 and *β*2 (topology: 2-1-3-4) and is referred to here as a Type 0 P-loop.

### Metabolic network expansion

The metabolic model described by Goldford *et al.* [12] was used with a single modification: the reactions producing H_2_O_2_ before the emergence of oxygenic photosynthesis were removed, as in [5]. This model, which includes both cofactor–reaction and enzyme–reaction associations, was derived from the Kyoto Encyclopedia of Genes and Genomes (KEGG) database [59] and corresponds to a biosphere-scale metabolic map. We note that constraint-based models such as these involve several core assumptions; see [60] for a detailed discussion. They do not enforce the need for biological energy conservation.

To analyze the metabolic roles of the Rossmann and P-loop evolutionary lineages at different stages of metabolic evolution, the network expansion algorithm [14] was employed. Briefly, this approach uses a proposed set of seed compounds and the constraints imposed by a metabolic model to produce a constraint-based pseudo-temporal reconstruction of reaction and metabolite discovery order. The seed compounds used here include prebiotically available compounds and organic substrates that could have plausibly been supplied by a prebiotic Wood–Ljungdahl-braced rTCA carbon fixation pathway [61]. The seed compounds are listed in Table S3 and are the same as those used in [12].

To explore the roles of Rossmanns and P-loops in early metabolic evolution, two analysis types were performed: association and ablation. In the first analysis, the number of reactions that are associated with either evolutionary lineage is enumerated at each stage of the metabolic expansion trajectory. In the second analysis, reactions that require either the Rossmann or P-loop evolutionary lineage at each iteration were ablated to determine the extent to which the final number of discovered compounds (scope) is contracted. Note that by eliminating reactions at each iteration, interdependence between reactions from different iterations is not accounted for. The network expansion algorithm is available as a Python package on GitHub (https://github.com/jgoldford/networkExpansionPy).

### *βαβ* **motif identification**

Unlike previously reported approaches [62], we also consider *βαβ* motifs in which the two *β*-strands are not directly interacting, referred to here as *open βαβ* motifs. Open *βαβ* motifs are common in Rossmann domains (namely, as the cross-over *βαβ* motif between the two opposing winds of the doubly wound topology) and P-loop domains. First, secondary structure assignment was performed with the Define Secondary Structure of Proteins (DSSP) algorithm version 4.6.1 [63] to identify all *β*-strands and *α*-helices within a domain. Next, a *β*-strand interaction network was constructed in which individual *β*-strands (at least 3 contiguous residues) are represented as nodes. Edges are introduced between pairs of nodes when the corresponding *β*-strands shared two or more inter-strand hydrogen bonds and the angle between the strand vectors, defined from the N-terminal to the C-terminal residue of the *β*-strand, is below a given threshold *θ* = 40^◦^.

If the angle is less than 40^◦^, the strands are considered to be parallel; if the angle is near 180^◦^ (|angle − 180^◦^| < 40^◦^) the strands are considered to be antiparallel. Connected components in this network were taken to represent *β*-sheets. Note that both parallel and antiparallel *β*-strands are included in the *β*-sheet network. Finally, pairs of *β*-strands were classified as forming a *βαβ* motif if (i) they belong to the same connected component of the strand network (i.e., are part of the same *β*-sheet, either directly or indirectly connected), (ii) are linked by at least one intervening *α*-helix of any length in the primary sequence, and (iii) have a parallel *β*-strand orientation. Note that a single *β*-strand may participate in up to two *βαβ* motifs, as either the first or second *β*-strand of the *βαβ* motif. A graphical explanation of the algorithm is provided in Figure S1. A comparison with PROMOTIF [62], which can identify closed *βαβ* motifs only, is presented in Figure S2. The sequences and structures of *βαβ* motifs were extracted from the representative structures of ECOD version 291 clustered at 40% identity (F40). We find that 92% (19,105/20,815) of *βαβ* motifs identified by PROMOTIF are identified by our approach as well. In addition, our approach identifies 27,986 *βαβ* motifs not identified by PROMOTIF, 57% (15,906/27,986) of which are open *βαβ* motifs. A Python implementation of the *βαβ* motif finder algorithm can be downloaded at https://github.com/liam-longo-lab/BABMiner.

### *βαβ* **similarity network analysis**

Similarity networks between protein evolutionary lineages were constructed by a pairwise comparison of *βαβ* motifs. *βαβ* motifs were extracted from the F40 representative domains of ECOD version 289, yielding 50,611 *βαβ* motifs in total (Figure S3, Figure S4). An all-versus-all comparison of the identified *βαβ* motifs was then performed using TM-align [64]. To enforce structural similarity as well as length similarity between *βαβ* motifs, the minimum of the two TM-scores reported by TM-align (referred to here as the minimum TM-score) was used for each *βαβ* motif pair (Figure S5). *βαβ* similarity networks were constructed such that each node corresponds to a *βαβ* motif and edges are drawn between similar *βαβ* motifs, subject to a minimum TM-score cutoff (Figure S6). X-group similarity networks were constructed such that each node corresponds to a protein evolutionary lineage and edges are drawn between X-groups that are associated with similar *βαβ* motifs. In addition to a minimum TM-score cutoff that defines similar *βαβ* motifs, X-group similarity networks are subject to an additional domain count threshold, which sets the number of domains of each X-group that must be connected to each other by similar *βαβ* motifs for an edge to be drawn. This additional criterion minimizes the potential consequences of a small number of misclassified domains.

### Identification of highly symmetric Rossmann domains

Rossmanns containing (i) exactly five annotated *βαβ* motifs and (ii) a single open *βαβ* motif with two intervening *β*-strands as the third *βαβ* motif in the sequence were identified. The *βαβαβ* elements (corresponding to two *βαβ* motifs with an overlapping *β*-strand) on either side of the open *βαβ* motif were then extracted and aligned using TM-align.

### Protein language model latent space analysis

Embeddings for each of the representative domains from ECOD F40 version 289 were calculated using the protein language model Contrastive Learning Sequence–Structure (CLSS) using both the CLSS-sub model (which was trained on randomly selected protein sub-sequences of ≥ 10 residues in length) and the CLSS-full model (which was trained only on full-domain sequences) [56]. Pairwise cosine distances between all domains were calculated from the 32-dimensional CLSS embeddings. To generate similarity networks between X-groups, two limits were imposed: a cosine distance cutoff, *D*, and a domain count threshold, *n*. As above, the domain count threshold requires a given number of domains from both evolutionary lineages under consideration be within a certain distance in the CLSS embedding space for an edge between X-groups to be drawn.

### Contact density and contact order calculation

Residue–residue contacts were computed from atomic coordinates for a filtered subset of ECOD representative domains (v289), clustered with CD-HIT (*c*=0.7, *n*=5). Domains were excluded from consideration for the following reasons: (i) a crystallographic resolution ≥ 2.5 Å, (ii) the presence of inserted domains, and (iii) a length shorter than 80 amino acids or longer than 300 amino acids. Two residues were considered to be in contact if any non-hydrogen atom of one residue, including side-chain atoms, was within 6 Å of any non-hydrogen atom of the other, based on the heavy-atom contact definition used by [65]. To better emphasize the global fold topology, residue pairs separated by two or fewer intervening residues in sequence (|*i* − *j* | ≤ 2) were excluded from consideration. Contact density (CD) was defined as the number of contacting residue pairs normalized by domain length:

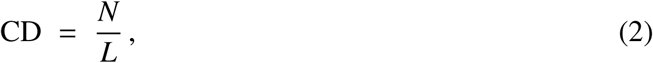

where *N* is the total number of contacting residue pairs satisfying the constraints above, and *L* is the number of residues in the domain. Relative contact order (RCO) was calculated as

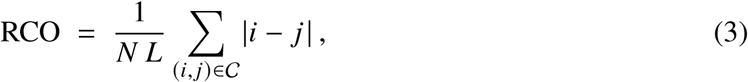

where |*i* − *j* | is the sequence separation between residue *i* and residue *j* for each contacting pair, C is the set of contacting pairs, *N* = |C| is the number of contacting pairs, and *L* is the domain length. As with contact density, residue pairs separated by two or fewer intervening residues in sequence (|*i* − *j* | ≤ 2) were excluded from consideration.

### Energy decomposition of representative Rossmann and P-loop structures

Representative Rossmann and P-loop structures were selected for folding analysis based on five criteria: (i) an experimentally determined structure with a crystallographic resolution of ≤ 2.75 Å, (ii) less than 200 residues in length, (iii) an absence of large inserted (nested) domains, (iv) a canonical strand topology, and (v) apparent folding independence. One P-loop domain (e5np9A1) has a non-canonical *β*-strand topology in which *β*2 of 51432 is in the antiparallel orientation. In the case of oligomers, only domains that can be reasonably hypothesized to fold prior to oligomerization (as opposed to a coupled process) were considered. For models with short (2–9 residues) regions of missing residues and/or selenium analogs of methionine and cysteine, Modeller version 10.7 [66] was used to generate the full-domain structure with canonical amino acids only. In total, 21 representative ECOD domains were selected for folding mechanism analysis: eleven Rossmann domains, five Type I P-loop domains, and five Type II P-loop domains. Details about each of the representative domains are provided in Table S4.

To calculate residue–residue interaction energies, the structures were prepared and equilibrated as described in [67]. Briefly, water molecules and ions were stripped from the experimental structures, and only one protomer was retained for homocomplexes. Using AmberTools25 [68], each structure was solvated in a truncated octahedral water box containing TIP3P water molecules [69]. The box extended 10 Å from the protein in all directions. Sodium or chloride ions were added to neutralize the system. The standard protonation states of ionizable side chains at physiological pH were used, as determined by AmberTools25. Protonation states were further validated using PROPKA3 [70].

Five replicas of each structure were independently equilibrated as follows. All simulations used the SHAKE algorithm [71] for bonds involving hydrogen, and the AMBER ff14SB force field [72]. Equilibration simulations were performed with a timestep of 1 fs, while the production simulation was performed with a timestep of 2 fs. The system was equilibrated using the following steps:

1. 1,000 steps of energy minimization (20 steps of steepest descent followed by 980 steps of conjugate gradient) under constant volume (NVT simulation), without positional restraints, to remove any poor initial contacts;
2. gradual heating of the system from 100 K to 300 K over 1 ns. During this heating step, all protein atoms were restrained using 100 kcal mol^−1^ Å^−2^ harmonic restraints;
3. equilibration at 300 K (as for all subsequent steps) for 1 ns, under constant isotropic pressure with isotropic volume changes (NPT simulation), keeping the restraints from step (ii);
4. NPT simulation for 1 ns, reducing the restraints on all protein atoms to 10 kcal mol^−1^ Å^−2^;
5. NVT simulation for 1,000 steps, with a restraint of 10 kcal mol^−1^ Å^−2^ on the protein backbone atoms;
6. NPT simulation for 1 ns, with restraints on the protein backbone atoms of 10 kcal mol^−1^ Å^−2^;
7. NPT simulation for 1 ns, reducing the restraints on the protein backbone atoms to 1 kcal mol^−1^ Å^−2^;
8. NPT simulation for 1 ns, reducing the restraints on the protein backbone atoms to 0.1 kcal mol^−1^ Å^−2^; and
9. NPT simulation for 1 ns with no restraints on any of the protein atoms.

Finally, the production MD was performed for 2 ns in the NPT condition at 300 K and 1 atm, using a Langevin thermostat (collision frequency 2 ps^−1^) and an isotropic Monte Carlo barostat. A 2 fs timestep was used, with coordinates saved every 10 ps. No positional restraints were applied.

Pairwise residue–residue interaction energies were computed using the Molecular Mechanics Poisson–Boltzmann Surface Area (MM/PBSA) method as implemented in MMPBSA.py [73]. For each replica, 20 snapshots were extracted from the production MD simulation at 100 ps intervals. The Poisson–Boltzmann calculations were performed using a nonlinear solver with an ionic strength of 0.15 M, and per-residue pairwise energy decomposition was carried out over all residues. The resulting pairwise interaction energies were assembled into residue–residue energy matrices and subsequently used as contact maps for downstream analysis. In each case, the equilibrated structure and the pairwise interaction energies between replicas were highly similar (Figure S7). Thus, for subsequent folding mechanism analyses, the C*_α_*–C*_α_* distance matrix and the pairwise residue interaction energies were averaged across the 5 independent energy-minimized replicas.

### Wako–Saitō–Muñoz–Eaton (WSME) folding pathway analysis

WSME is an Ising-type statistical mechanical model in which each protein residue occupies a folded or unfolded site state. The Hamiltonian for WSME models is built from native pairwise contact energies under a contiguity ansatz that permits a folded residue to interact with its native partners only when every intervening residue along the chain is also in the folded state. Three WSME variants were considered in this work, and are described in greater detail below. All three variants take the pairwise interaction energies, described above, as an input. Each WSME variant requires a per-residue entropic cost that governs the balance between the enthalpic gain of forming native contacts and the conformational penalty of ordering a residue; the specific value or determination procedure differs across the three methods and is treated as a hyperparameter (Table S5).

The three WSME variants considered here differ in their: (i) complexity, (ii) choice of order parameter, (iii) partition function construction/solution class, and (iv) hyperparameters. We provide a tutorial introduction to WSME-based folding mechanism modelling, including a detailed presentation of each model, in Appendix I.

#### Single-sequence approximation (SSA)

The simplest WSME variant uses a single-sequence approximation (SSA) to the WSME partition function [35] which restricts the sum to microstates containing at most one contiguous folded segment or contig. A domain of *D* residues admits *D*(*D* + 1)/2 possible contigs, which form a triangular lattice-based free-energy landscape (Figure 5A, Figure S8) with contigs of length 1 (individual residues) arrayed on the diagonal. The SSA is defined by an order parameter *K*, the contig length. A generic folding trajectory in this landscape is a continuous path connecting the (*K*=1) diagonal (contig start residue *i* = contig end residue *j*) with the terminal point (*K*=*D*, *i*=1, *j*=*N*). The order parameter therefore indexes a folding path. SSA landscape evaluation scales as *O*(*D*^2^) and a complete residue-level *φ*-value analysis—in which the native interactions of a target residue are dampened in the Hamiltonian and the partition function recomputed (described in more detail below)—scales as *O*(*D*^3^). The SSA has a single free parameter, the per-residue entropy penalty Δ*S*, which is set analytically from the constraint

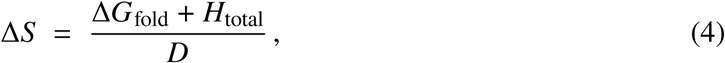

where *H*_total_ is the total contact enthalpy and Δ*G*_fold_ is the target folding stability (set here to a default value of −10 *k_B_T*; Table S5). Folding trajectories through the SSA landscape are enumerated and ranked by the Incursion-to-Topological-Network (I-TN) procedure described below.

#### WSME-exact

The WSME partition function can also be evaluated without approximation [3], WSME-exact, by a transfer-matrix dynamic-programming recursion that enumerates all possible combinations of non-overlapping contiguous folded stretches. Unlike SSA, multiple contigs can coexist in a single microstate, although the WSME contiguity ansatz still applies: a pairwise contact contributes to the energy only when every intervening residue is folded, so contacts between residues residing in different contigs are excluded. Because the state space now contains arbitrary combinations of non-overlapping contigs rather than a single stretch, the contig-length coordinate *K* of the SSA no longer suffices, and the order parameter becomes *q*, the total number of folded residues summed across all contigs. A single landscape evaluation scales as *O*(*D*^3^), and a complete *φ*-value analysis scales as *O*(*D*^4^). WSME-exact requires the contact energy map and a per-residue entropic cost vector; following Ooka and Arai [38], the per-residue entropic cost is fixed at Δ*S* = −3.5, leaving the energy scaling *ε* as the only free parameter, which is selected by minimizing

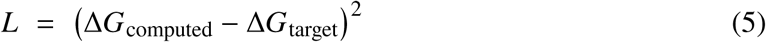

over a 1D adaptive grid search in *ε*, with Δ*G*_target_ = −10 *k_B_T* (Table S5).

#### WSME-L

Building on the WSME-exact model, Ooka and Arai [38] developed WSME-L, which introduces virtual linkers between non-contiguous folded stretches, capturing cross-gap interactions that may alter the thermodynamic viability of folding paths. Each virtual linker incurs a ring entropy penalty that is a function of C*_α_*–C*_α_* distances. Thus, WSME-L uniquely requires a C*_α_*–C*_α_* distance matrix as input. WSME-L has two free parameters: the energy scaling *ε* and an entropy scaling *H_S_* that modulates the ring-entropy terms. These parameters are jointly optimized via a two-dimensional grid search minimizing

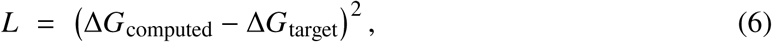

where Δ*G*_computed_ is the free-energy difference between the native and unfolded basins of the computed landscape and Δ*G*_target_ = −10 *k_B_T* [38]. Following Ooka and Arai [38], the per-residue entropic cost is fixed at Δ*S* = −3.5. WSME-L uses the same order parameter *q* as WSME-exact. To identify the dominant folding route and determine which structural block consolidates first, the *D* residues are partitioned into two blocks of size *n*_1_ and *n*_2_ (*n*_1_ + *n*_2_ = *D*), and the partition function is decomposed into two order parameters (*q*_1_, *q*_2_) counting the folded residues in each block. This approach yields a two-dimensional free-energy landscape *G*(*q*_1_, *q*_2_). Note that the domain split determines only the reaction-coordinate decomposition; virtual linkers may connect any residue pair (*u*, *v*) with a sufficiently attractive contact energy, whether within the same block or across blocks. An optimal one-dimensional folding coordinate through *G*(*q*_1_, *q*_2_) is then obtained by computing a monotone minimax path—the trajectory from the unfolded basin (minima in the region defined by (*q*_1_/*n*_1_, *q*_2_/*n*_2_) < 0.5) to the native basin (minima in the region defined by (*q*_1_/*n*_1_, *q*_2_/*n*_2_) > 0.5) that minimizes the maximum barrier height (rate-limiting transition state) encountered. WSME-L effectively scales as *O*(*C*_link_ · *D*^5^) per landscape, where *C*_link_ is the number of contacts below the linker cutoff (contact energies *e_u_*_,*v*_ < −0.6). A complete *φ*-value analysis repeats this calculation for each of the *D* residues in a domain. Due to this cost, and because Type II P-loops lack an unambiguous two-block partition required by the two-dimensional reaction coordinate of WSME-L, WSME-L was run on a sixteen-protein subset, excluding Type II P-loops (Table S5).

For all methods presented, the results of the analyses are robust to the choice of Δ*G*_fold_ (Figures S9 and S10). In addition, comparisons of replicate averages to results obtained from single replicates are highly similar (Figures S11 and S12).

### Relative folding precedence of the cofactor-binding loop: path-based approach (SSA)

The SSA landscape is indexed by the start and end residues (*i*, *j*) of the sole folded contig; the contig length *K* = *j* −*i* + 1 then labels the lattice diagonal and indexes path progression through the landscape (Figure 5B). We identified and ranked paths using an Incursion-to-Topological-Network (I-TN) procedure comprising three stages: incursion detection, network construction, and pathway assembly. A pseudocode implementation of this approach is described in Figure S13.

**Incursion detection.** Folding nucleation sites are located on or near the landscape diagonal (*K*=1). The Gaussian-smoothed landscape is thresholded at a low percentile of its free-energy distribution to mark cells that dip below the diagonal baseline, i.e., residue positions that can initiate spontaneous folding. These sub-baseline cells are partitioned into candidate incursion valleys by a watershed seeded at the landscape’s regional minima and trimmed to the residue window of each valley’s deep core; a valley is retained as a non-trivial incursion only if it is large enough and anchored at the diagonal (its minimum contig length reaches *K* ≈ 1). Each retained incursion is subsequently extended into the landscape interior with strictly increasing *K* during pathway assembly (below).

**Network construction.** The Gaussian-smoothed SSA landscape is partitioned into basins. The sign of the Hessian determinant first identifies critical-point territory, in which positive-determinant patches host local extrema (split into minima and maxima by the Hessian trace) and negative-determinant patches host saddles. Minima with contig length *K* ≤ *K*_thr_ are discarded, and the landscape interior is tessellated into basins by watershed segmentation seeded at the surviving minima. Adjacent basins are connected through the minimax-pass cell on their shared boundary, and each saddle together with its connecting minimum-energy path is refined by a simplified string method [74]. The nodes of the resulting Topological Network T are these refined minima and its edges record the refined first-passage saddles and their barrier heights, yielding a reduced graph of all kinetically relevant basins and the rate-limiting transitions between them.

**Pathway assembly.** Each incursion is anchored to the diagonal and stitched onto a *K*-monotone lattice path: a minimax segment runs from the (*K*=1) diagonal up to the incursion’s deep-core centre, and a second minimax segment continues from that centre to the native attractor at (*j*, *i*) = (*D*−1, 0). Both segments use only the two single-tile-flip moves that each raise *K* by one, so the assembled polyline is strictly *K*-monotone by construction. Each step minimises the maximum free energy encountered along the path (the rate-limiting bottleneck), with ties broken toward lower cumulative energy so the route hugs the valley floor. Every path’s bottleneck cell is then labelled with its nearest saddle in T, locating the rate-limiting transition on the reduced network. Pathways are ranked by their bottleneck barrier (the maximum free energy along the path), so that rank 1 is the lowest-barrier, most accessible folding route. For each ranked path, the cofactor-binding loop folding onset is defined as the normalized contig length *K*/*D* at which the growing contig first fully spans the cofactor-binding loop window. A low *K*/*D* value indicates that the functional loop becomes structured early on the folding path, that is, when only a small fraction of the domain is folded (Figure 5C).

### Relative folding precedence of the cofactor-binding loop: path-based approach (WSME-L)

Under WSME-L, the unfolded and native states are fixed by construction at the corners of the landscape, (0, 0) and (*n*_1_, *n*_2_) respectively. The optimal one-dimensional folding path is constructed as a monotone-minimax path in three segments,

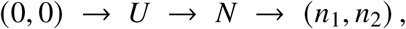

where *U* and *N* are the lowest-energy local minima within the lower-left and upper-right quadrants of the landscape. Each segment is required to be non-decreasing in both coordinates, with every step increasing exactly one order parameter by one (Δ*q*_1_ +Δ*q*_2_ = 1; 4-connected single-flip moves), and minimizes the maximum free-energy barrier traversed. The transition state of the path is the highest-energy cell of the *U* → *N* segment. For the Rossmann domain e3nklA1, whose two transition states are near-degenerate (bottleneck barriers within ≈ 0.14 *k_B_T*), we report the symmetry-related alternative pass. A pseudocode implementation of this approach is described in Figure S13.

### Relative folding precedence of the functional loop: perturbative ***φ***-value analysis (all models)

For each protein, the nucleotide-binding functional loop was defined as the 15-residue contiguous stretch spanning the annotated catalytic *βαβ* motif at the cofactor- or Walker-binding site (Table S4). Under WSME-exact and WSME-L, the order parameter *q* reports on global folding progress but does not directly encode which residues fold first. To extract residue-level folding order, we applied the perturbative theoretical *φ*-value analysis of Ooka and Arai [38]; see also [75]. This approach attempts to determine the order in which individual residues acquire their native energetic environment along the folding coordinate: if weakening residue *l*’s native contacts shifts the free energy at an intermediate folding stage *q* by a fraction comparable to the shift it produces in the fully native state, then residue *l* has already engaged its native partners by stage *q* and is folded earlier; if the perturbation has a negligible effect at *q*, residue *l* has not yet acquired its native context and folds later. Aggregating this comparison across all residues yields a complete residue-by-stage map of folding precedence that requires no assumption beyond the WSME ansatz already used to construct the landscape. For each residue *l* = 1, . . ., *D*, all attractive native contacts involving *l* are weakened by a factor *δ* = 0.1, the full partition function is recomputed with the perturbed Boltzmann weights, and the *φ* value *φ_l_* (*q*) is obtained as the ratio of perturbation-induced free-energy shifts at the intermediate and fully folded points of the relevant landscape:

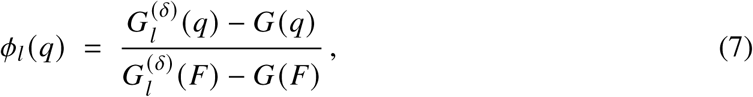

where *G*^(*δ*)^ (*X*) is the WSME free energy at landscape position *X* after weakening residue *l*’s native contacts by factor *δ*, *G*(*X*) is the unperturbed reference, and *U* and *F* are the unfolded and fully folded endpoints of the relevant landscape (Figure S14).

The result is a residue-by-reaction-coordinate *φ*-value matrix for each protein (Figure S15). Only WSME-exact (reaction coordinate *q*, with *U* = 0 and *F* = *N*) is one-dimensional by construction. For SSA and WSME-L the *φ* field is defined on a two-dimensional lattice—the contig lattice (*i*, *j*) for SSA and the two-block landscape (*q*_1_, *q*_2_) for WSME-L (with *U* = (0, 0) and *F* = (*n*_1_, *n*_2_))— and is reduced to a residue-by-path-step matrix by sampling along the folding path: the rank-1 I-TN path for SSA, and, for WSME-L, the monotone-minimax path that is non-decreasing in both *q*_1_ and *q*_2_ and minimises the maximum free-energy barrier traversed. In every case the sampled coordinate is parameterised by arc length so that residue-level results from all three variants share a common normalised 1D folding coordinate.

To summarize functional loop precedence from the *φ*-value heatmaps, we extracted threshold-crossing times *t_X_* for each residue: the normalized reaction coordinate at which *φ* first reaches threshold *X* (we report *t*_0.8_ in the main text; see Figure S16 for a comparison across thresholds), determined by linear interpolation. The functional loop was then summarized by the mean *t_X_* across its constituent residues. Because raw *t_X_* values are not directly comparable across proteins of different length or landscape topology, each summary statistic was converted to a percentile rank against a sliding-window null distribution: all contiguous windows of the same length in the same protein were scored with the identical metric, and the functional loop window’s rank within that distribution was reported. A percentile near zero indicates that the functional loop region folds earlier than nearly all other windows of comparable size in the same domain. Complete hyperparameter settings for the three methods are provided in Table S5.

