## Supplementary information for "Nucleotide-binding motifs nucleated folding of the first enzymes"

[†]Equal contribution.

[*]To whom correspondence should be addressed.

---

### 1 Supplementary figures

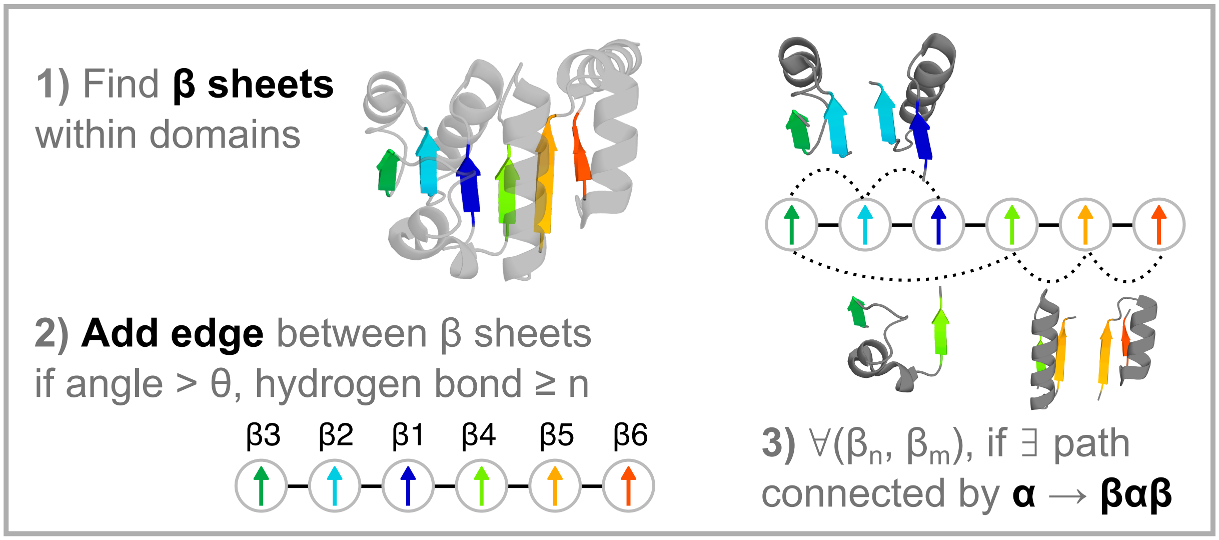

**Figure S1: Graphical explanation of the $\beta\alpha\beta$ motif finder algorithm.** See Methods for additional details.

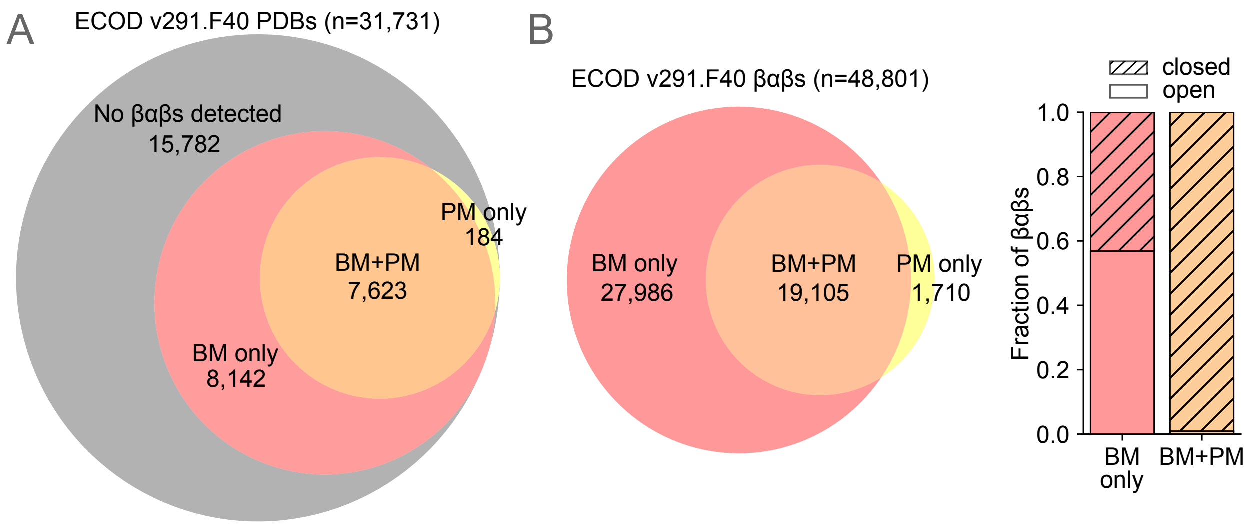

**Figure S2: Comparison of $\beta\alpha\beta$ motifs identified using BABMiner and PROMOTIF.** **(A)** Of the 31,731 PDB structures associated with F40 representative domains of ECOD version 291 [1], BABMiner and PROMOTIF [3] identified $\beta\alpha\beta$ motifs in 15,765 and 7,807 PDB structures, respectively. **(B)** Of the 48,801 $\beta\alpha\beta$ motifs found within the F40 representative domains of ECOD version 291, 92% (19,105/20,815) of $\beta\alpha\beta$ motifs identified by PROMOTIF are also identified by BABMiner (left). BABMiner identified 27,986 $\beta\alpha\beta$ motifs not identified by PROMOTIF, 57% (15,906/27,986) of which are open $\beta\alpha\beta$ motifs (right). BABMiner found $\beta\alpha\beta$ motifs in 182 additional X-groups, where the numbers of X-groups associated with each category are 487 (BM only), 296 (BM+PM), and 9 (PM only).

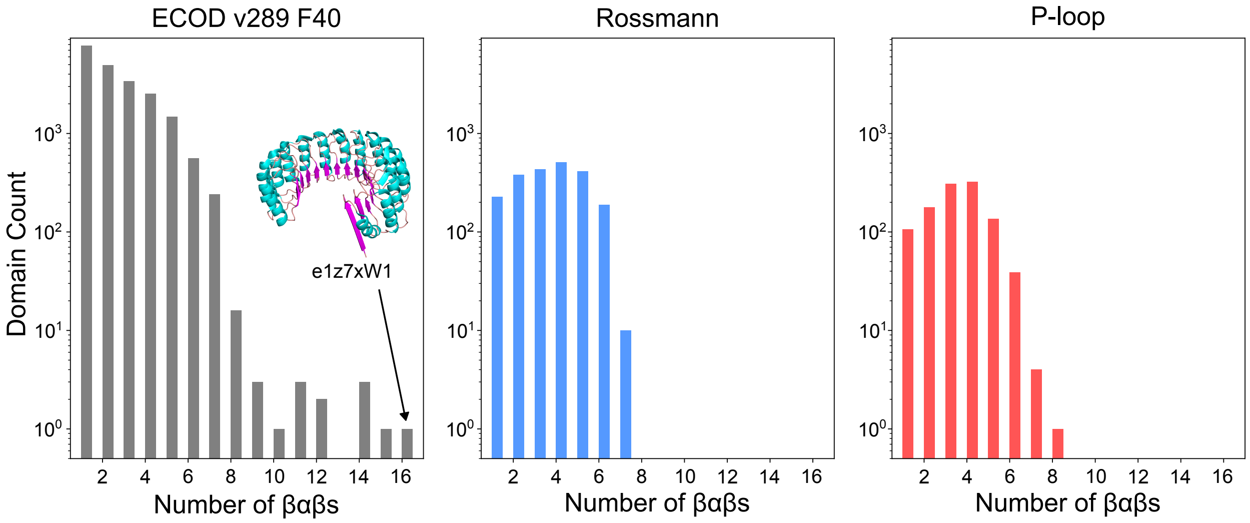

**Figure S3: Number of detected $\beta\alpha\beta$ motifs per domain across all ECOD F40, Rossmann, and P-loop representative domains.** Detected $\beta\alpha\beta$ motifs per domain in all ECOD F40 representative domains (left), representative Rossmann domains (middle), and representative P-loop domains (right). In total, 50,611 $\beta\alpha\beta$s were detected in 20,893 domains belonging to 831 X-groups. The domain with the most detected $\beta\alpha\beta$ motifs is a ribonuclease inhibitor protein, with a total of 16 motifs within a single domain (left panel, inset). Deviation from the idealized Rossmann and P-loop domains, which are expected to have 5 and 4 $\beta\alpha\beta$ motifs, respectively, relates to insertion or omission of secondary structure elements.

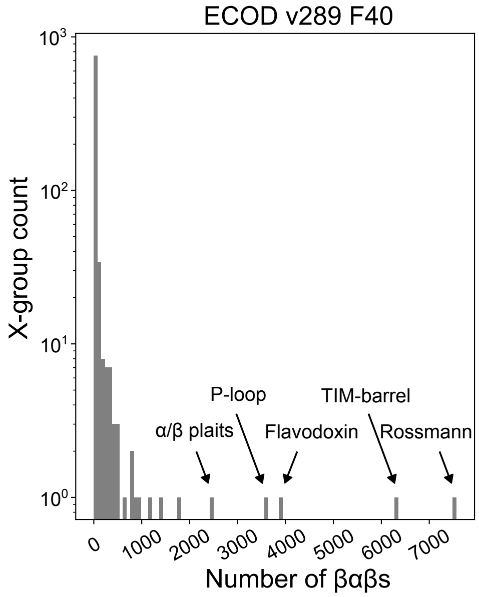

**Figure S4: The number of $\beta\alpha\beta$s per X-group.** In total, 50,611 $\beta\alpha\beta$s were detected in 20,893 domains belonging to 831 X-groups. 7,566 $\beta\alpha\beta$s were detected in 2,155 Rossmann domains and 3,610 $\beta\alpha\beta$s were detected in 1,090 P-loop domains.

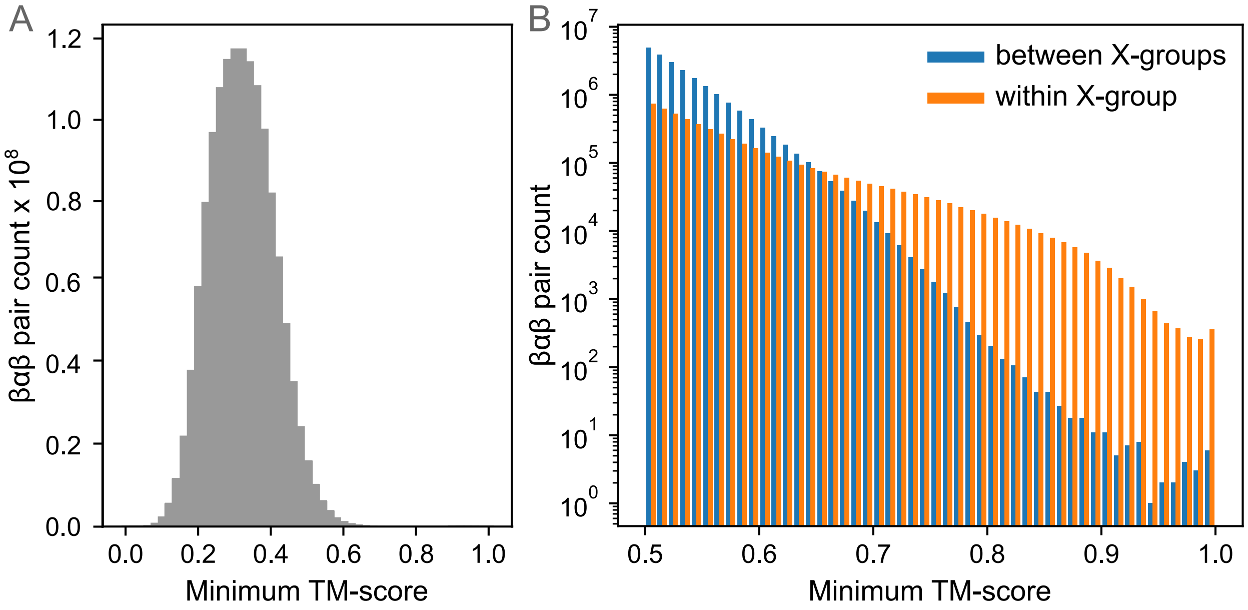

**Figure S5: Distribution of minimum TM-scores for all-vs-all $\beta\alpha\beta$ motif comparisons.** **(A)** Distribution of minimum TM-scores [10] from an all-vs-all comparison of the 50,611 $\beta\alpha\beta$ motifs detected within the representative domains of ECOD F40. Minimum TM-score refers to the minimum of the two TM-scores reported by TM-align [11] for a given structure pair, which relates to how TM-score is normalized by the length. Using the minimum TM-score thus penalizes $\beta\alpha\beta$ motifs that have significant differences in length (i.e., substructure similarity). **(B)** Distribution of minimum TM-scores from $\beta\alpha\beta$ pairs between (blue) and within (orange) X-groups. The plot shows 26,376,266 pairs with minimum TM-score $\geq 0.5$, of which 80.7% are between X-groups and 19.3% are within the same X-group. Above a minimum TM-score of 0.7, $\beta\alpha\beta$ pairs are more likely to be from within an X-group than between X-groups. $\beta\alpha\beta$ pairs with minimum TM-score above 0.7 were used for the analyses in the main text.

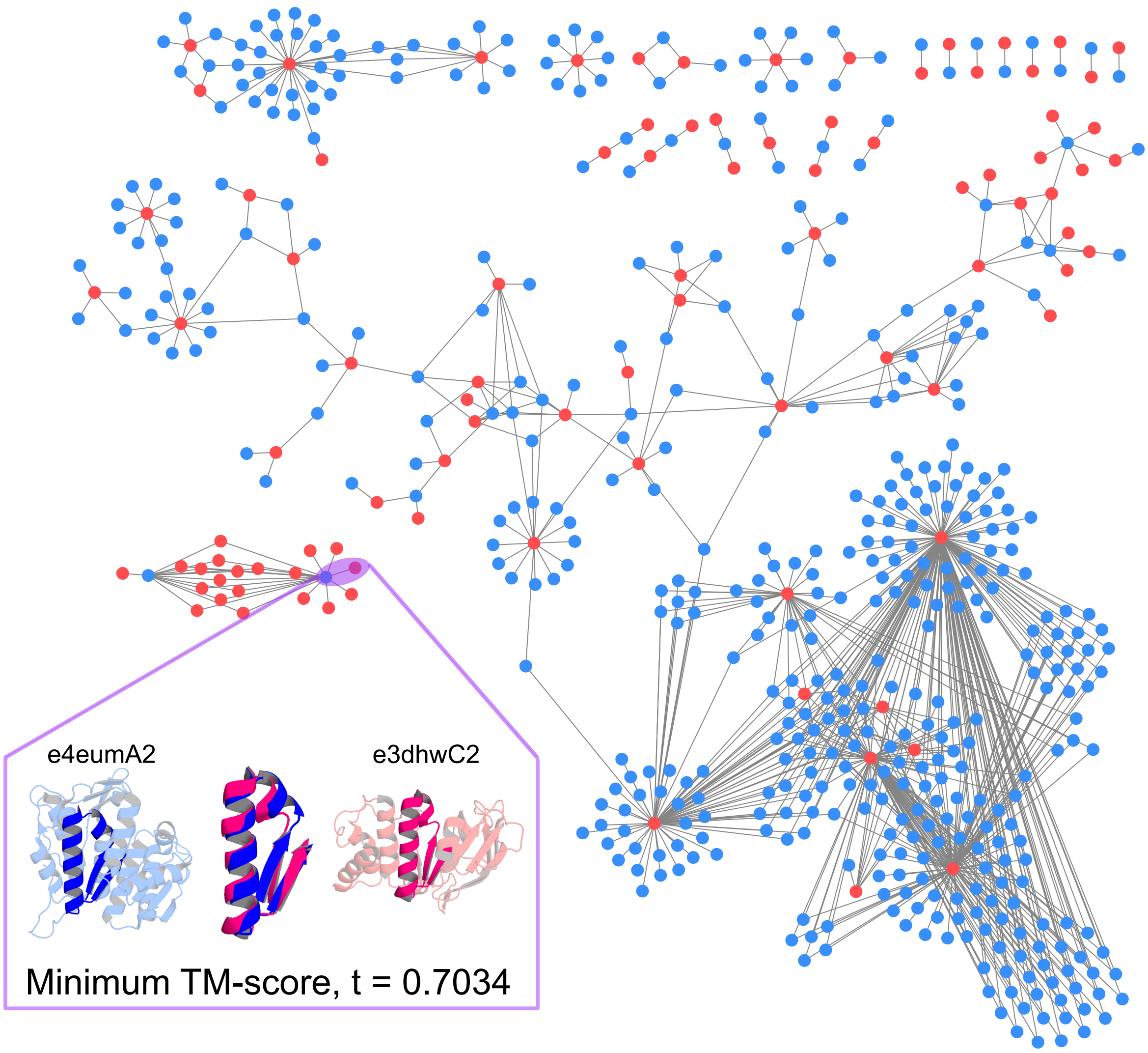

**Figure S6: Rossmann vs. P-loop $\beta\alpha\beta$ similarity network.** $\beta\alpha\beta$ similarity network between Rossmann $\beta\alpha\beta$s (blue, 463 $\beta\alpha\beta$s from 445 domains) and P-loop $\beta\alpha\beta$s (red, 93 $\beta\alpha\beta$s from 95 domains) with minimum TM-score cutoff = 0.7. An example TM-align result is shown in the bottom left.

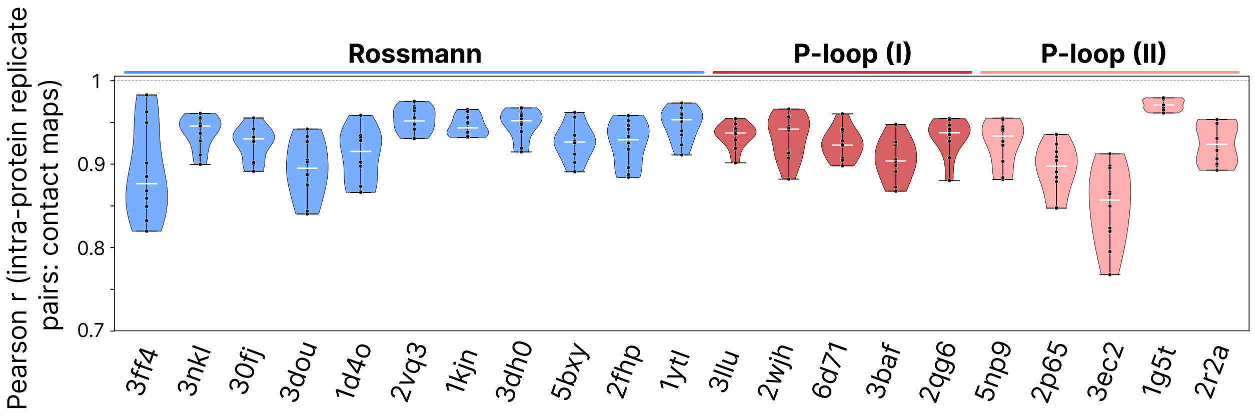

**Figure S7: Pairwise contact-map correlation across five MD replicates.** Each violin summarizes the 10 intra-protein pairwise Pearson correlations between the five 2 ns MD replicate contact maps for one protein. Black dots are individual replicate pairs; the white tick marks the median. Across all 21 proteins the median pairwise $r$ exceeds 0.85, with most distributions tightly above $r \approx 0.93$, suggesting that the per-replicate contact maps of a given protein are mutually consistent and that the per-protein average data is a faithful representative of the underlying replicate ensemble.

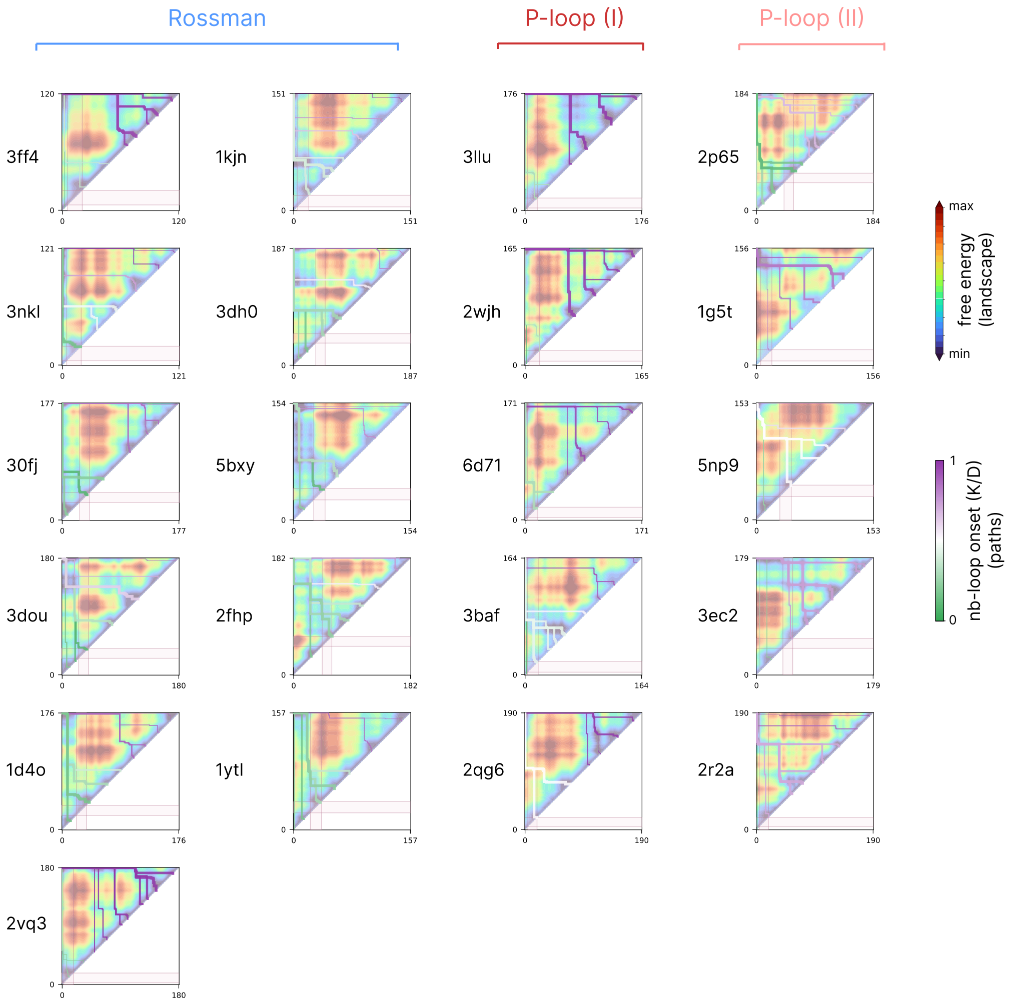

**Figure S8: SSA contig free energy landscapes with most thermodynamically favored paths for all 21 proteins.** Each panel shows the smoothed SSA [5] landscape ($-\log p$ over (start $i$, end $j$)) for one protein, overlaid with the top-$N$ folding pathways. Paths are colored by their $K/D$ at functional loop folding onset—the fraction of the domain folded at the path step the contig first fully covers the functional loop residues (blue = early, red = late). Path line-width decays from rank 1 (thickest, lowest barrier) to higher ranks; light pink $i/j$ bands indicate the functional loop residue range. Across the 21 proteins the path families consistently funnel toward a small set of attractors, with class-specific patterns of functional loop folding onset.

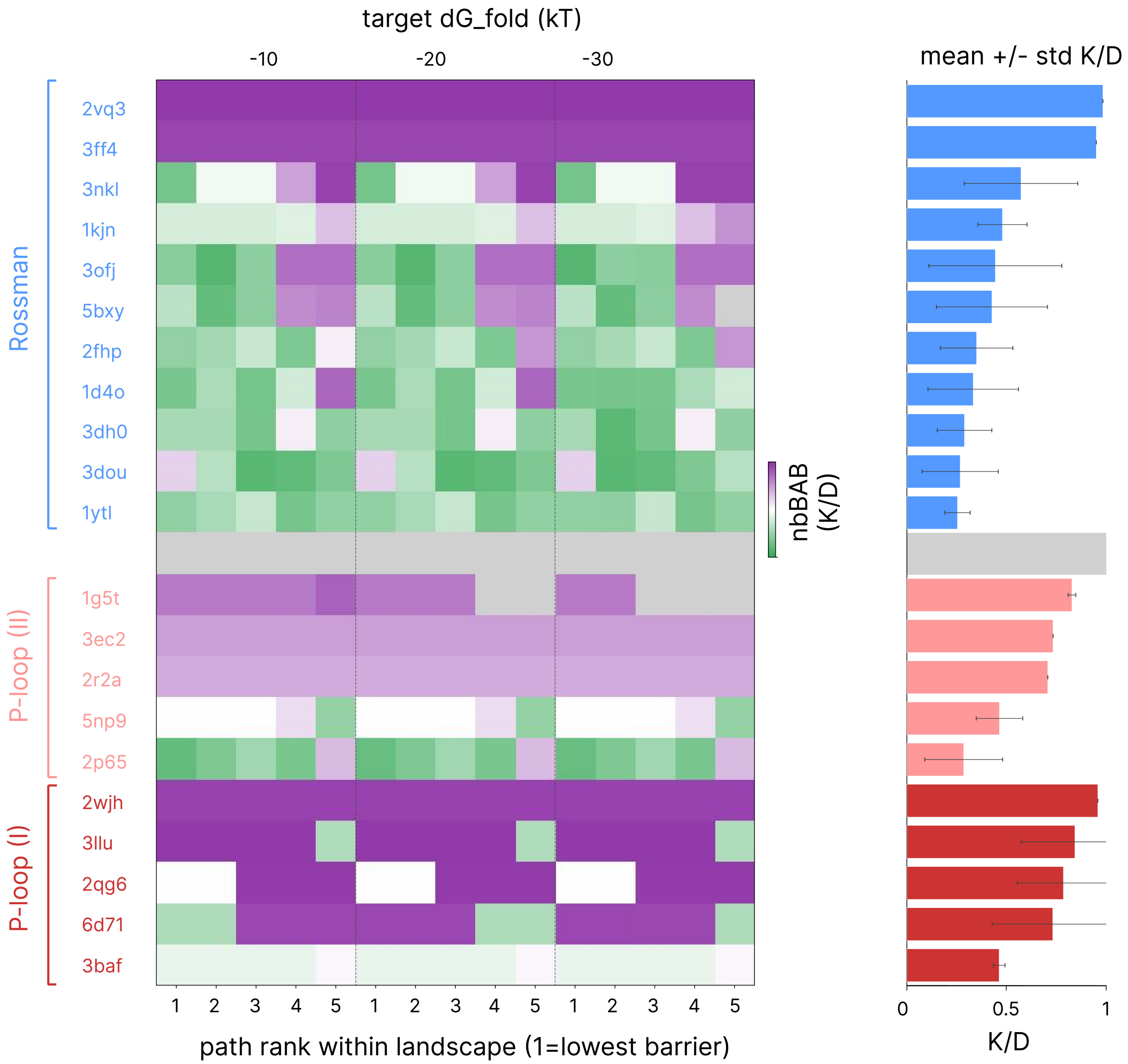

**Figure S9: Stability of SSA paths across hyperparameter target $\Delta G_{\text{fold}}$ values.** $\Delta G_{\text{fold}} = -10$ (manuscript default), $-20$, $-30$ kcal mol$^{-1}$—each with the top 5 paths, for 15 cells per row. Right: per-protein mean ± std $K/D$ across the 15 cells. The $K/D$ values, the rank 1 path identity, and the row ordering are essentially unchanged across the three $\Delta G_{\text{fold}}$ values, demonstrating that the functional loop fold ordering inferred from the top SSA paths is robust to the choice of folding free energy, suggesting that the qualitative conclusions of the main text do not depend on the exact hyperparameter setting.

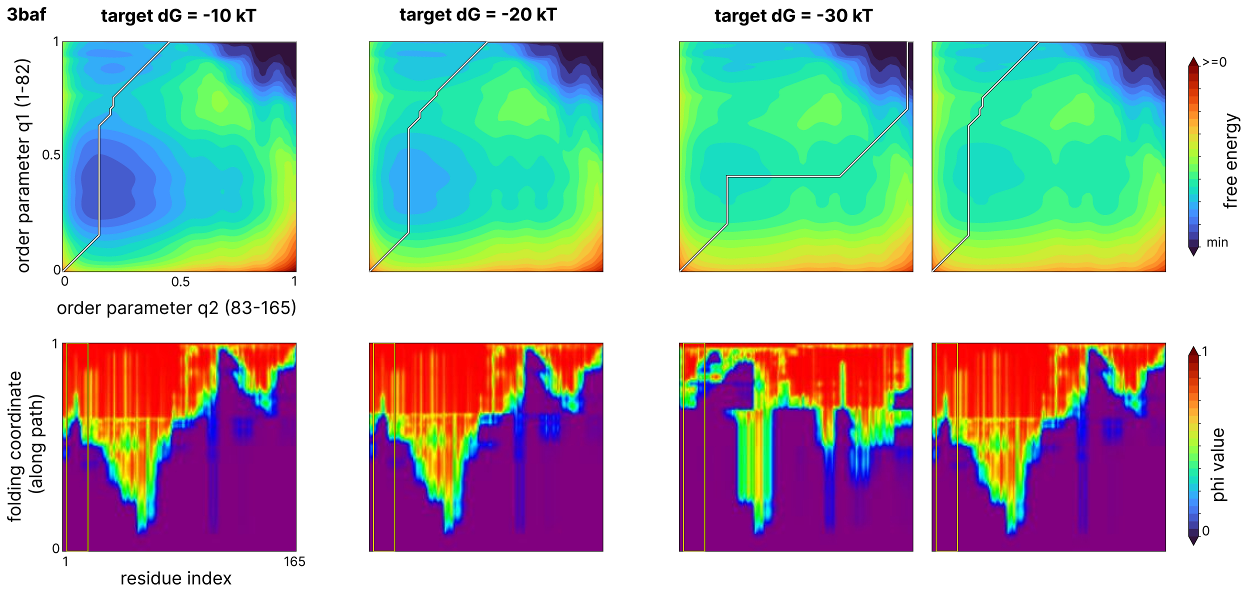

**Figure S10: Stability of WSME-L $\phi$-values along the optimal path across target $\Delta G_{\text{fold}}$ values (3baf).** Top: WSME-L [6] 2D free energy landscape at four conditions, with the folding path overlaid in gray—(1) $\Delta G_{\text{fold}} = -10$ (default), (2) $\Delta G_{\text{fold}} = -20$, (3) $\Delta G_{\text{fold}} = -30$, where the new optimal minimum free energy path shifts to a qualitatively different route (also viable in $\Delta G_{\text{fold}} = -10/-20$ landscapes), and (4) the $\Delta G_{\text{fold}} = -30$ landscape with the $\Delta G = -10$ optimal path (while disfavoured, it remains viable). Bottom: $\phi$-value heatmap along each of the four paths. The residue-level folding tendencies encoded by $\phi$-values are relatively robust to the choice of $\Delta G_{\text{fold}}$ provided.

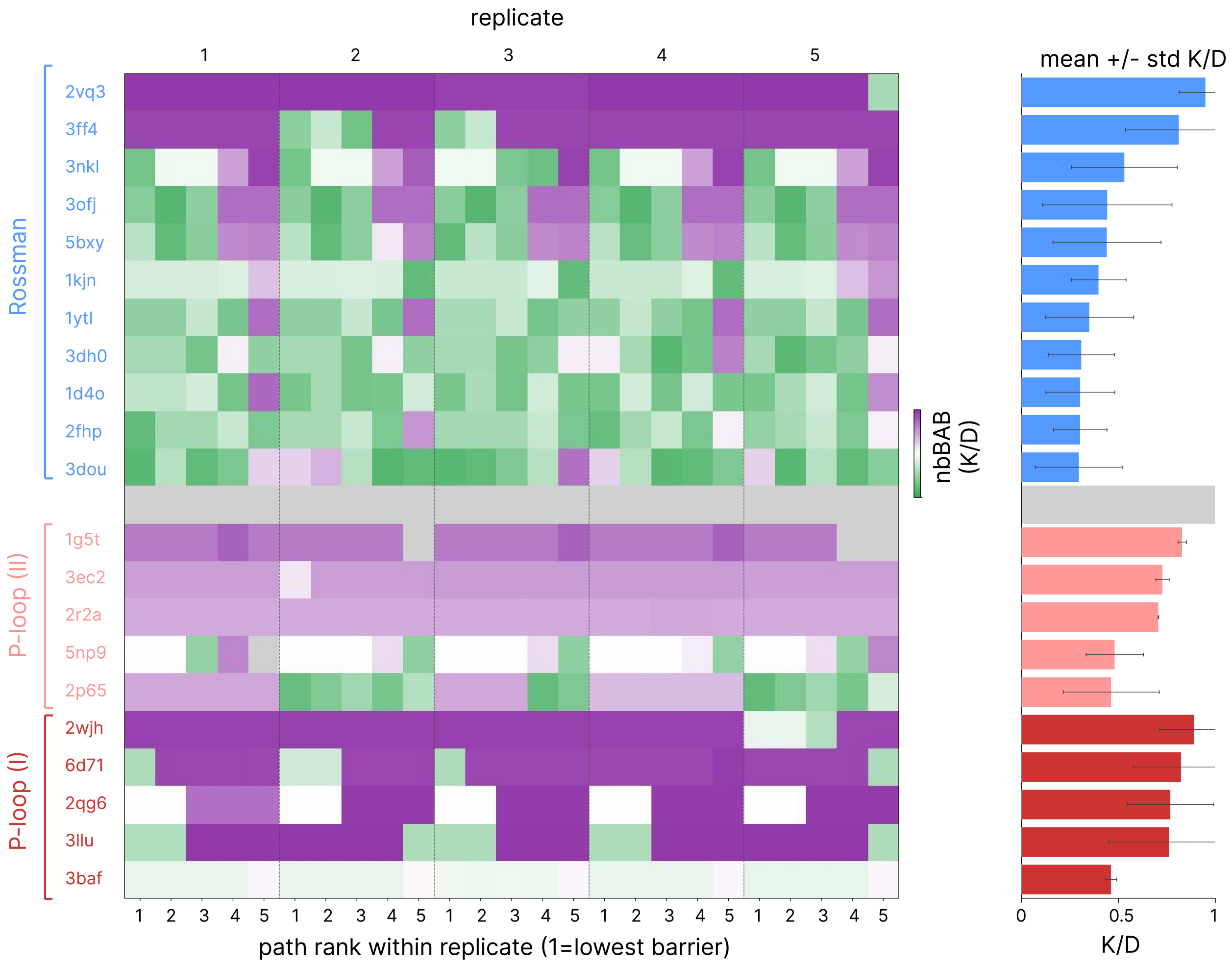

**Figure S11: Stability of SSA paths across replicates.** Left: heatmap of functional loop folding onset $K/D$ for the top 5 folding paths in each of the five replicates of every protein. The colormap encodes early (green) to late (purple) functional loop onset. Right: per-protein mean ± std of $K/D$ across all 25 cells, bar coloured by class. The narrow horizontal spread within each row in both summary panels shows that the functional loop fold ordering and the rank 1 path identity are stable across replicates: replicate-to-replicate variation in $K/D$ is relatively small relative to between-protein variation.

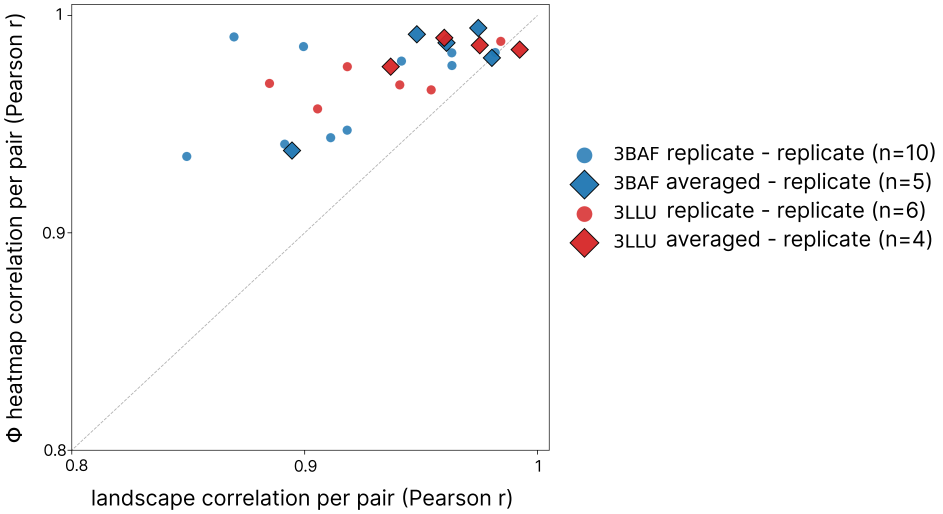

**Figure S12: Φ-value heatmap reproducibility versus landscape reproducibility across replicates, for 3baf and 3llu.** Every intra-protein pair from {averaged data reference, and replicates r0–r4} contributes one point: $x$ = Pearson $r$ of the flattened reference WSME-L free energy landscape $G(q_1, q_2)$, $y$ = Pearson $r$ of the flattened $\phi$-value heatmap (resampled to a common path step length). Diamonds mark pairs that include the averaged data reference; circles are replicate-vs-replicate pairs. The $\phi$-value heatmaps are consistently more reproducible than the underlying landscapes, demonstrating that even modest landscape perturbations preserve the residue-level folding ordering recovered by WSME-L.

**Algorithm 1** BuildTopologicalNetwork: minima and saddles of a free-energy surface

**Input:** raw landscape: triangular (SSA) $\mathbf{F}[j, i]$, or rectangular (WSME-L) $\mathbf{F}[q_1, q_2]$
**Output:** minima, maxima, saddles, and minimum–minimum connections

**Smooth:** Gaussian filter $\mathbf{F}$ ($\sigma = 1.5$)
**Hessian:** compute $F_{jj}, F_{ii}, F_{ji}$; set $\det H = F_{jj}F_{ii} - F_{ji}^2$ and $\operatorname{tr} H = F_{jj} + F_{ii}$
**Critical points:** in each $\det H > 0$ patch take one extremum — a minimum if $\operatorname{tr} H > 0$, else a maximum
**Basins:** watershed the surface from the minima (one Voronoi-like basin each)
**Saddles:** for each adjacent basin pair, take the lowest boundary pass $\min \max(\mathbf{F}[c], \mathbf{F}[c'])$
**Refine:** relax a string between each minima pair along $-\nabla\mathbf{F}$ perpendicular to its tangent; the highest point is the saddle
**return** network of minima and saddles

**Algorithm 2** MonotoneMinimaxPath: lowest-barrier monotone path

**Input:** surface $\mathbf{F}$; move set $\mathcal{M}$; sources $\mathcal{S}$; target $t$
**Output:** path minimising the bottleneck energy $\max_c \mathbf{F}[c]$

sweep cells in increasing reaction coordinate ($K = j - i + 1$ for SSA; $q_1 + q_2$ for WSME-L)
a source starts a path at its own energy; any other cell inherits $\max(\mathbf{F}[c],$ best predecessor$)$
break ties toward the lower cumulative energy, so the path hugs the valley floor
**return** backtrack from $t$ to its start

**Algorithm 3** SSA I–TN pathsfinder: one path per ingression valley

**Input:** protein landscape; native corner $t = (D - 1,\ 0)$
**Output:** ranked folding paths

load and smooth the C1 landscape
**Ingressions:** threshold at the 15th percentile, split into valleys by watershed, keep valleys that are large and touch the $K = 1$ diagonal
**Network:** BuildTopologicalNetwork (used only to label saddles)
allowed moves $\mathcal{M} = \{\, j \to j{+}1,\ i \to i{-}1 \,\}$ (single tile flip; $K$ increases)
**for** each valley **do**
    start from the deep core of the valley
    path ← MonotoneMinimaxPath: diagonal → valley, then valley → fully folded
    record the barrier and label its bottleneck with the nearest saddle
**end for**
**return** paths ranked by barrier

**Algorithm 4** WSME-L TN PATHSFINDER: one monotone path through $U$ and $N$

**Input:** $\mathbf{F}[q_1, q_2]$
**Output:** monotone path with its transition state

load and smooth the landscape
**Network:** BUILDTOPOLOGICALNETWORK
pick anchors: $U$ = lowest minimum in the unfolded corner, $N$ = lowest in the native corner
allowed moves $\mathcal{M} = \{\, q_1 \to q_1{+}1,\ q_2 \to q_2{+}1 \,\}$ (non-decreasing)
path ← MONOTONEMINIMAXPATH through $(0,0) \to U \to N \to (n_1{-}1,\ n_2{-}1)$
**TS** ← highest-energy cell on the $U \to N$ leg; barrier = $\mathbf{F}[\mathrm{TS}] - \mathbf{F}[U]$
**return** path, TS, barrier

**Figure S13: Pseudocode: procedure for identifying folding paths in WSME free energy landscapes.** Four algorithms used in the I-TN / TN pathfinding pipeline. BUILDTOPOLOGICALNETWORK extracts basins and refined saddles from a smoothed free-energy surface; MONOTONEMINIMAXPATH finds a lowest bottleneck monotone path; SSA I–TN PATHSFINDER and WSME-L TN PATHSFINDER assemble folding routes for SSA and WSME-L landscapes, respectively.

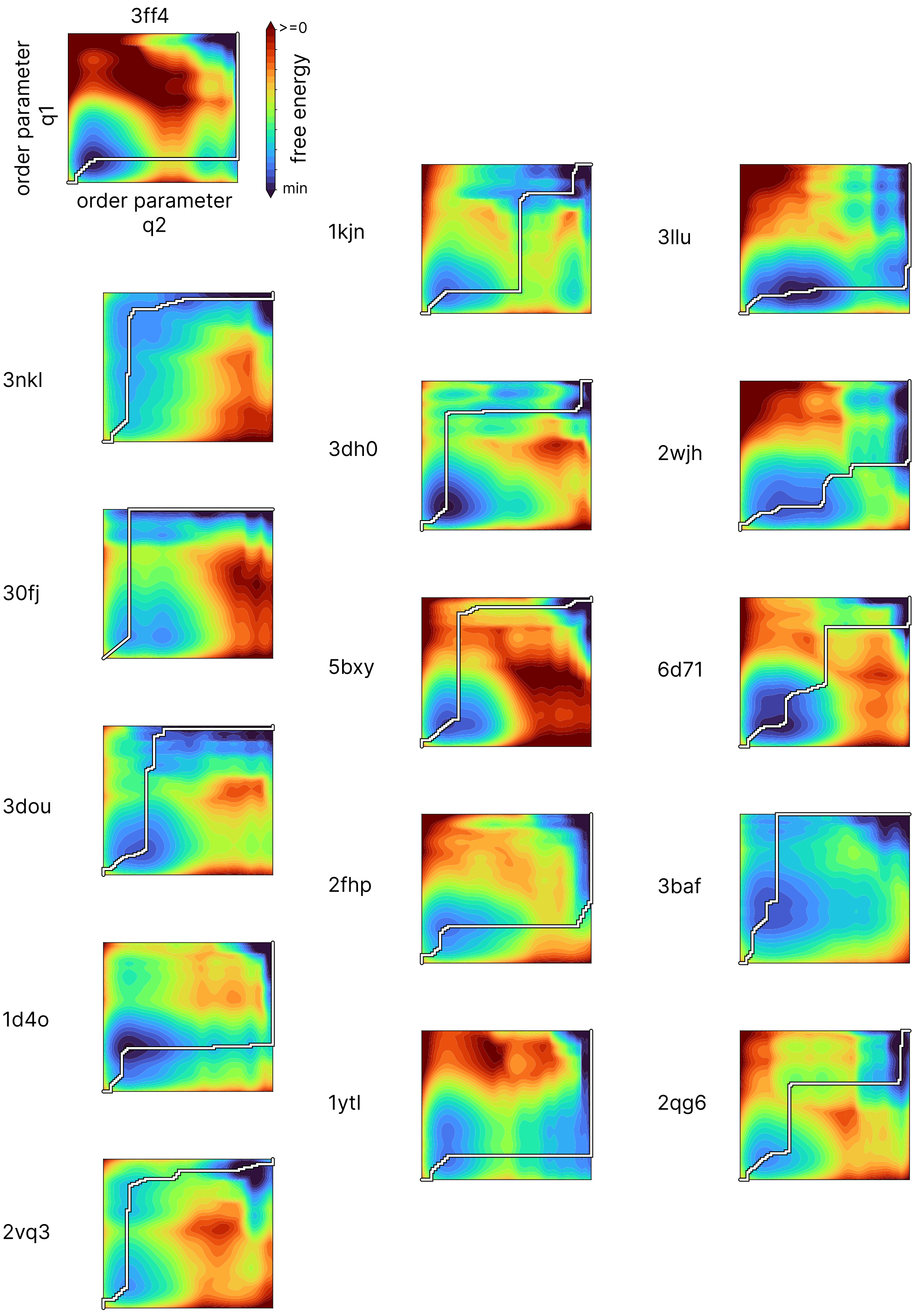

**Figure S14: WSME-L 2D landscapes for all 21 proteins.** Folding trajectories are shown in white.

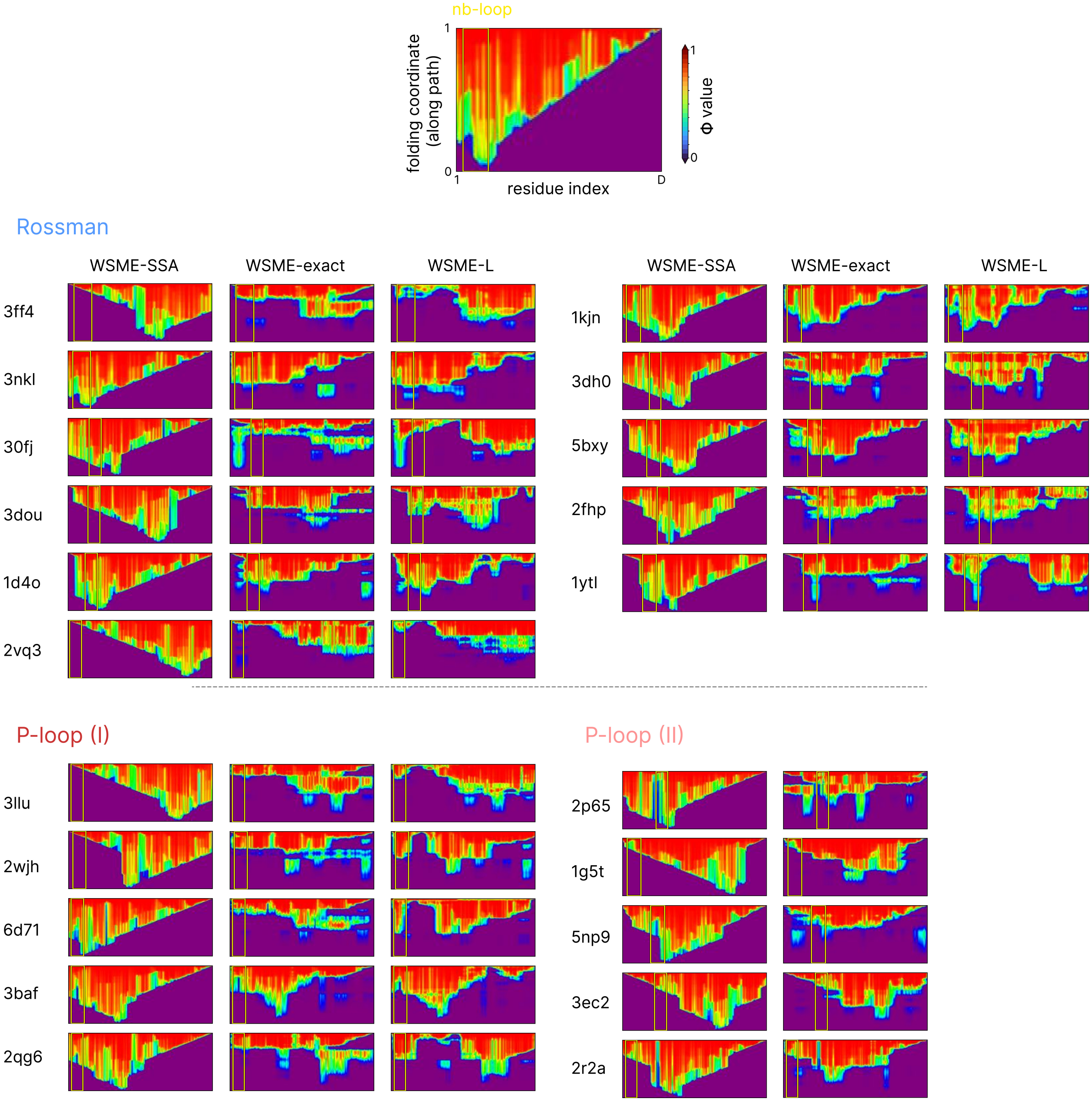

**Figure S15: Per-protein residue-resolved Φ heatmaps from SSA, WSME-exact and WSME-L.** The vertical axis is the method's native folding coordinate (along folding trajectories in SSA and WSME-L, and the model order parameter $q$ for WSME-exact); residue index runs along the horizontal axis. The yellow rectangle marks the folding loop. White overlays on SSA panels trace the rank-1 folding contig. WSME-L was not applied to type II P-loops.

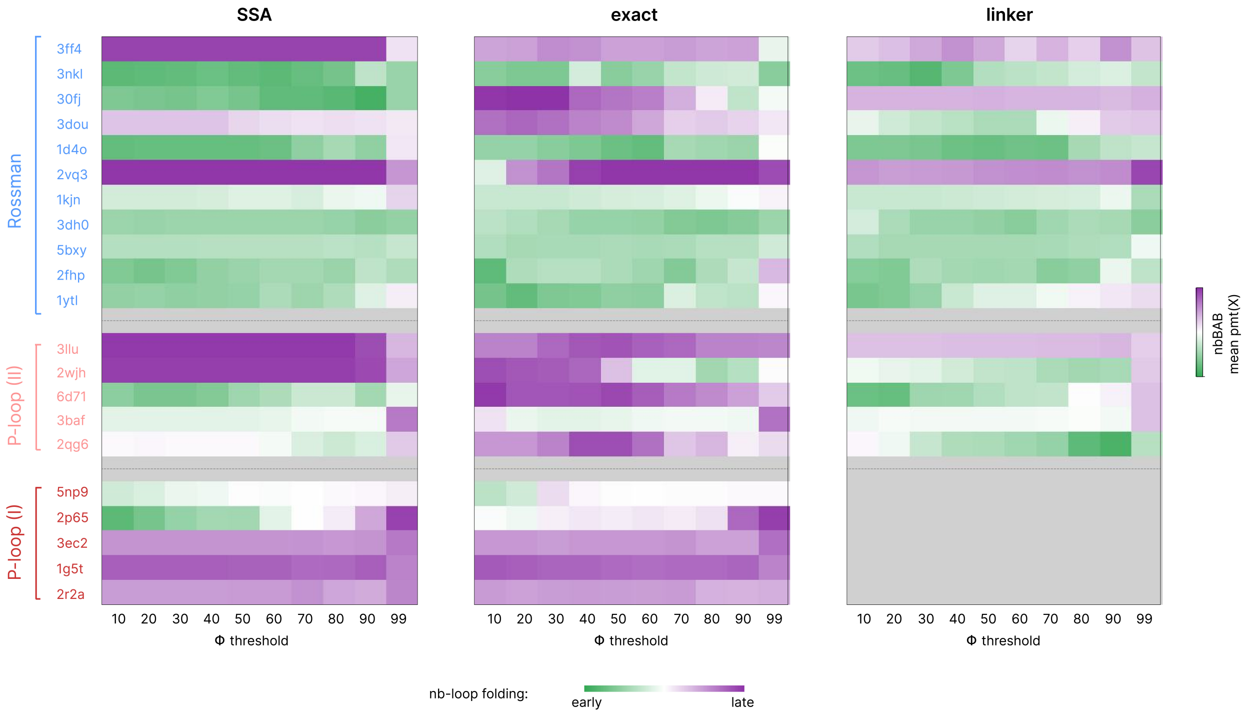

**Figure S16: Functional loop early-folding score across $\phi$-value thresholds, by method.** For each method (SSA, WSME-exact, and WSME-L) every cell shows the percentile rank of mean $t_X$ over the functional loop residues against the distribution of mean $p_{\text{mt}}(X)$ across all same-length contigs (sliding windows) in the same protein. Ten $\phi$-value thresholds $X \in \{0.1, 0.2, \ldots, 0.9, 0.99\}$. Green (low percentile) = the functional loop folds earlier than most equally long stretches in the same protein; purple = later. WSME-L was not applied to type II P-loops.

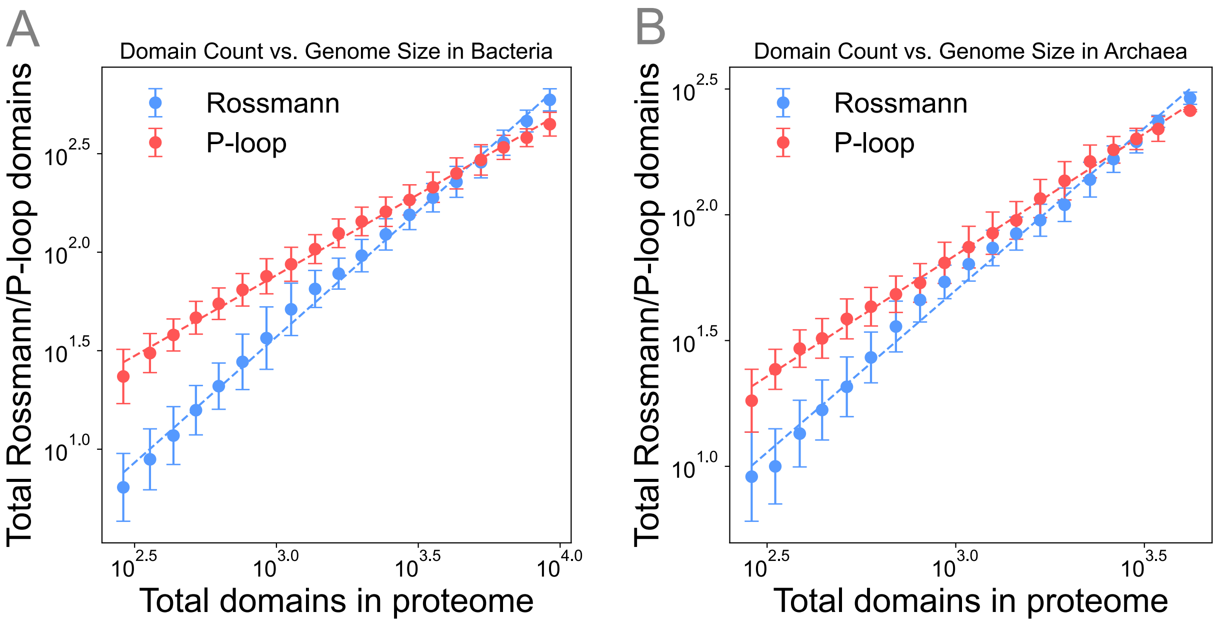

**Figure S17: Proteome-wide scaling of Rossmann and P-loop domains in Bacteria and Archaea.** **(A)** Bacterial genomes. **(B)** Archaeal genomes. The number of Rossmann (blue) and P-loop (red) domains is plotted against total proteome size across representative genomes from GTDB [7]. Both axes are on a $\log_{10}$ scale. Points show quality score-weighted means $\pm$ standard deviations within equal-width proteome size bins; dashed lines are power-law fits. P-loop domains predominate in smaller proteomes, whereas Rossmann domains scale more steeply, diversifying disproportionately in larger proteomes.

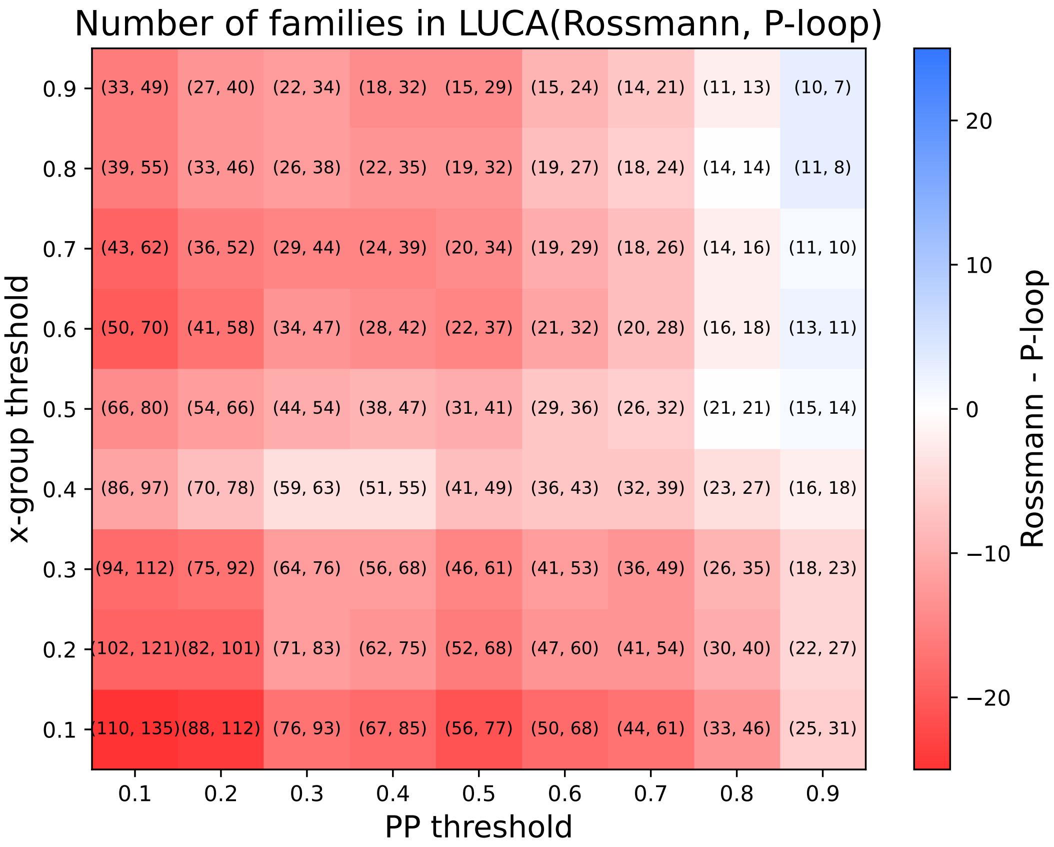

**Figure S18: LUCA reconstruction of Rossmann and P-loop gene families across COG probability and domain fraction thresholds.** Heatmap showing the number of Rossmann and P-loop gene families inferred to have been present in LUCA, evaluated across combinations of two thresholds. The $x$-axis represents the minimum posterior probability (PP) threshold from Moody *et al.* [4] for LUCA presence; COGs with PP $\geq$ PP threshold are counted as LUCA-present. The $y$-axis represents the minimum fraction of sequences within a COG annotated as Rossmann or P-loop domains by `hmmsearch` (see Methods); COGs with X-group fraction $\geq$ X-group threshold are classified as Rossmann or P-loop COGs. Each cell shows the number of qualifying COGs as (Rossmann, P-loop), and cell color indicates their difference (Rossmann − P-loop); blue indicates more Rossmann, red indicates more P-loop.

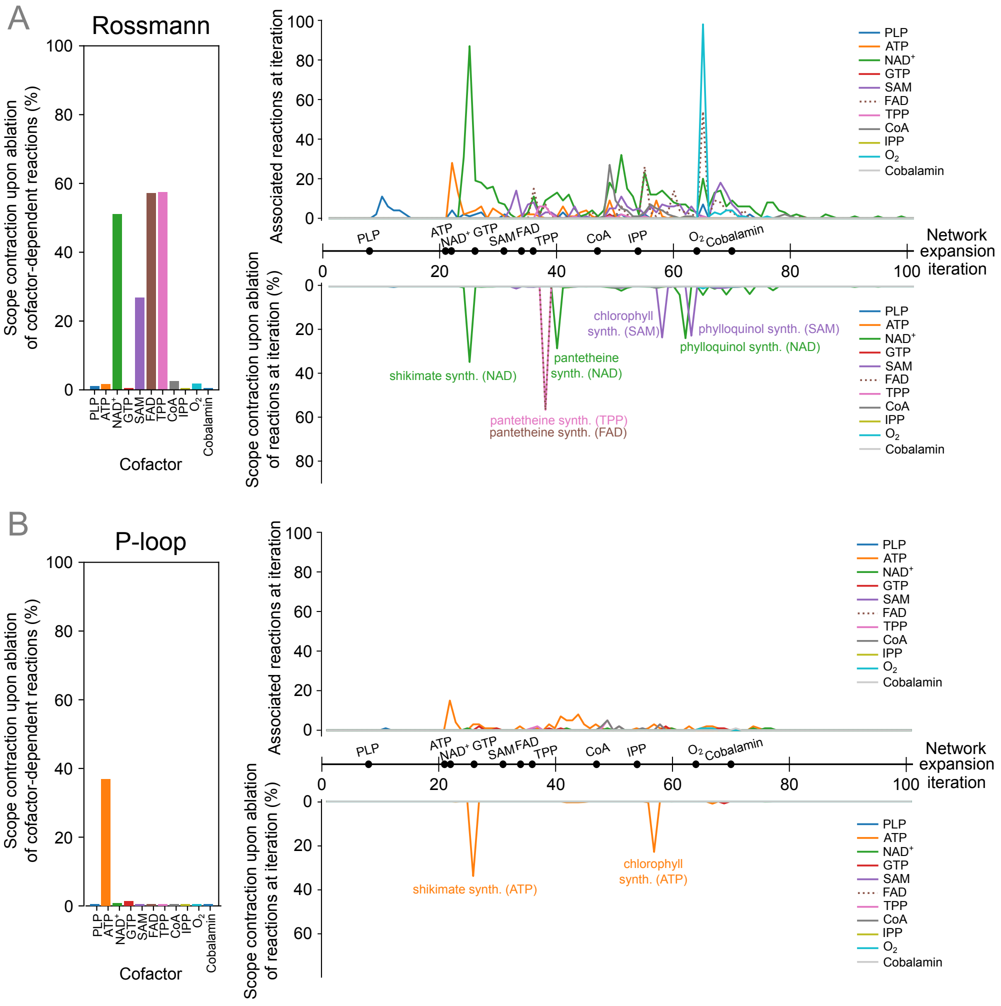

**Figure S19: Ablation of Rossmann- and P-loop-dependent reactions requiring each of the cofactor classes.** Significant scope contraction is observed when ablating Rossmann-dependent reactions requiring thiamine pyrophosphate (TPP, 57%), flavin adenine dinucleotide (FAD, 57%) and nicotinamide adenine dinucleotide ($NAD^+$, 51%). The trajectories on the right show the variations of main text Figure 2D, where the top panel shows the number of Rossmann- or P-loop-associated reactions that depend on each cofactor ($y$-axis) discovered at each iteration of the network expansion ($x$-axis), and the bottom panel shows the extent of scope contraction ($y$-axis, in % of full scope) upon ablation of cofactor-dependent reactions for which Rossmann or P-loop is required (present in all enzyme orthogroups) at that iteration.

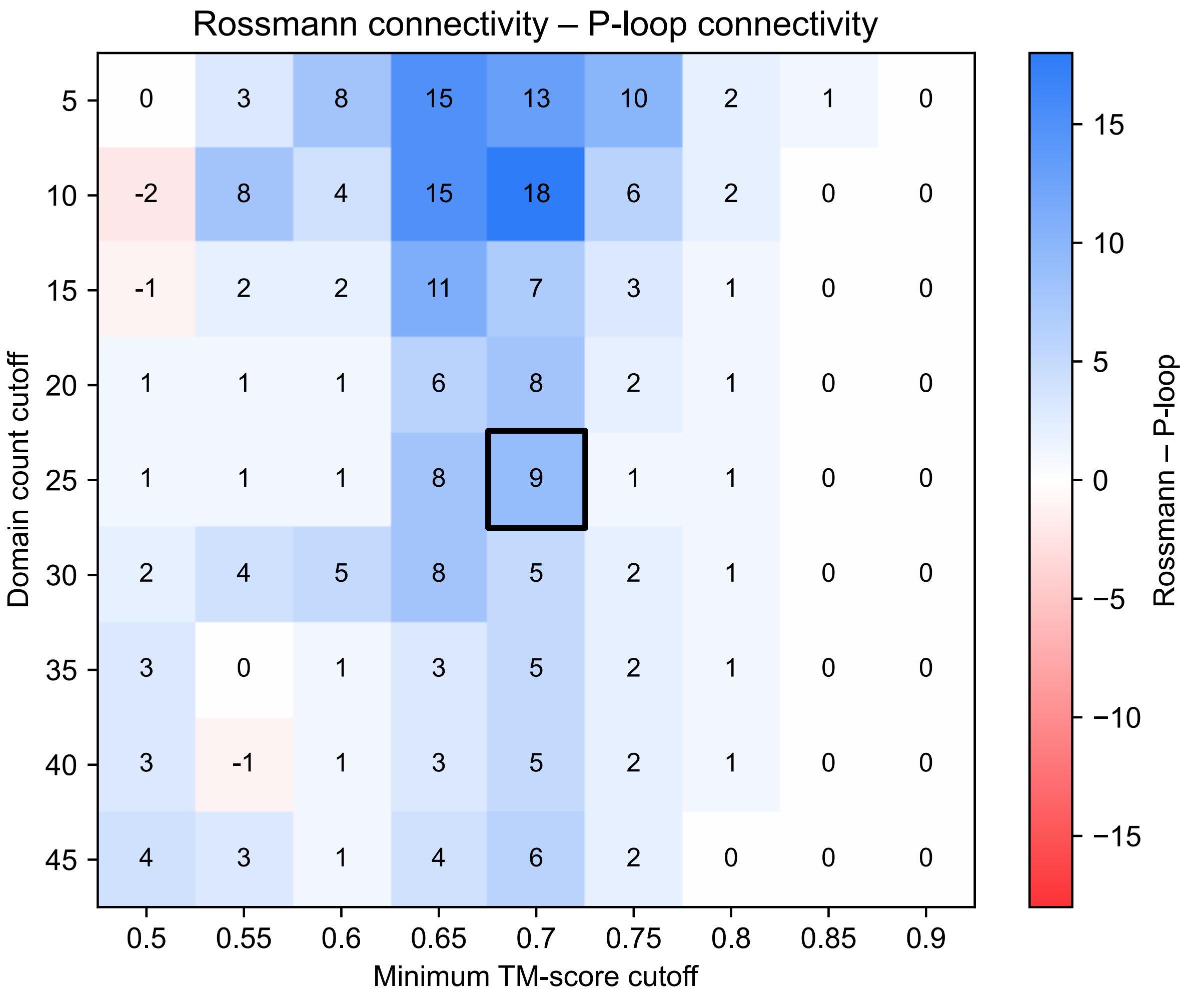

**Figure S20: Rossmann node degree exceeds P-loop across most TM-score and domain count cutoffs.** Heatmap of the difference between Rossmann and P-loop node degrees (Rossmann node degree – P-loop node degree) at various minimum TM-score and domain count cutoffs. The black box within the plot indicates the set of cutoffs used to generate the network in main text Figure 3A.

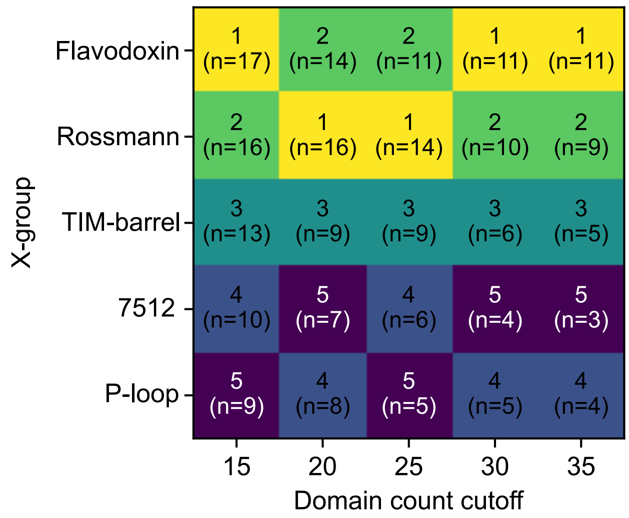

**Figure S21: X-groups with the highest node degrees.** Top 5 highest node degree X-groups at a minimum TM-score cutoff of 0.7 and a domain count cutoff ranging from 15–35. Rossmann and flavodoxin (X-group 2007) are consistently the most connected fold across all domain count cutoffs. Node degrees are indicated in parentheses.

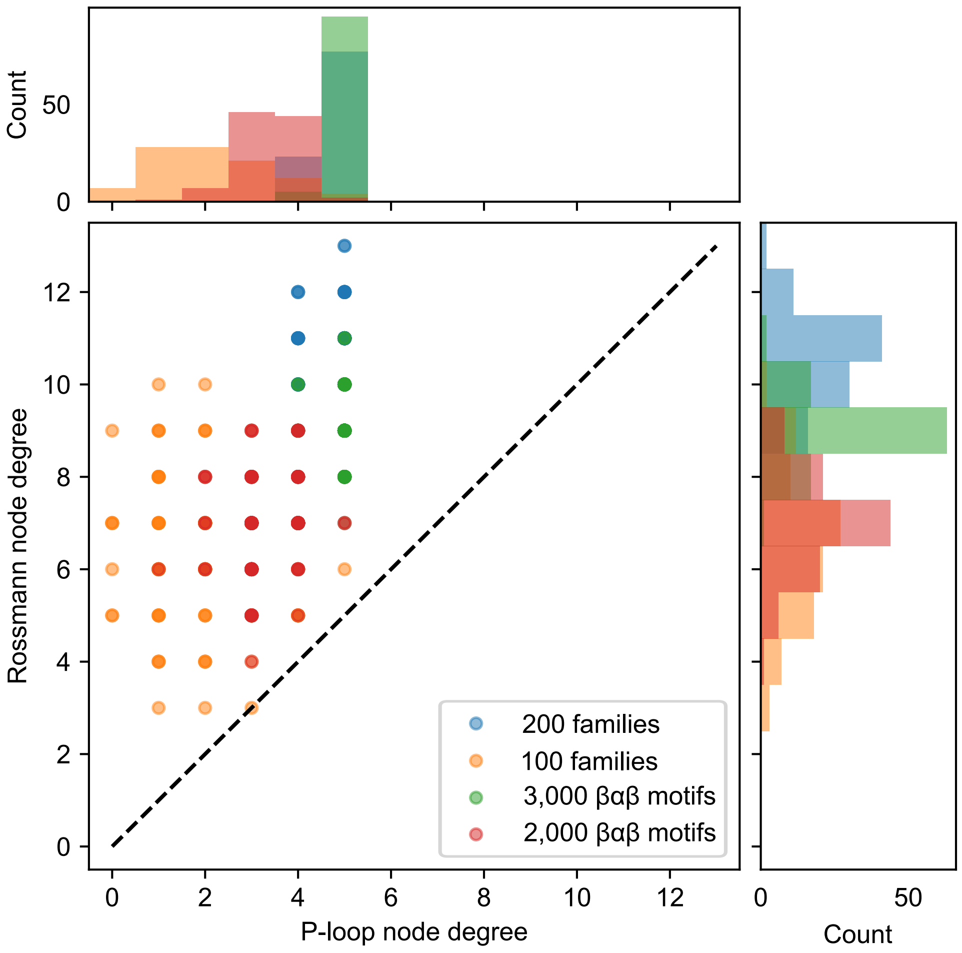

**Figure S22: Dominance of Rossmann over P-loop in the fold-lineage network is robust to subsampling.** Rossmann and P-loop node degrees within the fold-lineage network with subsampling. Subsampling at the level of families ($n$=200, 100) and $\beta\alpha\beta$s ($n$=3,000, 2,000) does not affect the dominance of Rossmann over P-loop. For each category, subsamples were taken randomly 100 times.

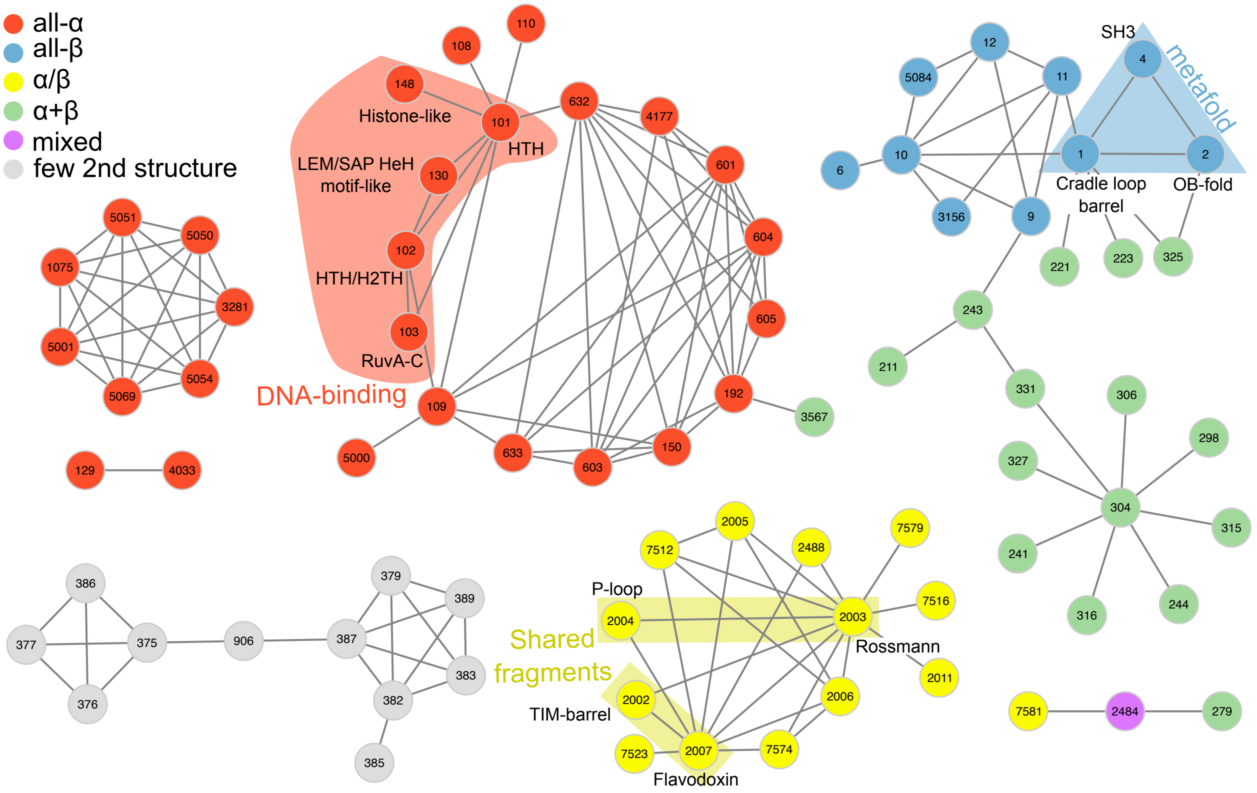

**Figure S23: Fold space feature captured by CLSS-sub** ($d$=0.3, $n$=25)**.** The network was generated using CLSS-sub [9]. The cutoff values used are those applied in Fig. 3B. Numbers within each node indicate X-group IDs in ECOD v289, and colors denote classification by architecture. At this cutoff, architectures are clearly partitioned, and the grouping captures metafolds for which structural transitions have been experimentally observed with few mutations, folds sharing fragments with high structural similarity, and characteristic features such as DNA-binding function.

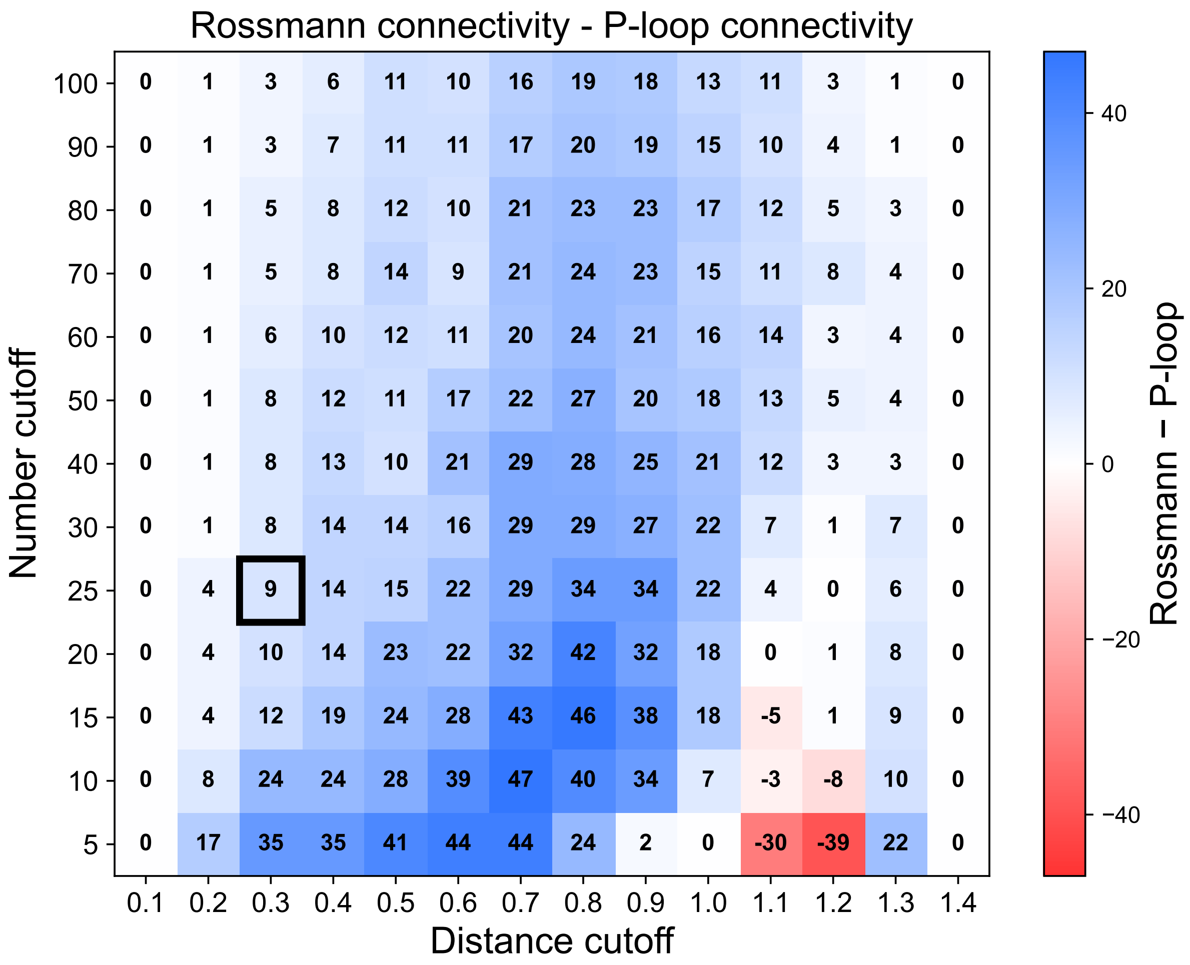

**Figure S24: Heatmap of each distance and number cutoff at the CLSS-sub model.** Heatmap of differences (Rossmann node degree – P-loop node degree) at various minimum distance cutoff and domain count cutoffs for the CLSS-sub model. The black box within the plot indicates the set of cutoffs used to generate the network in main text Figure 3B.

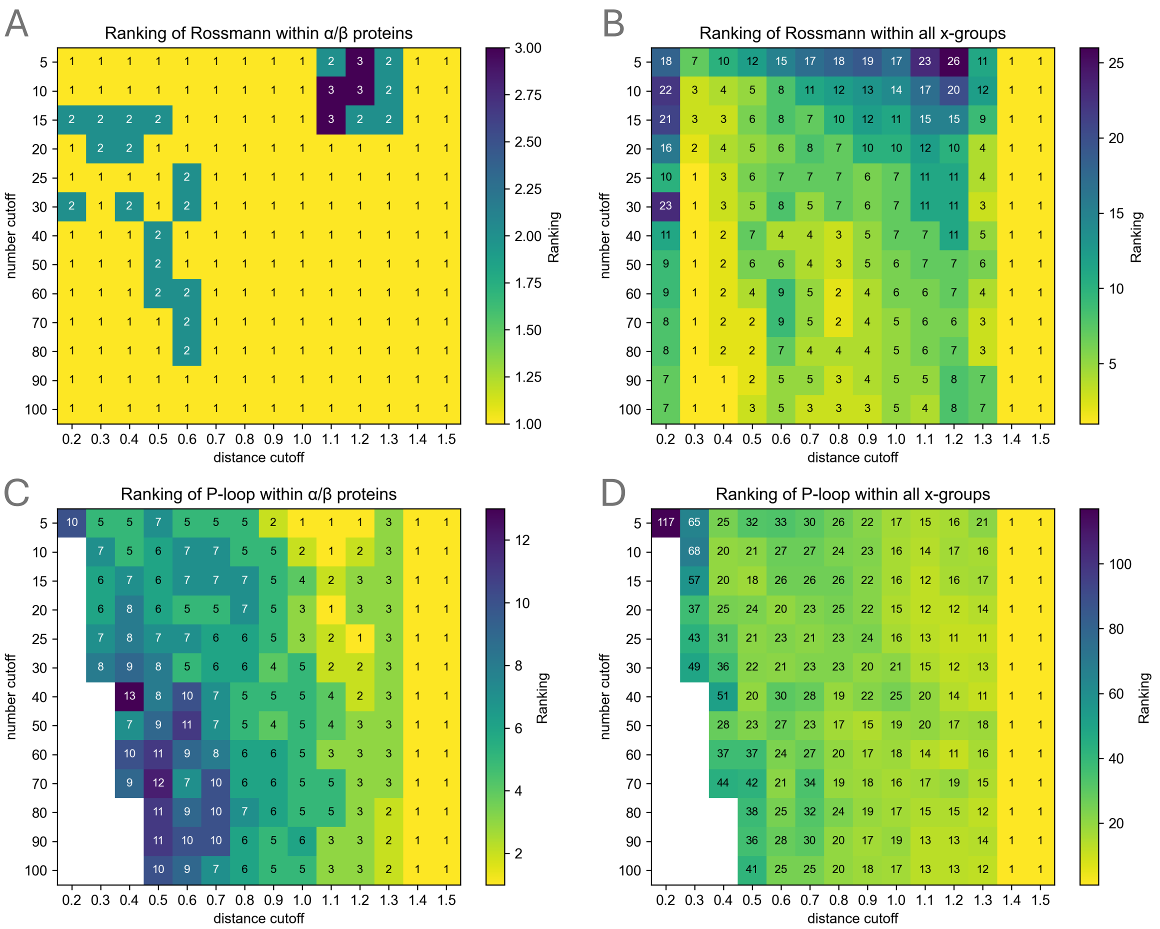

**Figure S25: Ranking of number of connections with all protein families among $\alpha/\beta$ proteins.** Rankings of the number of connections for Rossmann and P-loop folds at each cutoff threshold in CLSS-sub. **(A)** Ranking of Rossmann within $\alpha/\beta$ proteins. **(B)** Ranking of Rossmann across all protein folds. **(C)** Ranking of P-loop within $\alpha/\beta$ proteins. **(D)** Ranking of P-loop across all protein folds.

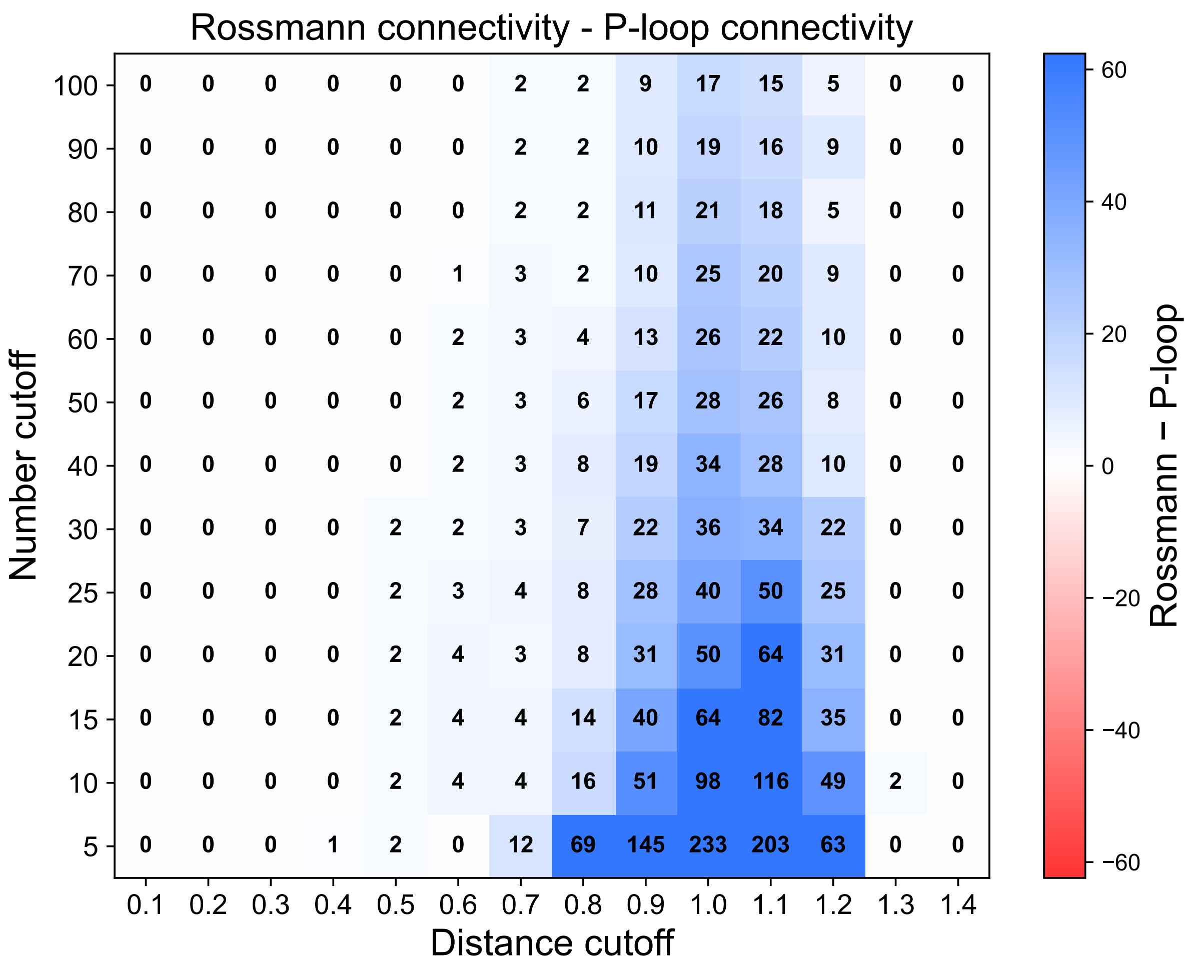

**Figure S26: Heatmap of each distance and number cutoff at the CLSS-full model.** Heatmap of differences (Rossmann node degree − P-loop node degree) at various minimum distance cutoff and domain count cutoffs for the CLSS-full model.

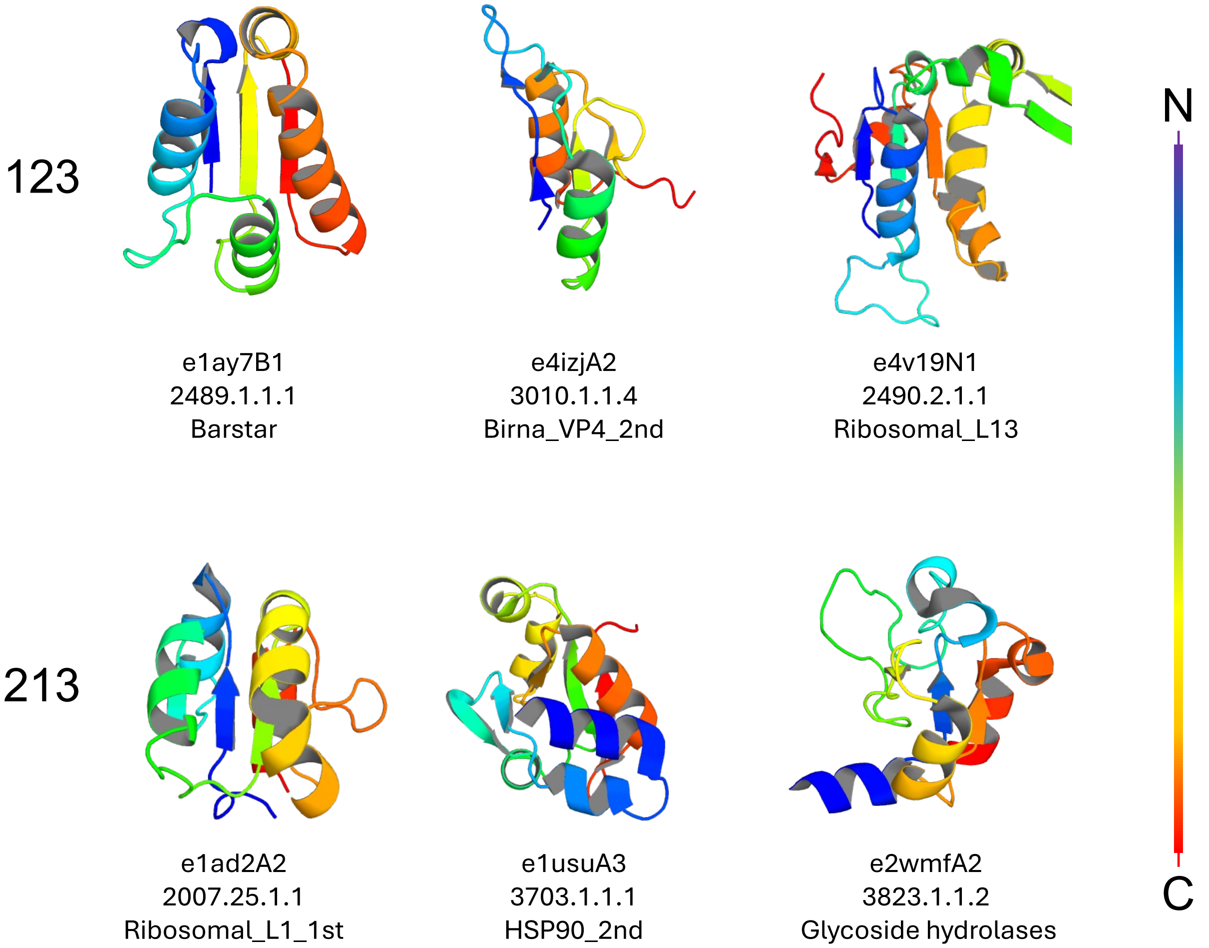

**Figure S27: Examples of 321 and 213 Rossmannoid proteins.** **(A)** Proteins with strand order 1–2–3. **(B)** Proteins with strand order 2–1–3. Structures are colored in rainbow from blue (N-terminus) to red (C-terminus). Only the ECOD domain region is shown. Labels below each structure indicate the ECOD domain ID, ECOD f-id, and ECOD f-name, respectively.

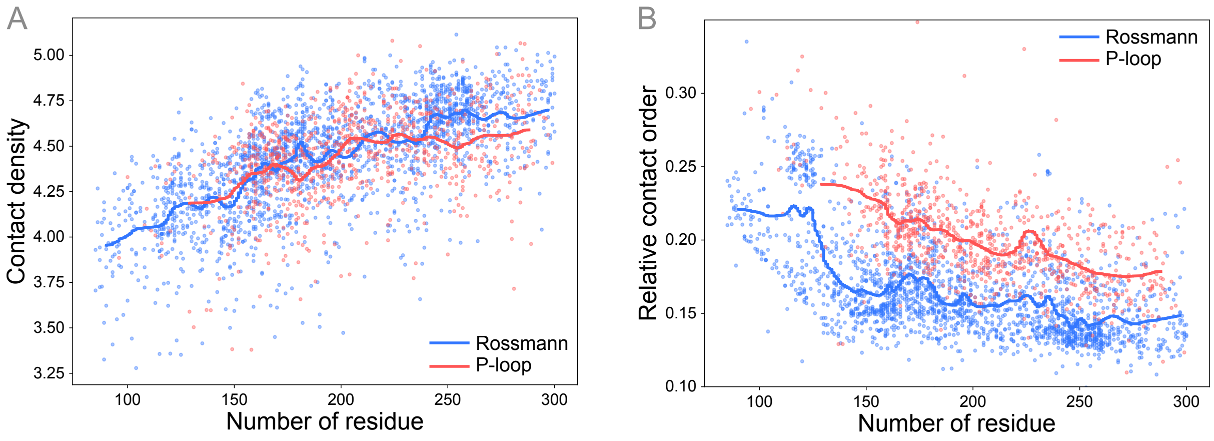

**Figure S28: Contact density and contact order of Rossmann and P-loop domains.** **(A)** Calculated contact density [2] and **(B)** relative contact order [8] for representative Rossmann and P-loop domains from ECOD (see Methods). In both panels, each point represents an individual domain, and the solid lines indicate moving averages.

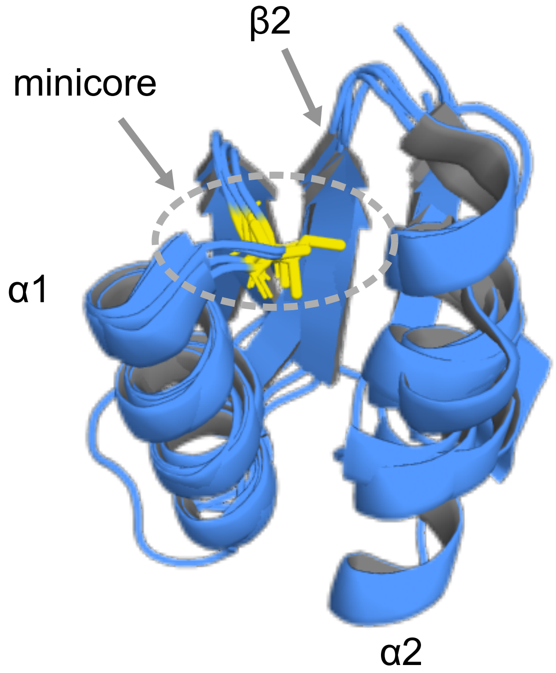

**Figure S29: Mini-core of representative Rossmann.** Structural superposition of the N-terminal strands 1–2–3 of canonical Rossmann folds (ECOD domain IDs: e1lssA1, e1nvtA1, e5k1sA1, and e5coaA1). Residues highlighted with yellow sticks represent the minicore, which faces inward toward $\alpha 1$, $\alpha 2$, and $\beta 2$, illustrating its contribution to loop formation.

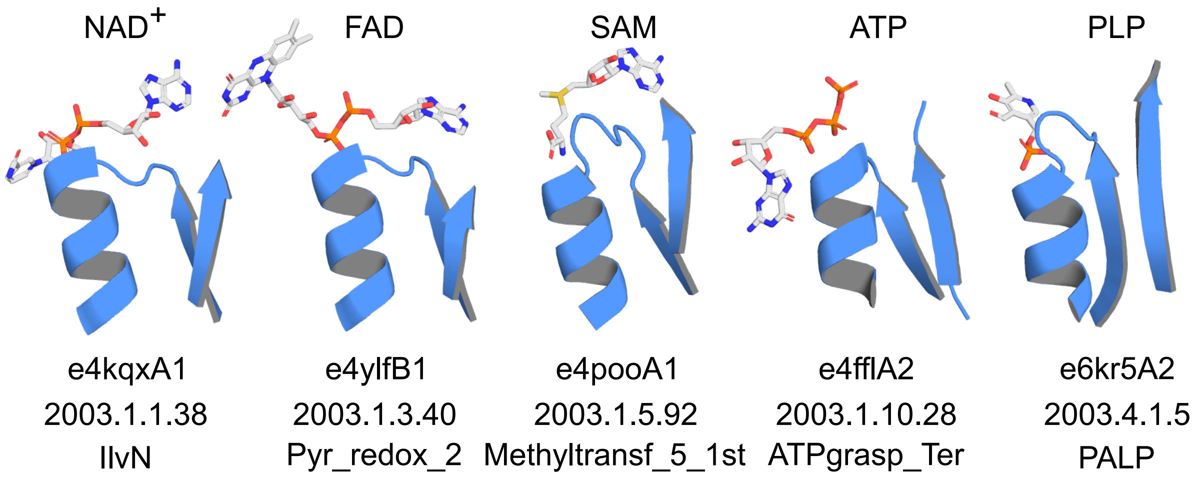

**Figure S30: The Rossmann binding motif is a flat conserved surface.** Representative cofactor-binding sites of Rossmann domains for $NAD^+$, FAD, SAM, ATP, and PLP are shown. Labels below each structure indicate the ECOD domain ID, ECOD f-ID, and ECOD f-name, respectively.

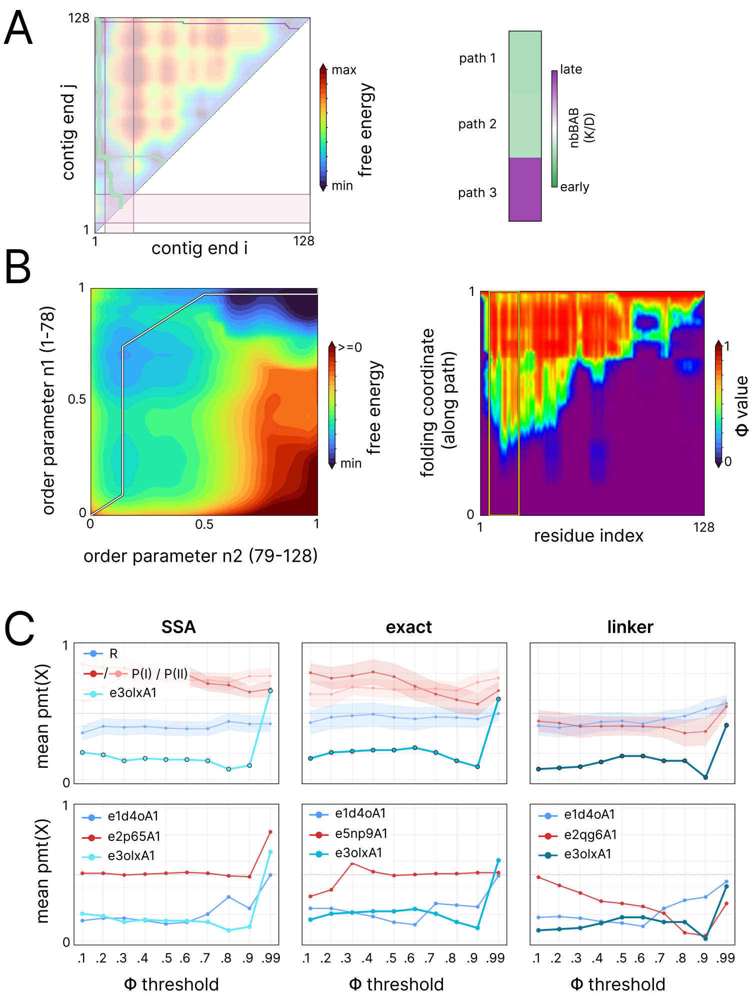

**Figure S31: Folding paths of flavodoxin.** Folding analyses were performed using e3olxA1. **(A)** Left, contig-based free energy landscape for e3olxA1 estimated via WSME-SSA. Right, heatmap of SSA functional loop onset $K/D$ for the three lowest-action paths. **(B)** Left, contig-based free energy landscape for e3olxA1 estimated via WSME-L. Right, residue-resolved Φ-value heatmaps from WSME-L. The region enclosed by the yellow box indicates the nbBAB motif. **(C)** Top, class-mean $p_{\mathrm{mt}}(X)$ versus Φ-value threshold $X$ for SSA, WSME-exact, and WSME-L. Bands indicate ±1 standard error of the mean. Bottom, representative examples of the fastest-folding nucleotide-binding motifs identified from each method, including flavodoxin, traced under all three methods (WSME-SSA: Rossmann, e1d4oA1; P-loop, e3bafA1; flavodoxin, e3olxA1. WSME-exact: Rossmann, e1d4oA1; P-loop, e5np9A1; flavodoxin, e3olxA1. WSME-L: Rossmann, e1d4oA1; P-loop, e2qg6A1; flavodoxin, e3olxA1).

### 2 WSME tutorial

This tutorial gives the physical and mathematical context needed to follow the WSME-based analyses used in the main text; parameter choices, hyperparameter-search protocols, percentile-rank statistics, and dataset specifications are described in the Methods and elsewhere in the SI. The three variants used in this work (SSA, WSME-exact, WSME-L) are presented in a single physicochemical convention. Throughout this work, we set inverse temperature $\beta = 1/(k_B T)$, attractive contact energies negative ($\epsilon_{u,v} \leq 0$).

#### 2.1 Native-centric statistical mechanics of protein folding

A folded single-domain protein is, to a good first approximation, a small thermodynamic system whose accessible states are dominated by configurations close to its native fold. The Gō hypothesis [12] takes this observation as a modeling axiom, according to which the folding mechanism is determined predominantly by the contact topology of the native state, and only contacts present in that native state contribute to the energy. The associated consistency principle states that the most stable structure of any local fragment is consistent with the native structure of the full-length protein, and is regarded as equivalent to the *principle of minimal frustration* [19]: in foldable proteins, the energetic frustration arising from non-native contacts is minimal.

The Wako–Saito–Muñoz–Eaton (WSME) family of models [13, 14, 15] is the simplest serious instantiation of this idea. Each residue $k$ carries an Ising-like binary variable $m_k \in \{0, 1\}$ that is either native-like ($m_k = 1$) or unfolded ($m_k = 0$). A microstate is a string of those binary variables; the energy is a structurally constrained sum of pairwise contact energies plus a per-residue entropic cost. At the level of a transform, the WSME model takes as input the contact energies of the native fold and outputs a free-energy landscape as a function of one or two coarse-grained order parameters (here we consider the fraction of native residues and the fraction of native contacts formed).

The key approximation that makes the WSME partition function tractable is the *contiguity ansatz*: a native contact between residues $i$ and $j$ contributes only when every residue between them is also folded. This forces folding to be initiated by local interactions and to spread to distal regions through the growth and docking of native segments [19].

Brute-force enumeration of the $2^D$ microstates is infeasible for $D \gtrsim 60$ ($2^{60} \approx 10^{18}$ states), so the WSME literature offers a hierarchy of computational strategies that all sum the same Hamiltonian but over different microstate subsets [19]: single, double and triple sequence approximations (SSA, DSA, TSA) restrict the number of simultaneously folded segments to $\leq 1, 2, 3$; the Bruscolini–Pelizzola transfer-matrix solution [16] sums the full $\{0, 1\}^D$ space exactly in polynomial time; and the linker (DSA/L and WSME-L) variants add cross-gap contacts between two folded segments separated by an unfolded loop, paid for by a polymer ring-closure entropy. In this work we use three variants of WSME of increasing mathematical and physical completeness: the SSA, the Bruscolini–Pelizzola exact solution (denoted WSME-exact), and the linker-augmented model of Ooka & Arai [6] (denoted WSME-L).

#### 2.2 Common machinery shared by all three variants

##### 2.2.1 Microstates and the contiguity ansatz

Let the protein have $D$ residues. A microstate is a binary string

$$m = (m_1, m_2, \ldots, m_D), \qquad m_i \in \{0, 1\}, \tag{1}$$

where $m_i = 1$ denotes that residue $i$ is in its native conformation. The microstate space is $\{0, 1\}^D$, of size $2^D$.

The *contiguity rule* is that the contact $(i, j)$ contributes $\epsilon_{i,j}$ to the microstate energy if and only if $m_i = m_{i+1} = \cdots = m_j = 1$. A maximal stretch of consecutive folded residues is a *contig* $c = [i, j]$ with length $K = j - i + 1$. Every microstate decomposes uniquely into a (possibly empty) collection of non-overlapping contigs separated by at least one unfolded residue, and the energy is a sum over those contigs.

##### 2.2.2 The Boltzmann weight of a single contig

For a contig $c = [i, j]$, the standard folding free energy and Boltzmann weight are

$$\Delta G(i, j) = \sum_{i \le u \le v \le j} \epsilon_{u,v} \; - \; T \sum_{k=i}^{j} \Delta S_k, \tag{2}$$

$$w(i, j) = \exp\!\big(-\beta\, \Delta G(i, j)\big), \quad \beta = \tfrac{1}{k_B T}. \tag{3}$$

The first sum captures all pairwise contact energies entirely contained inside the contig (favourable, $\le 0$); the second is the entropic cost of ordering the $K = j - i + 1$ residues of the contig (unfavourable: $\Delta S_k < 0 \Rightarrow -T\,\Delta S_k > 0$). With $k_B = 1.987 \times 10^{-3}\,\mathrm{kcal\,mol^{-1}\,K^{-1}}$ and uniform $\Delta S_k \equiv \Delta S = -3.5\,k_B$ in this work, equation (3) reduces to

$$w(i, j) = \exp\!\Big(-\beta \sum_{i \le u \le v \le j} \epsilon_{u,v} \; + \; \tfrac{K\,\Delta S}{k_B}\Big). \tag{4}$$

##### 2.2.3 Partition function, order parameter, and free-energy landscape

The full partition function is the sum of Boltzmann weights over the allowed microstate set $\mathcal{M}$,

$$Z = \sum_{m \in \mathcal{M}} \; \prod_{\text{contigs } c \in m} w(c), \tag{5}$$

where $\mathcal{M}$ is "at most one contig" for the SSA, "any non-overlapping collection of contigs" for WSME-exact, or that same set augmented by virtual-linker configurations for WSME-L. Microstates are bucketed by an integer-valued order parameter $X(m)$ to obtain the one-dimensional thermodynamic picture

$$Z(X) = \sum_{m:\, X(m) = X} \; \prod_{c} w(c), \qquad G(X) = -k_B T \ln Z(X). \tag{6}$$

The choice of $X$ most cleanly distinguishes the three variants:

- SSA uses $K$, the length of the (single) contig.
- WSME-exact uses $q = \sum_i m_i$, the total number of folded residues.

- WSME-L uses either $q$ in single-domain mode, or a pair $(q_1, q_2)$ of folded-residue counts in two user-defined residue blocks.

The unfolded reference $m = (0, \ldots, 0)$ contributes a Boltzmann weight of 1 in all variants. A stably folded system has a low-free-energy unfolded basin near the minimum of $X$, a low-free-energy folded basin near the maximum of $X$, and one or more barriers in between.

#### 2.3 Variant 1: SSA, the single-sequence approximation

##### 2.3.1 Restriction and partition function

The SSA restricts $\mathcal{M}$ to configurations containing *at most one* contiguous folded stretch [15]. There is one unfolded microstate plus $D(D+1)/2$ single-contig microstates indexed by $(i, j)$ with $1 \leq i \leq j \leq D$. With the unified Boltzmann weight of equation (3), the partition function is the explicit sum

$$Z_{\text{SSA}} = 1 + \sum_{1 \leq i \leq j \leq D} w(i, j), \tag{7}$$

evaluable in $O(D^2)$ time, where the leading 1 is the weight of the fully unfolded reference. Bucketed by contig length $K$,

$$Z(K) = \sum_{i=1}^{D-K+1} w(i,\, i + K - 1), \quad K = 1, \ldots, D, \quad Z(0) = 1, \tag{8}$$

so the one-dimensional free-energy profile is $G(K) = -k_B T \ln Z(K)$. More directly, every cell $(i, j)$ of the triangular lattice $\{(i, j) : 1 \leq i \leq j \leq D\}$ carries a Boltzmann probability

$$p(i, j) = \frac{w(i, j)}{Z_{\text{SSA}}}, \tag{9}$$

the equilibrium occupancy of the corresponding single-contig microstate. The SSA free-energy landscape used below is, up to an additive constant, $G(i, j) = -k_B T \ln p(i, j)$.

##### 2.3.2 Setting the entropy parameter analytically

The SSA has a single free thermodynamic parameter, the uniform per-residue entropy $\Delta S \equiv \Delta S_k$. It is fixed analytically by demanding that the folding free energy of the fully folded contig $\Delta G(1, D)$ match a target value $\Delta G_{\text{fold}}$. From equation (2),

$$\Delta G(1, D) = \sum_{1 \le u \le v \le D} \epsilon_{u,v} - TD\,\Delta S = H_{\text{tot}} - TD\,\Delta S \stackrel{!}{=} \Delta G_{\text{fold}}, \tag{10}$$

where $H_{\text{tot}} \le 0$ is the total contact enthalpy summed over all surviving contacts. Solving,

$$\Delta S = \frac{H_{\text{tot}} - \Delta G_{\text{fold}}}{TD}. \tag{11}$$

Where experimental folding free energies are available (e.g. from differential scanning calorimetry or chemical denaturation), they are used directly; in their absence we set $\Delta G_{\text{fold}} = -10\,k_B T$ as the default for stably folded proteins (Methods). This completely specifies the SSA without any grid search. For a stably folded protein, $|H_{\text{tot}}| > |\Delta G_{\text{fold}}|$, so the numerator of equation (11) is negative and $\Delta S < 0$ as required.

##### 2.3.3 The SSA free-energy landscape as a triangular lattice and folding paths

Each contig $(i, j)$ corresponds to a single cell of the triangular lattice $\{(i, j) : 1 \le i \le j \le D\}$, with contig length $K = j - i + 1$ increasing from the diagonal $i = j$ ($K = 1$, single residues) to the apex $(1, D)$ ($K = D$, fully folded). Plotting $G(i, j)$ on this triangle yields a two-dimensional surface whose topology is amenable to direct enumeration of kinetically meaningful folding routes.

A folding path on this landscape is a sequence of contigs

$$(i_0, j_0) \to (i_1, j_1) \to \cdots \to (i_T, j_T) = (1, D), \tag{12}$$

starting from a single seeded residue on the diagonal ($K_0 = 1$, $i_0 = j_0$) and ending at the apex ($K_T = D$, fully folded), such that the contig length $K_t = j_t - i_t + 1$ is strictly monotone in $t$. Each step extends the current contig by one residue at the N-terminus, the C-terminus, or both, encoding the progressive growth of a single folded segment. The maximum free energy traversed along a path is its barrier; ranking paths by ascending barrier and intersecting them with structural windows of interest is the geometric content of the I-TN analysis used in the main text (full procedure: companion file `itn_pseudocode.tex`).

#### 2.4 Variant 2: WSME-exact, the Bruscolini–Pelizzola transfer-matrix solution

##### 2.4.1 What changes relative to the SSA

The exact WSME partition function sums over the full microstate space $\{0, 1\}^D$, equivalently over all combinations of non-overlapping contigs [16]. Every contiguous folded stretch contributes its own intra-contig $\Delta G(i, j)$ via the unified weight $w(i, j)$ of equation (3), and every folded residue still pays its entropic cost. The contiguity ansatz is preserved at the level of each individual contig: a contact between residues $u$ and $v$ contributes only when $u$ and $v$ lie in the *same* contig. Two folded contigs separated by even a single unfolded residue therefore cannot share a contact, even if their constituent residues are spatially close in the native structure — the central restriction motivating the WSME-L extension of Section 2.5.

The order parameter changes accordingly. The SSA could use $K$ because there was only ever one contig; with multi-contig microstates, the natural global progress variable is the total number of folded residues across all contigs,

$$q(m) = \sum_{i=1}^{D} m_i \in \{0, 1, \ldots, D\}. \tag{13}$$

The output is a one-dimensional free-energy landscape $G(q) = -k_B T \ln Z(q)$ for a single domain. For two-block analyses (used as the basis of the WSME-L 2D landscape in Section 2.5) the $D$ residues are split into two blocks of sizes $n_1$ and $n_2$ with $n_1 + n_2 = D$, the folded residues in each block are counted separately, and the result reported as $G(q_1, q_2)$.

##### 2.4.2 The transfer-matrix recursion

Naively summing over $2^D$ microstates is infeasible. Bruscolini & Pelizzola showed that the contiguity structure allows the partition function to be built by a one-pass dynamic-programming recursion that enumerates microstates by the position and length of the *trailing* contig. Define

$$Z_n(q) = \sum_{\substack{m\in\{0,1\}^n \\ \sum_i m_i = q}} \prod_{\text{contigs } c\subset m} w(c), \tag{14}$$

the WSME partition function restricted to the first $n$ residues at total folded count $q$, with $w(c)$ the unified Boltzmann weight of equation (3). Then $Z_n$ is built from smaller subproblems by considering whether residue $n$ is unfolded or belongs to a trailing contig of length $K$:

$$Z_n(q) = Z_{n-1}(q) + \sum_{K=1}^{n} Z_{n-K-1}(q-K)\, w(n-K+1,\, n), \tag{15}$$

with the conventions $Z_{-1}(\cdot) = Z_0(0) = 1$ and $Z_0(q > 0) = 0$. The factor $Z_{n-K-1}$ encodes the requirement that residue $n-K$ (immediately before the trailing contig) is unfolded. After processing all $D$ residues, the global landscape is read off as $Z(q) = Z_D(q)$. The recursion is $O(D^3)$ in time and $O(D^2)$ in memory.

##### 2.4.3 What is gained and what is still missing

Relative to the SSA, WSME-exact resolves the contributions of multi-contig microstates exactly within the WSME ansatz. This admits genuine cooperativity in the folding dynamics, with intermediates supported by simultaneous growth of two or more fragments, and yields well-defined 1D and 2D global progress coordinates. It is also the appropriate setting for the standard theoretical $\phi$-value analysis of Section 2.7, because the resulting landscape $G(q)$ is the correct one to perturb.

The contiguity ansatz nevertheless still forbids any native contact whose endpoints lie in distinct contigs: a long-range tertiary interaction across an unfolded loop or a terminal $\beta$-strand pairing is silently set to zero in the energy. This blind spot to long-range tertiary contacts is the central motivation for the WSME-L extension recently developed by Ooka & Arai [6], which we describe next.

#### 2.5 Variant 3: WSME-L, the linker extension

##### 2.5.1 The cross-gap problem and the virtual-linker idea

In the vanilla model, two contigs separated by an unfolded segment are energetically decoupled: any native contact whose endpoints lie in different contigs is excluded by the contiguity rule. WSME-L [6] relaxes this by allowing a *virtual linker* to be placed across any sufficiently attractive native pair $(u, v)$ with

$$\epsilon_{u,v} < c_{\text{cut}} = -0.6\, k_B T, \tag{16}$$

the linker contact cutoff fixed at this value throughout the work. The virtual linker recognises that the unfolded segment between $u$ and $v$ may still adopt a conformation in which the contact $(u, v)$ is realised, with the rest of that segment forming a closed loop. Loop closure is enthalpically favourable (the contact energy $\epsilon_{u,v} \le 0$ is gained) but entropically costly (the loop must close).

##### 2.5.2 Ring-closure entropy

The entropic cost of closing a loop of $n_{\text{gap}}$ unfolded residues whose endpoints reach a fixed $C_\alpha$–$C_\alpha$ distance $d_{u,v}$ is the classical *ring-closure entropy* of polymer physics [18]. For a Gaussian chain the corresponding probability density of finding the chain ends at separation $d_{u,v}$ scales as

$$P(d_{u,v} \mid n_{\text{gap}}) \propto n_{\text{gap}}^{-3/2}\, \exp\!\left(-\frac{3\, d_{u,v}^2}{2\, n_{\text{gap}}\, b^2}\right), \tag{17}$$

with $b$ the effective Kuhn length, so the ring entropy carries a $-(3/2)\, k_B \ln n_{\text{gap}}$ contribution from chain length plus a contribution that depends on $d_{u,v}$ relative to the typical end-to-end distance of an unfolded chain of length $n_{\text{gap}}$. The exact functional form used here is that of [6], adopted without modification. This ring entropy is a *purely geometric* quantity: it depends only on $n_{\text{gap}}$ and $d_{u,v}$, and not on the sequence identity of the loop residues or on any energetic perturbation. Consequently, WSME-L uniquely requires the $C_\alpha$–$C_\alpha$ distance matrix $d_{i,j}$ as input; SSA and WSME-exact do not.

Denote the resulting ring-entropy contribution for a virtual linker at $(u, v)$ in a microstate of folded count $q$ as $\Delta S_{\text{ring}}^{(u,v)}(q) < 0$ (a *loss* of entropy, since the unfolded loop has fewer accessible conformations once its endpoints are pinned). WSME-L introduces an entropy-scaling hyperparameter $\xi$ that multiplies $\Delta S_{\text{ring}}^{(u,v)}(q)$ globally; $\xi$ is jointly tuned with $\epsilon$ by a 2D grid search (Methods).

###### 2.5.3 The augmented partition function

The WSME-L partition function adds, on top of the vanilla $Z(q)$, a contribution from each eligible linker that replaces the vanilla weights for the relevant fragment pair by linker-specific weights and applies the ring-entropy correction:

$$Z_L(q) = Z(q) + \sum_{(u,v):\,\epsilon_{u,v}<c_{\text{cut}}} \left[\, Z^{(u,v)}(q) - Z(q) \,\right] \exp\!\left(\frac{\Delta S_{\text{ring}}^{(u,v)}(q)}{k_B}\right). \tag{18}$$

Here $Z^{(u,v)}(q)$ is calculated using the WSME-exact recursion of equation (15), but the unified fragment weights $w(i,j)$ are locally augmented to include the cross-gap contact energy $\epsilon_{u,v}$ that the linker now allows on either side of the gap; the redundant direct contacts in the immediate neighbourhood of $(u,v)$ are removed to avoid double-counting. The structure of the recursion is unchanged — only the fragment weights that feed it differ. Because $\Delta S_{\text{ring}}^{(u,v)}(q) < 0$, the prefactor $\exp(\Delta S_{\text{ring}}^{(u,v)}/k_B) < 1$ correctly suppresses linker configurations by their entropic cost. Subtracting $Z(q)$ inside the bracket makes the linker contribution incremental, so the absence of any linker (or an infinitely costly loop) recovers $Z_L(q) = Z(q)$.

###### 2.5.4 Two-dimensional landscape and 1D folding coordinate

To resolve which sub-structure of a domain folds first, WSME-L is operated in a two-block mode: the $D$ residues are partitioned into blocks of sizes $n_1$ and $n_2$ (the choice of split is documented per protein in the Supplementary Methods), the partition function is bucketed by the pair of folded-residue counts $(q_1, q_2)$, and the output is the two-dimensional free-energy landscape

$$G(q_1, q_2) = -k_B T \ln Z_L(q_1, q_2). \tag{19}$$

The block split is purely a reaction-coordinate choice: virtual linkers are still allowed to connect any eligible $(u,v)$ pair, whether within or across blocks. The decomposition controls how the same partition function is *projected* onto two axes, not which microstates are summed.

To compare WSME-L results to the inherently one-dimensional outputs of the SSA and WSME-exact, the 2D landscape is reduced to a 1D folding coordinate by a *monotone minimax path* from the unfolded corner $(0,0)$ to the native corner $(n_1, n_2)$. Among all paths in the lattice that are non-decreasing in both $q_1$ and $q_2$ at every step, the monotone minimax path minimises the maximum free-energy barrier traversed. Once the path is chosen, an arc-length parameterisation along it gives the 1D folding coordinate along which residue-resolved $\phi$-values are reported. Where multiple paths are nearly degenerate in their maximum barrier, this is reported explicitly because the choice can reorder the relative timing of secondary-structure consolidation.

#### 2.6 Limitations of the WSME family

The convenience of the WSME ansatz is bought at the price of several structural blind spots. We collect the limitations relevant to this work in one place; each implies a class of folding phenomena to which the corresponding WSME variant is, by construction, insensitive.

**Native-centric (Gō) bias.** All WSME variants assume that only native contacts contribute to the energy, so they are fundamentally blind to energetic frustration and to non-native interactions that frequently create transient kinetic traps. Proteins for which folding is dominated by misregistered intermediates or by deep non-native traps fall outside the model's domain of validity.

**Dynamic topological shifts.** The model has no intrinsic mechanism for slow post-translational topology changes such as proline *trans*-to-*cis* isomerisation, disulfide rearrangement, or domain-swap interconversion. Capturing such phenomena requires manual surgery on the rate matrices [19], not a change of the partition function.

**SSA size limitation.** Because the SSA strictly prohibits simultaneous multi-nucleation events, it cannot represent the cooperative co-assembly of two or more independent fragments and so does not scale to large globular proteins whose folding intrinsically requires multiple coexisting nuclei.

**WSME-exact topological rigidity.** Even with the full $\{0,1\}^D$ microstate space, the contiguity ansatz still demands that every residue between two contacting partners be folded. WSME-exact therefore assigns zero stabilisation energy to native tertiary contacts that bridge a disordered loop, and fails on discontinuous domains.

**WSME-L computational overhead.** Evaluating the augmented ensemble partition function requires iterating the WSME-exact transfer-matrix recursion once per eligible linker. With $O(D^2)$ candidate linkers and an $O(D^3)$ recursion per linker (Section 2.4), the full WSME-L landscape is $O(D^5)$, and residue-resolved $\phi$-values add a factor of $D$ on top. A joint two-dimensional grid search over the energy and entropy hyperparameters $(\epsilon, \xi)$ adds another $O(N_\epsilon N_\xi)$ multiplicative cost, creating a substantial computational footprint for large proteins.

**Ensemble-view limitation of WSME-exact and WSME-L (1D mode).** The order parameter $q = \sum_i m_i$ is a non-injective map from the $2^D$-microstate space onto $\{0, \ldots, D\}$. Properties that are constant on level sets of $q$ (total contact-energy budget, gross folded mass) survive the projection; everything that distinguishes microstates with the same $q$ — *which* residues are folded, *where* the nucleation sites are, *how many* simultaneous nuclei coexist — is averaged out. Two competing nucleation routes at opposite termini sharing the same intermediate $q$ are bucketed into the same $G(q)$ bin as a single big contig of identical mass, and whether $G(q)$ at that $q$ is dominated by a single-contig microstate or by an ensemble of two-contig microstates is invisible at the 1D level. Hypotheses about competition or cooperation between nucleation sites are therefore precluded by the very choice of $q$ as the reaction coordinate. Phi-value analysis (Section 2.7) restores per-residue timing but not pairwise cooperativity between nucleation sites; the WSME-L 2D landscape $G(q_1, q_2)$ recovers competition *between* blocks but coalesces competition *within* a block; only the SSA's full $G(i, j)$ retains the microstate-level information — which is one reason the I-TN analysis (Section 2.3.3) operates on the SSA lattice.

#### 2.7 Reading residue-level structure: theoretical $\phi$-value analysis

The order parameters $K$ (SSA) and $q$ (WSME-exact, WSME-L) are global progress variables: they say how much of the protein is folded but not *which* residues. Theoretical $\phi$-value analysis is a per-residue read-out that complements them and makes the WSME family directly comparable to experimental $\phi$-value measurements. The variant-independent recipe below follows [17, 6].

##### 2.7.1 The in-silico mutation perturbation

For each residue $l \in \{1, \ldots, D\}$, a destabilising mutation is simulated by weakening every attractive native contact involving $l$ by a small fixed factor $\delta = 0.1$. The perturbation matrix is

$$\eta_{i,j}^{(l)} = \begin{cases} \delta\,|\epsilon_{i,j}| & \text{if } (i = l \text{ or } j = l) \text{ and } \epsilon_{i,j} < 0, \\ 0 & \text{otherwise,} \end{cases} \tag{20}$$

so that $\eta_{i,j}^{(l)} \geq 0$ measures the magnitude of attractive interaction *removed*. The mutant fragment free energy becomes

$$\Delta G_{\text{mut}}^{(l)}(i,j) = \Delta G(i,j) + \eta_{\text{sum}}^{(l)}(i,j), \tag{21}$$

with $\eta_{\text{sum}}^{(l)}(i,j) = \sum_{i \leq u \leq v \leq j} \eta_{u,v}^{(l)}$, and the mutant Boltzmann weight is

$$w_{\text{mut}}^{(l)}(i,j) = w(i,j)\,\exp\!\big(-\beta\,\eta_{\text{sum}}^{(l)}(i,j)\big) < w(i,j), \tag{22}$$

correctly destabilising every fragment containing residue $l$. Computationally, $\eta_{\text{sum}}^{(l)}(i,j)$ is evaluated in $O(1)$ per query by precomputing a 2D prefix-sum table of $\eta^{(l)}$ in $O(D^2)$, so the mutant landscape is obtained by re-running each variant's partition-function machinery on the reweighted fragment weights.

Three points to flag. First, only attractive contacts ($\epsilon_{i,j} < 0$) are perturbed, matching the convention of [6], Eq. 26. Second, the perturbation acts on *all* attractive contacts touching residue $l$, not on a single side-chain interaction; the resulting $\phi_l$ is therefore a coarse-grained folding-environment measure for residue $l$, not the literal predicted $\phi$ of a specific point mutation. Third, in WSME-L the geometric ring entropy $\Delta S_{\text{ring}}^{(u,v)}$ is *not* perturbed: it depends only on $d_{u,v}$ and the gap length, both unchanged by an energetic mutation.

##### 2.7.2 The $\phi$ formula and threshold-crossing summaries

Let $G_{\text{ref}}(X)$ denote the reference (wild-type) landscape and $G^{(l)}_{\text{mut}}(X)$ the landscape recomputed with the residue-$l$ perturbation, both at the same order-parameter value $X$. The theoretical $\phi$-value at residue $l$ and reaction-coordinate position $X$ is [17, 6]

$$\phi_l(X) = \frac{\left[G^{(l)}_{\text{mut}}(X) - G^{(l)}_{\text{mut}}(U)\right] - \left[G_{\text{ref}}(X) - G_{\text{ref}}(U)\right]}{\left[G^{(l)}_{\text{mut}}(N) - G^{(l)}_{\text{mut}}(U)\right] - \left[G_{\text{ref}}(N) - G_{\text{ref}}(U)\right]}, \tag{23}$$

where $U$ and $N$ are the unfolded and native endpoints of the landscape ($X = 0$ and $X = D$ for SSA and WSME-exact; $(0, 0)$ and $(n_1, n_2)$ for the WSME-L 2D landscape, evaluated along the monotone minimax path). Numerator and denominator are clipped together so that $\phi_l \in [0, 1]$; by construction $\phi_l(U) = 0$ and $\phi_l(N) = 1$. The interpretation is the standard one: $\phi_l(X) \approx 0$ means residue $l$ is unfolded-like at folding stage $X$, while $\phi_l(X) \approx 1$ means residue $l$ has already developed its native energetic environment.

To convert the resulting $D \times$ (landscape size) $\phi$ matrix into per-residue *timing* information we extract, for each residue $l$, the threshold-crossing time

$$t_X(l) = \min\left\{ X/X_{\max} \ : \ \phi_l(X) \geq X_{\text{thr}} \right\}, \quad t_X(l) \in [0, 1], \tag{24}$$

the normalised reaction coordinate at which $\phi_l(\cdot)$ first crosses a chosen threshold $X_{\text{thr}}$, with linear interpolation between adjacent points. The main text uses $t_{80}$ and $t_{99}$ (thresholds 0.80 and 0.99). A small $t_X(l)$ means residue $l$ achieves a native-like environment early in folding; aggregating $t_X$ over the residues of the catalytic $\beta\alpha\beta$ window gives the mean and maximum used for the percentile-rank summaries reported in the main text.

##### 2.7.3 Per-variant differences in the perturbation step

The $\phi$ formula and the perturbation matrix $\eta^{(l)}$ are common to all three variants. What differs is *which* internal quantities are reweighted before the partition function is recomputed:

- **SSA.** The perturbation is applied directly to the contig Boltzmann weights $w(i, j)$. For each candidate residue $l$, the prefix-sum table over $\eta^{(l)}$ is built in $O(D^2)$, each contig's $\eta^{(l)}_{\text{sum}}$ is queried in $O(1)$, and the mutant stage-decomposed $Z_{\text{mut}}(K)$ is obtained in $O(D^2)$; iterating over all $D$ residues gives $O(D^3)$ overall.

- **WSME-exact.** The perturbation modifies the fragment Boltzmann weights $w(i, j)$ feeding the transfer-matrix recursion of equation (15). The recursion itself is unchanged; it is re-run for each residue with reweighted fragment weights. Total cost is $O(D^4)$, parallelising trivially across residues.
- **WSME-L.** Three quantities are reweighted: (i) the fragment Boltzmann weights, exactly as in WSME-exact; (ii) the cross-region linker weight matrices, requiring a rectangular prefix-sum query across cross-region contacts touching the mutated residue; and (iii) the contact energies in the immediate neighbourhood of each virtual-linker pair $(u, v)$, namely $(u, v)$ itself and the four adjacent positions $(u \pm 1, v)$ and $(u, v \pm 1)$. The full WSME-L recursion (vanilla recursion plus per-linker recursion) is then re-run on the reweighted inputs. As above, $\Delta S^{(u,v)}_{\text{ring}}$ is not perturbed.

In all three variants, the mutant landscape is then plugged into equation (23) together with the reference landscape, and the threshold-crossing extraction of equation (24) is performed identically.

#### 2.8 Comparative summary

Going SSA $\to$ WSME-exact adds multi-fragment cooperativity and intermediates supported by simultaneous growth of two or more contigs, replacing the single-contig progress variable $K$ by the global folded-residue count $q$ at the price of an $O(D^3)$ recursion. Going WSME-exact $\to$ WSME-L adds energetic coupling between folded fragments separated by an unfolded gap, paid for by a Gaussian-chain ring-closure entropy that uses the native $C_\alpha$–$C_\alpha$ distances and brings $D$ as a second required input. Theoretical $\phi$-value analysis is layered on top of any of the three landscapes by perturbing only the attractive contacts touching a target residue and re-running the variant's partition-function machinery. The SSA's microstate-level $G(i, j)$ retains all spatial information at the cost of forbidding multi-nucleation; WSME-exact and WSME-L recover multi-nucleation at the cost of projecting onto $q$ (or $(q_1, q_2)$ for WSME-L's 2D mode); and only the WSME-L 2D landscape together with $\phi$-value analysis simultaneously captures cross-gap energetic coupling and per-residue timing.
